# High-throughput biochemical phenotyping of SHP2 variants reveals the molecular basis of diseases and allosteric drug inhibition

**DOI:** 10.64898/2026.03.30.715055

**Authors:** Albert A Lee, Daniel A Mokhtari, Emily D Egan, Stephen C Blacklow, Daniel Herschlag, Polly M Fordyce

**Affiliations:** Department of Genetics, Stanford University, Stanford, CA 94305; Department of Biochemistry, Stanford University, Stanford, CA 94305; Velocity Bio, San Carlos, CA 94070; Department of Biological Chemistry and Molecular Pharmacology, Blavatnik Institute, Harvard Medical School, Cambridge, MA 02115; Department of Cancer Biology, Dana Farber Cancer Institute, Boston, MA 02215; Broad Institute of Harvard and MIT, Cambridge, MA 02142; Department of Bioengineering, Stanford University, Stanford, CA 94305; Sarafan ChEM-H, Stanford University, Stanford, CA 94305; Chan Zuckerberg Biohub San Francisco, San Francisco, CA 94110

## Abstract

Interpreting clinical and functional consequences of genetic variants remains challenging due to limited quantitative biochemical data at scale. We applied high-throughput microfluidic enzyme kinetics to profile 190 clinical variants of SHP2, a protein tyrosine phosphatase linked to developmental disorders and cancers. Through >300,000 reaction progress curves, we derived kinetic and thermodynamic parameters quantifying variant effects on catalysis, autoinhibition, stability, phosphopeptide binding, and drug responses. This multidimensional dataset reveals that dysregulated autoinhibition, rather than altered stability or catalysis, predominantly determines SHP2-associated pathogenesis. Thermodynamic modeling reveals that clinical-stage allosteric inhibitors preferentially stabilize a previously underappreciated, partially active conformation over the fully inactive state, leading to variant-dependent drug responses. Our high-throughput biochemical framework establishes a general approach to decipher the biochemical logic connecting protein variants to clinical outcomes.

## Introduction

Understanding how disease-associated mutations affect protein function, drive pathology, and can be therapeutically addressed is a central goal of biomedical science (*1–5*). Beyond its clinical importance, studying disease mutations has long fueled key discoveries of fundamental principles of molecular and biological function (*6*, *7*). While next-generation sequencing has unveiled a vast reservoir of human protein-coding variants (*8*, *9*), comprehensive biochemical and biophysical characterization of these variants at scale remains limited. As a result, most variants are only cataloged at the level of sequence and clinical observation, with their functional and clinical significance remaining largely unresolved (*4*, *10*). While mutational scanning assays can provide clinically relevant information by estimating variant impact on function using cellular fitness, these composite readouts typically do not directly measure the distinct physical parameters required to understand complex disease mechanisms. Here, we address this gap by systematically measuring biochemical and biophysical properties of 190 clinically documented variants of the disease-relevant protein tyrosine phosphatase (PTP) SHP2, providing a functional atlas that links mutations to disease phenotypes, therapeutic responses, and fundamental aspects of SHP2 biochemistry.

SHP2 is a ubiquitously expressed PTP encoded by the *PTPN11* gene and plays an essential role in regulating cellular processes, including metabolism, cell growth, differentiation, and immune response (*11–15*). Consistent with this central role, mutations in SHP2 are a major cause of multiple diseases (*16–21*). Germline SHP2 mutations cause autosomal dominant developmental disorders and are responsible for 50% of Noonan syndrome (NS) and 90% of Noonan syndrome with multiple lentigines (NSML) cases, disorders affecting 1 in 1000 live births (*17*, *22*, *23*). Somatic SHP2 mutations are associated with various types of cancer, including 35% of juvenile myelomonocytic leukemia (JMML) cases (*21*).

Structurally, SHP2 consists of two SH2 domains (nSH2 and cSH2), a catalytic PTP domain, and an unstructured C-terminal tail (**Fig. 1A**). SHP2 regulation is canonically described via a two-conformational-state model in which the protein interconverts between a closed, inactive conformation and an open, active conformation. In the autoinhibited conformation, the nSH2 domain sterically occludes the PTP active site and prevents catalysis (*24*, *25*); in the open conformation, the SH2 domains are rotated away such that the catalytic site is fully exposed (*26*). The open state can be further stabilized by the SH2 domains binding to phosphotyrosine (pTyr) motifs on activated receptors or scaffold proteins (*24*, *27–31*). The importance of SHP2 in proliferative signaling makes it a compelling target for cancer therapy (*15*, *32*, *33*), and at least 14 SHP2 inhibitors have progressed to clinical trials (*32–37*). Nearly all clinical-stage SHP2 inhibitors exploit SHP2’s native autoinhibitory mechanism. These compounds bind to a tunnel-like pocket at the interface of the SH2 and PTP domains, distal to the active site (*34*, *35*); binding is thought to stabilize the closed conformation, thereby potently and selectively inhibiting SHP2 relative to other PTPs.

**Figure 1.**
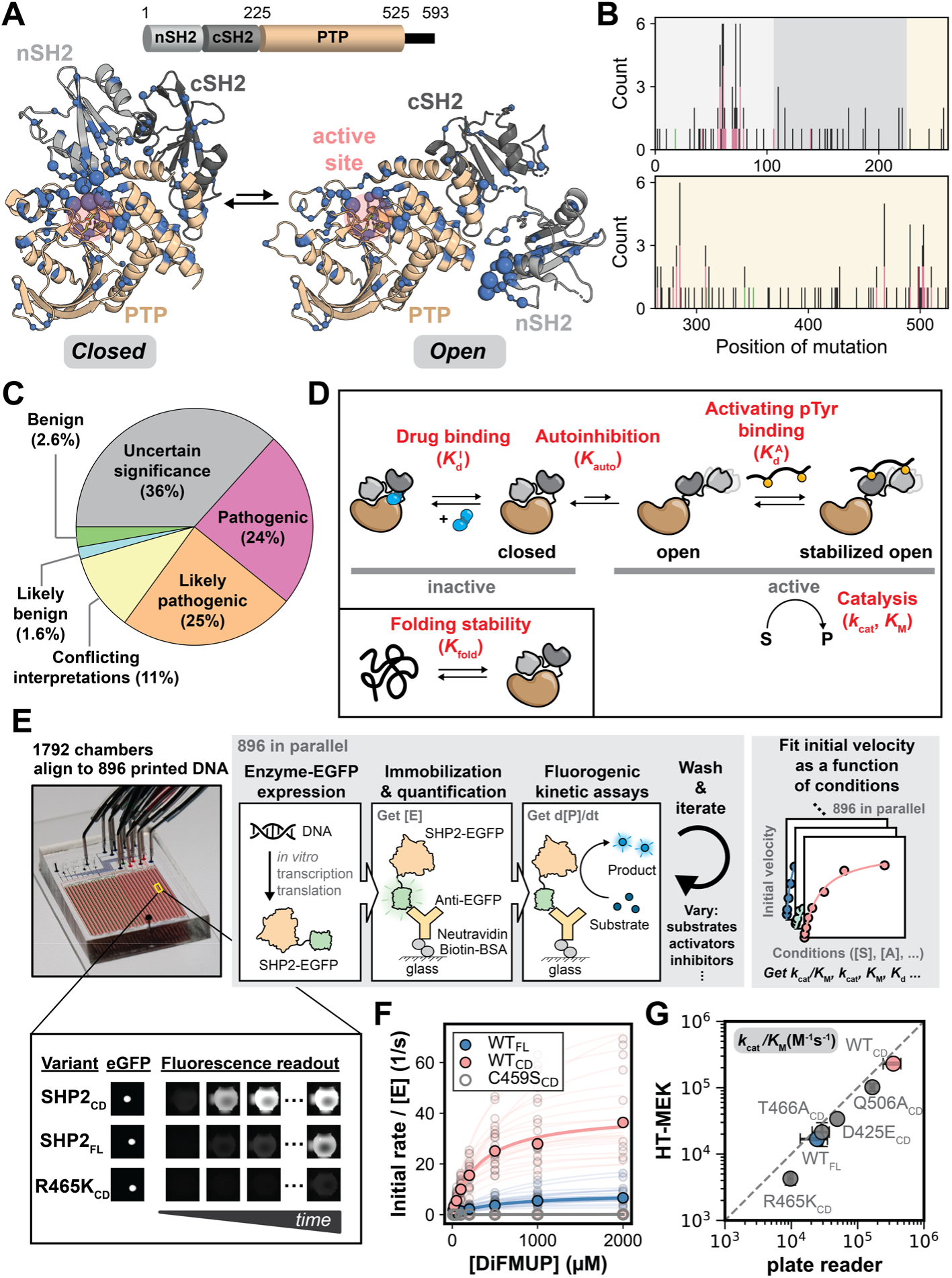
Overview of SHP2 and assaying variants in high throughput. **(A)** Crystal structure of SHP2 (PDB ID: 4DGP, left & 6CRF, right) highlighting positions mutated in the library in blue spheres; sphere size corresponds to the number of mutations at that position. **(B)** Distribution of unique single amino acid variants by position. Variants are color-coded by clinical classification: pathogenic (magenta), benign (green), and others (gray). **(C)** ClinVar classification of SHP2 variants in this study. **(D)** Schematic representation of key biochemical processes constituting SHP2 function. **(E)** Overview of HT-MEK microfluidic device and SHP2 on-chip expression, purification, and assay procedure. **(F)** Example Michaelis-Menten fits to initial rates. Opaque lines and points indicate the median (from 5–23 replicates), while semi-transparent lines and points represent individual replicates. **(G)** Comparison of median *K_cat_* /*K_M_* for DiFMUP hydrolysis measured with HT-MEK and plate reader.

Despite progress in understanding SHP2 pathology and function, over half of the >500 *PTPN11* missense variants in the ClinVar database are variants of uncertain significance (VUS), complicating genetic test interpretation and clinical management (*10*). Moreover, current data remain insufficient to define the biochemical basis of overlapping and divergent clinical phenotypes of SHP2 variants. For instance, NS and NSML share clinical features and cancer predisposition (*38–40*), yet NS mutations enhance SHP2 basal catalytic activity, while NSML mutations reduce it (*41–44*). Prior studies show that some NSML variants also reduce autoinhibition (*25*, *44*), a defect shared with NS variants, suggesting that disease phenotypes may depend on an interplay between catalytic and conformational effects. However, no systematic comparison has defined how catalytic and conformational effects combine to account for NS and NSML phenotypes. Similarly, it is unclear why both hyperactive and hypoactive SHP2 variants are associated with cancer (*15*, *33*, *45*), and why some mutations are cancer-specific while others overlap with developmental disorders (*46*, *47*). In-depth knowledge of the biochemical consequences of disease-causing variants at scale is needed to resolve these paradoxes.

Our understanding of SHP2 sensitivity towards clinical-stage inhibitors is also critically limited, as the inhibitor sensitivity of most SHP2 variants remains unknown. Moreover, while detailed studies of several cancer-associated mutants, including the canonical E76K, show that they shift the conformational ensemble toward the open state and confer resistance to the allosteric inhibitor SHP099 (*26*, *48*), it is unclear if this mechanism applies broadly across different variants and drug scaffolds. Addressing this limitation requires a systematic map of variant-specific drug sensitivity, coupled with quantitative biochemical analysis to resolve the mechanistic basis of these changes.

Here, we performed deep biochemical phenotyping on a library of 190 clinically documented SHP2 variants via HT-MEK, a microfluidic platform for high-throughput and quantitative biochemistry (*49*, *50*). We isolated and quantified distinct biochemical mechanisms by which mutations alter SHP2 function, including catalytic activity, stability, autoinhibition, and its response to physiological activators and clinical inhibitors, yielding ten kinetic and thermodynamic parameters that collectively define the biochemical phenotype of each variant. Analysis of this dataset revealed distinct biochemical signatures that predict pathogenicity and stratify variants by disease class, providing a biochemical basis that reconciles paradoxes in SHP2-associated pathologies. Profiling responses to three clinical-stage inhibitors uncovered a ∼100-fold range in sensitivity across variants and revealed an unexpected trend that contradicts the canonical two-conformational-state (closed/open) model, in which mild loss of autoinhibition leads to highest drug sensitivity. These findings led us to a three-conformational-state model of allosteric inhibition that accurately accounts for all observed measurements of SHP2 inhibition and suggests strategies to enhance therapeutic potency. This work provides an in-depth mechanistic framework for SHP2 pathology and therapeutic strategies, establishing a scalable approach for causally linking genotype to molecular function, disease, and therapeutic response that can be applied to many additional clinically relevant proteins.

## Results

### HT-MEK enables high-throughput expression, purification, and activity assays for SHP2 variants

To systematically characterize impacts of human allelic variation on SHP2 biochemical function, we generated a library of 189 missense variants documented in ClinVar within SHP2’s structured domains (residues 1-525, hereafter SHP2 or SHP2_FL_) (**Fig. 1A; Supplementary Table 1**) (*10*). These mutations are distributed throughout the linear sequence and across the structured domains of SHP2 (**Figs. 1A**&**B**). The library includes variants classified as pathogenic (46), benign (5, including WT), of uncertain significance (VUS; 69), or with conflicting/less certain annotations (70) (**Fig. 1C**).

Mutations of SHP2 can alter different aspects of biochemical function, including thermodynamic stability, intrinsic catalytic activity (activity of the isolated PTP domain), autoinhibition, regulation by activating pTyr peptide, or inhibition by drugs (**Fig. 1D**). To separately quantify the impacts on each of these properties for all variants with high accuracy and precision, we used High-Throughput Microfluidic Enzyme Kinetics (HT-MEK) (*49*), a microfluidic platform that enables recombinant expression, purification, and quantitative functional characterization of 1000s of enzymes in parallel (**Fig. 1E**). In HT-MEK, microfluidic devices containing 1798 reaction chambers are aligned with printed plasmid arrays. Introduction of cell-free expression reagents enables parallel expression of a unique C-terminally EGFP-tagged enzyme variant per chamber. The expressed proteins are then immobilized on device surfaces pre-patterned with anti-EGFP antibody, enabling purification and quantification. Integrated on-chip valves protect the immobilized enzymes during reagent exchange and permit reactions to be initiated and terminated with precise temporal control, allowing iterative assays for high-throughput measurement of thermodynamic and kinetic parameters (**Fig. 1E**).

To test if HT-MEK can accurately measure kinetic constants of SHP2-mediated catalysis, we quantified hydrolysis of the fluorogenic substrate 6,8-difluoro-4-methylumbelliferyl phosphate (DiFMUP) by a control library consisting of SHP2_FL_, a truncated construct containing only the catalytic domain (SHP2_CD_), four active site variants (D425E_CD_, R465K_CD_, Q506A_CD_, T466A_CD_), and a catalytically dead variant (C459S_CD_) (**Figs. S1**&**S2**). Measured activities for C459S_CD_ were indistinguishable from background activity (**Figs. 1F**&**S3**). HT-MEK-derived Michaelis-Menten parameters for these constructs were consistent (within ∼2-fold) with values obtained from traditional plate-based assays (**Figs. 1F**&**G**) and within the range of literature values (**Fig. S4**) (*51*, *52*).

### SHP2 variants in different structural regions substantially increase or decrease basal activity

Next, we applied HT-MEK to measure *k*_cat_, *K*_M_, and *k*_cat_/*K*_M_ for DiFMUP hydrolysis for 190 SHP2_FL_ allelic variants (**Fig. 2A**). Data were consistent across 10 devices (**Figs. S5**&**S6**), and we aggregated measurements across all replicates (median n = 19) to report median values for each variant (**Figs. 2B**, **S7**, **S8**).

**Figure 2.**
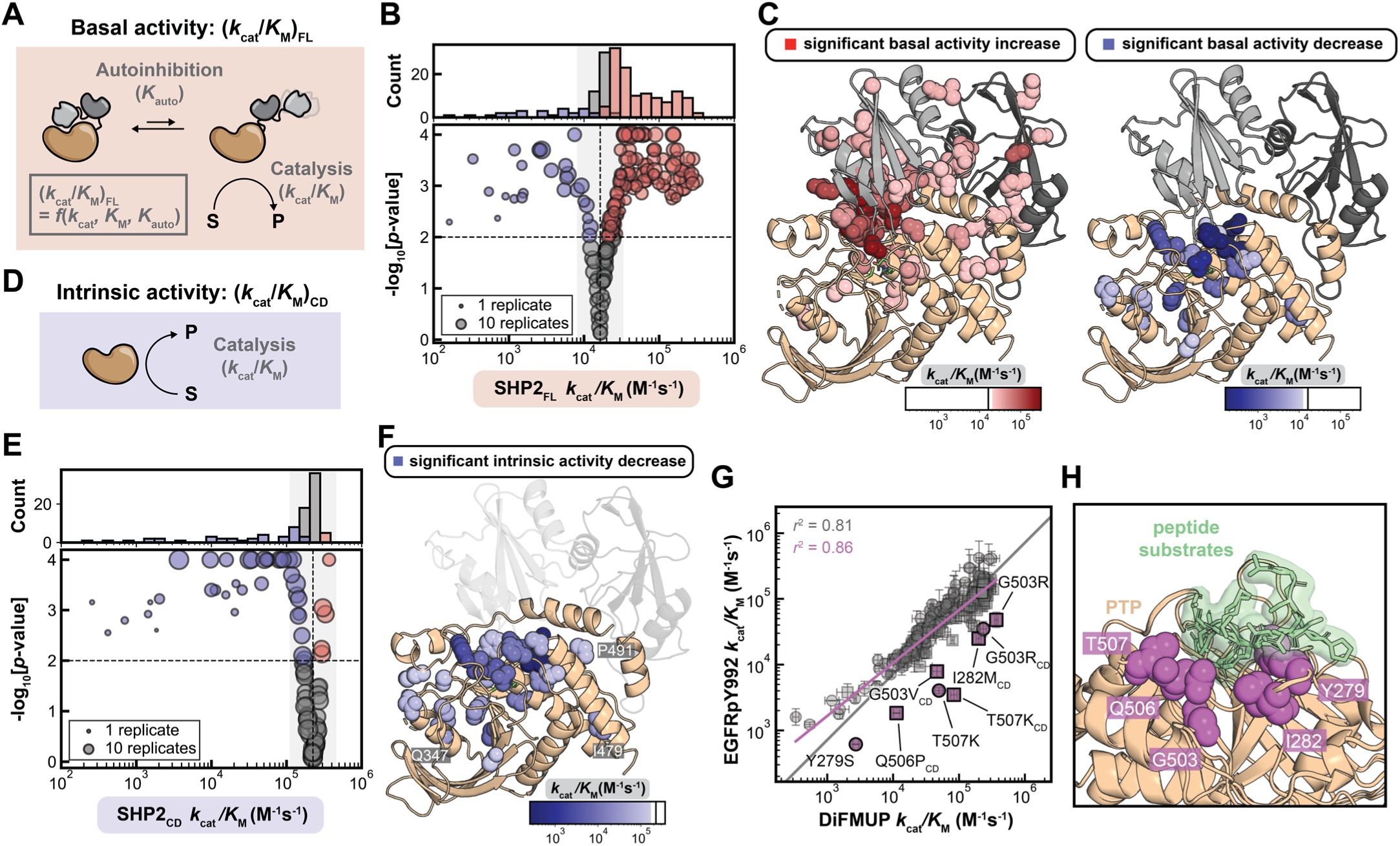
Widespread SHP2 variant activity reveals distinct structural hotspots for activity modulation. **(A)** Basal activity is defined as the *K_cat_* /*K_M_* of SHP2_FL_, which is determined by the autoinhibition equilibrium and the catalytic domain activity. **(B)** Median *K_cat_* /*K_M_* for DiFMUP hydrolysis across 190 SHP2_FL_ variants. Red: 120 significantly activating variants (*p* < 0.01). Blue: 26 significantly deactivating variants (*p* < 0.01). Gray: not significantly different from WT (*p* ≥ 0.01). Gray shaded area is within 2-fold from WT. **(C)** Structural mapping of mutations with significantly increased (left) or decreased (right) basal SHP2 activity. Color intensity of spheres reflects average variant activity at each position. The WT activity is marked by a black reference line on scale bar. **(D)** Intrinsic activity is defined as the *K_cat_* /*K_M_* of SHP2_CD_. **(E)** Median *K_cat_* /*K_M_* for DiFMUP hydrolysis across 95 SHP2_CD_ variants. Red: 5 significantly activating variants (*p* < 0.01). Blue: 36 significantly deactivating variants (*p* < 0.01). Gray: not significantly different from WT (*p* ≥ 0.01). **(F)** Structural mapping of mutations with significantly decreased intrinsic activity. Blue spheres indicate side chain positions and color intensity reflects average variant activity at each position. **(G)** Comparing mutational effects on hydrolysis of DiFMUP and EGFRpY992 peptide substrate. Grey line: log-log fit constrained to slope = 1 (*r*² = 0.81); purple line: unconstrained log-log linear regression (*r*² = 0.86, slope = 0.81). Variants deviating from the regression (residual |*z*-score| > 3) are colored in purple. **(H)** Structural location of the mutants (panel **G**, purple) with differentially altered activity against DiFMUP versus EGFRpY992 (PDB ID: 7PPL, 7PPM, 7PPN; overlayed).

Observed changes in *k*_cat_/*K*_M_ within the SHP2_FL_ construct (“basal activity”) ranged from a 30-fold increase to a 30-fold decrease in *k*_cat_/*K*_M_ (**Fig. 2B**). Impacts of different substitutions at the same position varied in magnitude but generally shifted *k*_cat_/*K*_M_ in the same direction (**Fig. S8**). Most variants (120/189) significantly increased activity, while 26 significantly decreased activity and 43 did not show a statistically significant change from WT (via bootstrap hypothesis testing) (**Fig. 2B**). The large number of replicates allowed us to confidently detect even subtle effects, including *k*_cat_/*K*_M_ decreases and increases of 26% and 27%, respectively (**Fig. 2B**). Mutations that increased the basal activity clustered around the interface between the PTP and the nSH2 domain (**Fig. 2C**), consistent with reduced autoinhibition. Mutations that reduced the basal activity were exclusively found in the PTP domain and concentrated around the active site, consistent with reduced intrinsic activity of the PTP domain itself (**Fig. 2C**), a model that we directly tested below.

### SHP2 catalytic domain variants predominantly reduce intrinsic activity

Changes in basal activity upon mutation could result from altered phosphate ester hydrolysis of the catalytic PTP domain (“intrinsic activity”) (**Fig. 2D**), altered amount of natively folded protein, or altered autoinhibitory regulation. To eliminate effects from changes in autoinhibition, we measured the impact of the 94 unique mutations within the isolated catalytic PTP domain (SHP2_CD_) on DiFMUP hydrolysis (*k*_cat_, *K*_M_, and *k*_cat_/*K*_M_) (**Figs. S9-S12**). In contrast to SHP2_FL_, variants in the isolated domain were predominantly inactivating. Thirty-six variants significantly reduced *k*_cat_/*K*_M_ (1.3- to >800-fold reductions), while none increased *k*_cat_/*K*_M_ more than two-fold (**Fig. 2E**). As expected, deactivating mutations clustered around the active site (**Fig. 2F**). However, three deactivating mutations were found >20 Å from the catalytic cysteine (Q347R, I479T, P491H). Notably, I479 and P491 reside on helices flanking a conserved loop known to be allosterically coupled to the active site conformation in the homolog PTP1B (**Fig. S13**) (*53*, *54*), highlighting HT-MEK’s ability to detect allosteric mutations and potential allosteric sites. The reduced activity of the remaining distal variant, Q347R, is instead fully accounted for by loss of natively folded protein (see below).

### Global thermodynamic destabilization of the SHP2 native state does not account for most observed activity changes

Enzymes must fold to be catalytically active, and it has been suggested that folding stability defects account for a majority of disease associated alleles (*55–57*). Measuring loss of enzymatic activity as a function of increasing chaotropic destabilizing agent (*e.g.* urea) can quantify impacts of mutations on folding stability (ΔG_fold_) for enzymes that undergo a reversible, two-state folding transition (**Fig. S14A**). While plate-based measurements of SHP2_CD_ DiFMUP hydrolysis as a function of urea show a fast, reversible two-state transition (**Fig. S14B**), HT-MEK measurements revealed a slow, irreversible loss of activity in addition to the reversible transition, likely due to interactions between the unfolded protein and the device surface (see **Supplementary Text 1**). Accounting for both processes allowed high-throughput quantification of SHP2_CD_ variants’ apparent native state stability (ΔG_native,app_) from HT-MEK data that agree with those measured in solution (estimated error < 10%) (**Figs. S14-S17**, **Supplementary Text 1**).

Of 95 SHP2_CD_ variants, 88 were sufficiently catalytically active to allow reliable quantification of urea-dependent loss of activity. Of these, 25/88 significantly reduced stability and none were significantly stabilized (*p* < 0.01) (**Fig. S14F**). Destabilizing mutations clustered on one face of the catalytic pocket, adjacent to the phosphotyrosine recognition loop and Q loop (**Fig. S14G**). Only five variants (G268C, C333S, Q347R, R501K, G503V) were less than 50% natively folded under our standard assay conditions (in the absence of urea) (**Fig. S14H**); these variants showed matching fold-decreases in natively-folded fraction and intrinsic activity (**Fig. S14I**), indicating that activity loss is fully accounted for by loss of native protein, including the observed distal effect of Q347R. Only three of the remaining 83 variants lose >20% of natively folded protein under urea-free assay conditions. Thus, the observed changes in intrinsic activity for most variants reflect direct effects on catalysis, not loss of natively folded protein (**Fig. S14I**).

Interpreting urea-dependent differences in activity for SHP2_FL_ is challenging, as multidomain SHP2_FL_ is not expected to follow simple two-state unfolding (**Supplementary Text 1**). Nevertheless, low concentrations of urea (1 M) primarily destabilize the PTP domain while leaving SH2 domains intact (*58*), making it possible to infer the stability of the PTP domain in SHP2_FL_ (**Fig. S18A**). We estimated the PTP domain within all SHP2_FL_ variants is >88% natively folded under normal assay conditions). In general, the PTP domain within SHP2_FL_ is more stable than when isolated, as expected from stabilization via interdomain contacts (**Figs. S18C**&**D**). Overall, the slight change in fraction native (a median effect of 1.7% in SHP2_FL_) cannot explain the much larger observed impacts on measured *k*_cat_/*K*_M_ values for SHP2_FL_ variants (**Fig. S18E**). Thus, these impacts are likely due to perturbations within the native state (*i.e.* altered autoinhibition).

### Comparative substrate profiling identifies mutations impacting peptide recognition

DiFMUP is a convenient probe of catalytic activity but it lacks the full set of interactions formed by physiological peptide substrates (*52*). To determine whether mutations perturb recognition of peptide substrates, we compared DiFMUP hydrolysis with dephosphorylation of a phosphopeptide derived from a known SHP2 target (EGFRpY992, DADEpYLIPQQG) (*44*, *59*). In this assay, we quantified hydrolysis by detecting inorganic phosphate release via fluorescence increase upon binding to a labeled phosphate-binding protein (*60*). The EGFRpY992 hydrolysis *k*_cat_/*K*_M_ values for both SHP2_FL_ and SHP2_CD_ constructs closely matched those for DiFMUP (*r*^2^ = 0.81, **Figs. 2G**, **S19-S22**), indicating that most variants do not alter peptide recognition and suggesting that DiFMUP is a reliable proxy for general catalytic activity. Mutations at positions 279, 282, 503, 506, and 507 significantly reduced *k*_cat_/*K*_M_ (residual |*z*-score| > 3) for EGFRpY992 relative to DiFMUP; all of these residues are located near the peptide-binding cleft, consistent with an effect on peptide substrate recognition (**Fig. 2H**) (*61*).

### Most variants increase basal activity by disrupting autoinhibition

To deconvolve and quantify impacts of mutations on autoinhibition, we compared DiFMUP *k*_cat_/*K*_M_ values for each variant within the SHP2_FL_ and SHP2_CD_ constructs. This comparison revealed four mechanistic categories (**Fig. 3A**): (1) WT-like variants with no change in either construct; (2) variants with reduced SHP2_CD_ and a proportional reduction in SHP2_FL_ activity, indicating decreased intrinsic activity; (3) variants with increased SHP2_FL_ activity but no change in SHP2_CD_ activity (including variants with mutations in the SH2 domains), consistent with impaired autoinhibition; and (4) variants with a greater decrease in SHP2_CD_ than SHP2_FL_ activity (or reduced SHP2_CD_ activity with mildly increased SHP2_FL_ basal activity), suggesting decreases in both intrinsic activity and autoinhibition.

**Figure 3.**
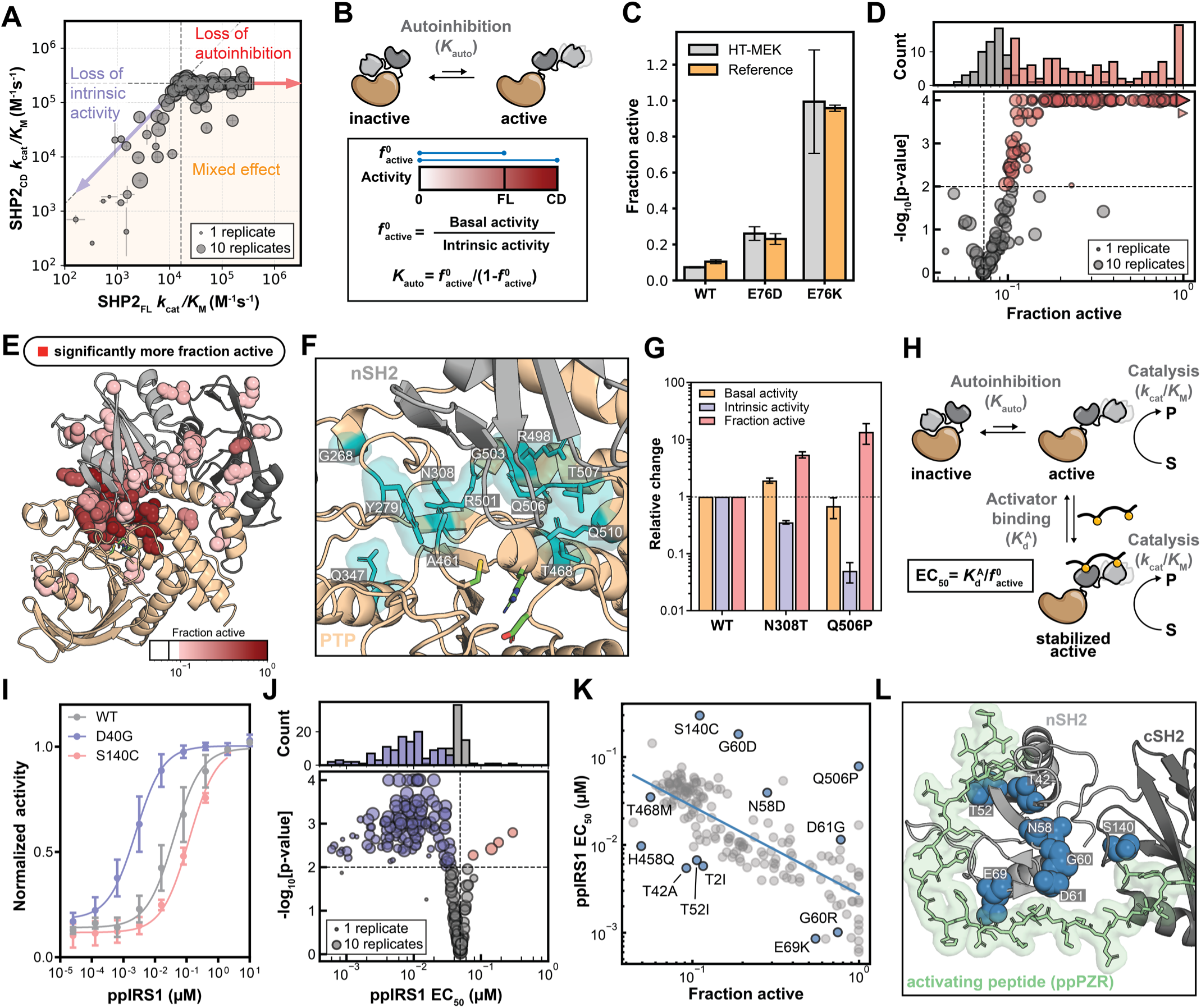
Autoinhibition equilibrium regulates SHP2 variant basal activity and ligand-dependent activation. **(A)** *K_cat_* /*K_M_* of DiFMUP hydrolysis in SHP2_CD_ versus SHP2_FL_ construct. WT values are marked by the intersecting dash lines. Error bar: bootstrap standard error. Square markers indicate mutations in the SH2 domains. **(B)** Calculation of fraction active. **(C)** HT-MEK measured fraction active compared to literature values (*48*). **(D)** Median fraction active of 185 SHP2 variants. Red: 121 significantly activating variants (*p* < 0.01). Gray: not significantly different from WT (*p* ≥ 0.01). **(E)** Positions with mutations causing significantly more fraction active shown as red spheres, with intensity reflecting the average effect at each position. **(F)** nSH2–PTP autoinhibitory interface. Positions with mutations that significantly reduce intrinsic activity and increase fraction active are highlighted in cyan. **(G)** Variants with significantly reduced intrinsic activity and increased fraction active, but <2-fold change in basal activity relative to WT. Data are presented as mean ± SEM. **(H)** Regulation of SHP2 by activators and parameters influencing EC_50_. **(I)** Example HT-MEK-measured dose-response curves for activating peptide ppIRS1. **(J)** Median EC_50_ of 188 SHP2 variants. Red: significantly activating variants (*p* < 0.01). Blue: significantly deactivating variants (*p* < 0.01). **(K)** ppIRS1 EC_50_ versus fraction active. Blue line indicates fitted model (**Eqn. 1**) assuming a constant affinity to activator 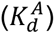. Highlighted variants (blue) significantly deviate by >3-fold from the global fit (*p* < 0.01) and have at least 3 replicates. **(L)** Mutations within the SH2 domains with significant deviation from the global EC_50_ versus fraction active fit from panel **K** plotted onto the structure (PDB ID: 9MQ5).

The extent of autoinhibition determines the fraction of SHP2 molecules in the open, active conformation. As SHP2_CD_ activity is approximately the same as the activity of the SHP2_FL_ open state (*26*, *48*), we quantified autoinhibition by calculating the ratio of *k*_cat_/*K*_M_ values for SHP2_CD_ and SHP2_FL_ for each mutation (**Fig. 3B**). By this measure, only 7.3% of WT SHP2 is in the open, active state (fraction active, 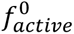) (95% CI: 6.5–8.1%). Two variants previously shown to lessen autoinhibition, E76D and E76K, are substantially more open, with 26% (95% CI: 22–30%) and 100% (95% CI: 70–100%) 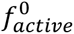, respectively (**Fig. 3C**). All three values are consistent with prior NMR-based estimates (*48*).

Overall, the calculated 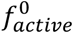 for SHP2 variants ranged from <5% to >99%, with most variants (121/185) significantly increasing 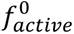 (**Fig. 3D**). These mutations predominantly clustered at the nSH2-PTP domain interface (**Fig. 3E**), consistent with the established role of this region in maintaining autoinhibition (*24*).

Mutations beyond this interface also significantly altered 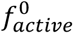, with most occurring within the SH2 domains (**Fig. 3E**) where they presumably perturb the SH2 domain conformational ensemble to weaken the autoinhibitory interaction (*24*, *28*, *30*). Since the PTP domain active site lies at the nSH2-PTP autoinhibitory interface, many mutations near the active site impair intrinsic catalysis while simultaneously relieving autoinhibition (“mixed effect” in **Fig. 3A**; **Fig. 3F**) such that they have little net impact on basal activity (**Figs. 3G**&**S23**). These mutants would be misclassified as “WT-like” when measuring basal catalysis alone, highlighting the value of acquiring multidimensional data.

### Conformational state and activator affinity determine the sensitivity of SHP2 variants to activation

SHP2 autoinhibition is regulated by the binding of pTyr-containing peptide (activator) to the tandem SH2 domains in the open, active conformation, typically described by a two-state conformational selection model (**Fig. 3H**) (*27*). To quantify impacts of mutations on activator sensitivity, we measured rates of DiFMUP hydrolysis as a function of increasing concentrations of ppIRS1, a synthetic bisphosphorylated peptide derived from IRS1 that binds both the nSH2 and cSH2 domains of SHP2 (**Fig. 3I**) (*32*). Fitting activation dose-response curves for 188 SHP2 variants (median replicate n = 9) yielded half-maximal effective peptide concentrations (EC_50_ values) that varied by >100-fold (**Figs. 3J**, **S24-S26**).

Based on a model in which one ppIRS1 binds per SHP2 molecule (**Fig. 3H**), EC_50_ (fitted with Hill slope of 1) is quantitatively determined by: (1) the intrinsic fraction active of SHP2 in the absence of activating ligand 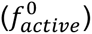, and (2) the affinity of the SH2 domains for the activator 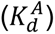 (**Eqn. 1**, see **Supplementary Text 2**):

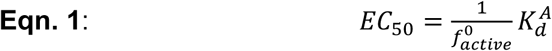

As most variants have mutations located far from the activator binding site, we expected that they would minimally impact affinity 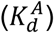 such that the EC_50_ would simply be inversely related to 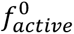. Consistent with this expectation, EC_50_ for most variants (176/188) were generally captured by their 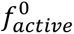 with a single globally-fitted 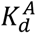 (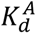 = 2.77 ± 0.16 *nm*, *r*^2^ = 0.63; **Fig. 3K**); a subset of variants (12/188) deviated significantly from this expectation more than 3-fold (*p* < 0.01). Consistent with altered ligand binding affinity, 9/12 of the deviating mutations appeared within the SH2 domains and this group included T42A, a variant known to increase ppIRS1 binding affinity (**Figs. 3L**&**S27**) (*62*). Our measurements of activity, stability, autoinhibition, and activator sensitivity are all consistent with the canonical understanding of SHP2 biochemistry and regulation. This suite of data established a foundation for decoding the biochemical basis of SHP2 pathogenesis, as explored below.

### Pathogenic variants converge on impaired autoinhibition and activator hypersensitivity

There is a basic expectation that variant-induced changes in biochemical functions ultimately drive disease phenotypes (*63–65*). To first test if measured biochemical parameters can distinguish pathogenic from benign SHP2 variants, we hierarchically clustered 179 variants based on five *z*-score-normalized biochemical features: basal activity, intrinsic activity, fraction active, stability, and activator EC_50_ (**Fig. 4A**). ClinVar-assigned pathogenic and benign variants (**Supplementary Table 1**) clustered separately, supporting the association of distinct biochemical profiles with pathogenicity. Notably, basal activity and stability—commonly emphasized in variant characterization and often assumed to drive disease (*55–57*)—were less effective predictors of SHP2 pathogenicity (**Fig. S28**). Instead, altered fraction active and activator EC_50_ were most strongly predictive of pathogenicity by leave-one-out cross-validation; indeed, using just these two features performed identically to using all five biochemical parameters (**Fig. S28**).

**Figure 4.**
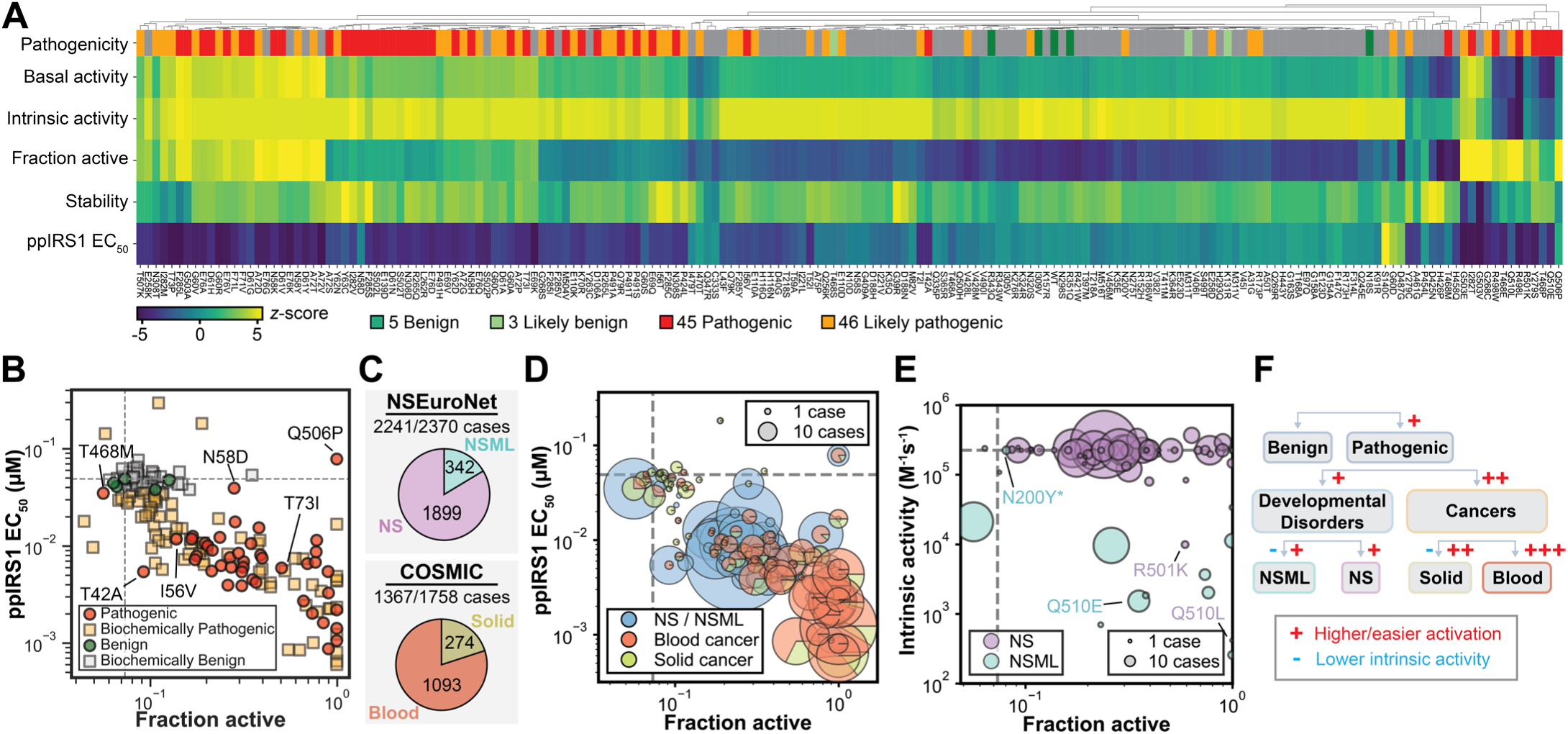
Biochemical signatures segregate SHP2 variants by disease class. **(A)** Hierarchical clustering of 179 SHP2 variants based on five measured biochemical parameters. **(B)** ppIRS1 EC₅₀ versus fraction active for 179 SHP2 variants. Variants with classification other than pathogenic or benign in ClinVar (squares) were classified based on *k*-nearest neighbor model. **(C)** NSEuroNet: 2241 (1899 NS; 342 NSML) of 2370 cases harbor variants in our library. COSMIC: 1367 (1093 blood; 274 solid) of 1758 cancer cases harbor variants in our library. **(D)** Comparing measured ppIRS1 EC_50_ versus fraction active values for variants reported in developmental disorder versus cancer cases; marker area is proportional to the number of case reports for each variant. **(E)** Intrinsic activity versus fraction active for variants reported as NS or NSML in the NSEuroNet database. Marker area is proportional to the number of case reports for each variant. *Confirmed misannotated variant. **(F)** Hierarchical classification of SHP2-related diseases and the biochemical distinction between subclasses.

Overall, 42/45 pathogenic variants have significantly higher 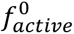 than benign variants, consistent with impaired autoinhibition predominantly driving pathogenicity. The remaining three (T42A, I56V, and T468M) were significantly more sensitive to ligand-induced activation (**Fig. 4B**). Thus, pathogenic SHP2 variants converge on a shared biochemical phenotype: dysregulated autoinhibition achieved via either destabilization of the autoinhibited state or enhanced sensitivity to activating ligands, regardless of their catalytic activity. This convergence suggests that the open conformation may drive pathogenesis primarily through its role as a signaling scaffold (*15*, *66–69*). Notably, AlphaMissense (*70*), a machine learning predictor for pathogenicity, identifies many variants with benign-level 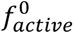 and activator EC_50_ as pathogenic (**Fig. S29**). This tendency to overpredict pathogenicity has also been observed in prior evaluations (*71–73*).

Next, using the two most informative measurements identified above (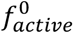 and activator EC_50_), we applied a *k*-nearest neighbor classifier to predict likely clinical impacts of 139 SHP2 variants with less certain clinical annotations. This approach predicted pathogenicity for 1/3 “likely benign” variants (33%), 19/60 VUS (32%), 15/19 “conflicting” variants (79%), 10/14 “likely pathogenic” variants (71%), and 32/33 “pathogenic/likely pathogenic” variants (97%) (**Figs. 4B**&**S30**). Thus, a substantial proportion of VUS may have pathogenic potential and warrant further clinical investigation. Since the start of our study in 2022, only one variant (T73I) has been upgraded to “pathogenic” in ClinVar (originally classified as conflicting interpretations of pathogenicity); our model’s prediction is consistent with this reclassification, highlighting the prognostic value of biochemical phenotyping (**Fig. 4B**).

### Quantitative differences in biochemical signatures differentiate disease subtypes

Next, to assess if different disease subtypes have distinct biochemical signatures, we compared measured biochemical parameters for cases classified as NS, NSML, or cancer within the expert-curated NSEuroNet (www.nseuronet.com) and COSMIC databases (*74*), respectively (**Fig. 4C**). These databases record the number of cases and specific disease subtypes associated with each variant, enabling statistical analyses of biochemical features across clinical phenotypes. Compared to NS/NSML cases, cancer cases tend to carry variants with a higher fraction active and activator sensitivity (both *p* < 0.01) (**Figs. 4D**&**S31A**). This distinction may arise because strongly activated germline variants are embryonically lethal and therefore observed only as non-inherited somatic mutations (*75*, *76*); conversely, many weakly activated variants observed in developmental disorders may lack sufficient oncogenic drive to cause cancer. Consistent with this trend, NS/NSML patients carrying variants with higher fraction active show an increased leukemia risk (**Fig. S32**). Within cancers, blood cancer cases carry variants with greater fraction active and activator sensitivity than do solid cancer cases (both *p* < 0.01) (**Figs. 4D**&**S33A**), with JMML cases carrying variants showing the largest increases (**Fig. S33B**).

Previous observations of hypoactive variants in solid cancer cases have led to the proposal that SHP2, a well-established proto-oncogene, may also act as a tumor suppressor, suggesting dual roles in cancer (*15*, *45*). While we also identified a subset of cancer cases involving variants with decreased basal activity (3% of blood, 20% of solid cancer cases) (**Fig. S34**), our comprehensive phenotyping offers an alternative interpretation. Regardless of basal activity, most cancer cases carry variants with significantly increased fraction active or activator sensitivity (1244/1260 cases with variants of increased, and 75/87 cases with variants of reduced basal activity) (**Fig. S34**), suggesting that increased open-state propensity, common to variants with both increased and decreased basal activity, is the primary biochemical determinant of SHP2-related oncogenesis.

It has been reported that NS variants associate with elevated basal activity, while NSML variants associate with diminished basal activity due to lower intrinsic activity (*41–44*). Here, a larger dataset (72 NS and 11 NSML variants, as assigned in NSEuroNet) confirms that NS and NSML variants form distinct clusters in biochemical space, consistent with these prior observations (**Figs. 4E**&**S35**). Three variants (N200Y, R501K and Q510L) deviated from this pattern; further investigation revealed that N200Y was misannotated as NSML-associated and instead causes NS (NSEuroNet, personal communication). In addition, we suggest that Q510L and R501K, currently annotated as NS, are more likely NSML based on their biochemical characteristics (**Figs. 4E**&**S35**); supporting this hypothesis, the related variant Q510E is associated with NSML.

Overall, an increased propensity to open, driven by either higher fraction active or activator sensitivity, appears to drive pathogenesis for developmental disorders and cancer. A greater magnitude of this effect predisposes to cancer, and additional reductions in intrinsic activity further stratify specific disease subtypes (**Fig. 4F**). Thus, our high-throughput, multi-dimensional biochemical pipeline can effectively identify functional abnormalities associated with disease types and propose likely disease outcomes for allelic variants.

### Shifts in SHP2 conformational equilibrium alter allosteric inhibitor sensitivity

Allelic variation can impact the potency of drug inhibition and thus the choice of clinical treatment regimens. Here, we quantified the impacts of SHP2 variants on inhibition by three allosteric drugs advanced to clinical trials (TNO155, RMC-4630, and GDC-1971) (*33–36*). Crystal structures show that these compounds bind at the interface between the nSH2-cSH2 linker, cSH2 domain, cSH2-PTP linker, and PTP domain, with the nSH2 domain blocking the active site in a manner closely resembling the autoinhibited state (**Fig. 5A**) (*34*, *35*). These structures thus support a conformational selection mechanism in which these drugs stabilize the autoinhibited state to inhibit SHP2 (**Fig. S36A**) (*48*).

**Figure 5.**
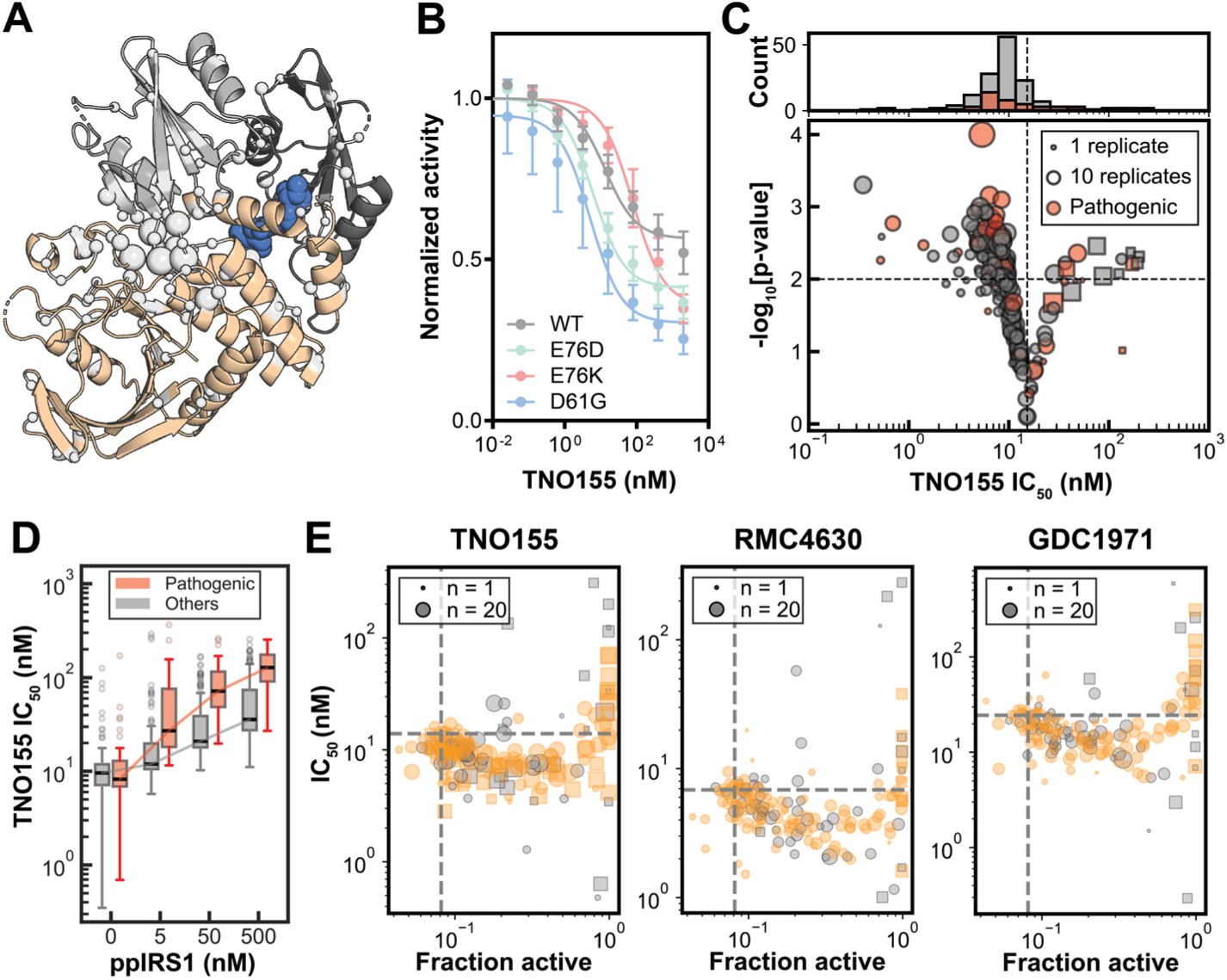
Profiling SHP2 variants reveals variable allosteric inhibitor sensitivity. **(A)** Crystal structure of TNO155-bound SHP2 (PDB ID:7JVM); the drug (blue) is located between the PTP and SH2 domains. White spheres indicate positions mutated in the library; sphere size reflects mutation count. **(B)** HT-MEK measured inhibition dose-response curves for select variants. Data shown in mean ± SEM. **(C)** Volcano plot showing significance of difference from WT (*p*-values) versus median TNO155 IC_50_ for 190 SHP2 variants. Red: Pathogenic variants in ClinVar. Variants with IC_50_ measurements outside the range of inhibitor concentrations used are plotted in squares. **(D)** TNO155 IC_50_ across different ppIRS1 concentrations. Pathogenic variants are marked in red. **(E)** TNO155, RMC-4630 and GDC-1971 IC_50_ versus 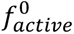. Gold: variants with mutation > 15 Å away from the bound drug. Dash lines show WT values.

Because cellular inhibition occurs in the presence of activating ligands (*e.g.,* ppIRS1) that shift SHP2 toward the open, active conformation and reduce inhibitor potency (**Fig. S36B**) (*26*), we measured DiFMUP hydrolysis across 9 drug concentrations and 4 ppIRS1 concentrations. Fitting dose-response curves with 1:1 binding isotherms (with a Hill slope fixed at 1) yielded an IC_50_ at each ppIRS1 concentration per drug per variant (**Figs. 5B**, **S37-S50**); these values were consistent with plate reader measurements and correlated with drug binding affinity measured directly by microscale thermophoresis (**Fig. S37D**).

In the absence of ppIRS1, measured IC_50_ values varied by ∼100-fold across SHP2 variants for each allosteric inhibitor, with many pathogenic variants exhibiting increased drug sensitivity relative to WT (**Figs. 5C**&**S50**). Under the canonical two-conformational-state inhibition model (**Fig. S36A**), changes in inhibitor IC_50_ couldreflect differences in 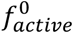 and/or the affinity of the drug for the closed state 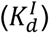 (**Supplementary Text 2**):

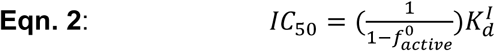

Many mutations with impacts on IC_50_ were far from the drug binding site (**Fig. S51**) and measured IC_50_ values were well-correlated for the three drugs despite their different structures (**Fig. S52**), suggesting most of the observed differences in IC_50_ arise from changes to 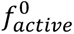 and not ligand binding affinity in the closed state 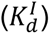.

Consistent with prior literature and an antagonistic binding model, increasing ppIRS1 reduced potency of all three allosteric inhibitors across all tested SHP2 variants (**Figs. 5D**&**S53**), and variants with a greater fraction active exhibited a stronger dependence of IC_50_ on ppIRS1 levels (**Fig. S53**) (*26*). Thus, in cancer cells with higher concentrations of SHP2-activating ligands (*77*, *78*), SHP2 will be less sensitive to these allosteric inhibitors, and this effect will be even more pronounced for cancer cells bearing variants with higher fraction active. Overall, the potency of allosteric inhibitors depends strongly on the identity of the variant and conditions within the cell. These data therefore define biomarkers (*i.e.* variants and activator levels) that can identify patients most likely to respond to therapy, and provide mechanistic support for combination strategies that suppress upstream activation to enhance therapeutic efficacy (*79*).

### The canonical two-conformational-state model fails to explain allosteric inhibition by clinical drugs

Beyond providing a biochemical phenotype for each SHP2 variant, the suite of kinetic and thermodynamic parameters measured here provides a unique opportunity to rigorously test mechanistic models of SHP2 function. As detailed previously, SHP2 allosteric inhibition is typically represented using a two-conformational-state model, in which the enzyme can be either open (active) or closed (autoinhibited), and drugs bind and stabilize the closed state (**Fig. S36A**, **Eqn. 2**, **Supplementary Text 2**) (*48*). However, two experimental observations are at odds with this model. First, this model predicts that SHP2 activity should be completely inhibited at high drug concentrations (**Fig. S54**). Contrary to this prediction, substantial residual SHP2 activity persists at high drug concentrations when using either DiFMUP (**Figs. S37**&**S55**) or the larger peptide substrate EGFRpY992 (**Fig. S56**). Second, this model predicts that more open variants should be less sensitive to drugs if the drug affinity in the closed state 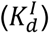 is unchanged (**Fig. S57A**). Contrary to this prediction, IC_50_ values for TNO155, GDC-1971, and RMC-4630 initially decreased with increasing 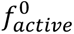 (**Figs. 5E**&**S57B**). This discrepancy was consistent across all three drugs in HT-MEK assays, plate-reader activity assays, and direct binding measurements (**Fig. S58**). Moreover, this discrepancy remained even when considering only variants >15 Å away from the inhibitor in the crystal structure, eliminating those mutations most likely to impact 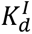 (**Figs. 5E**&**S59**). These findings establish that the current two-conformational-state framework is an inadequate description of SHP2 allosteric inhibition and point to the need for a revised mechanistic model.

### Accounting for variant-specific differences in inhibition requires a partially active intermediate state that preferentially binds drugs

The simplest model capable of explaining all observations is a three-conformational-state model that introduces an intermediate state (*E_i_*) in addition to the canonical closed (*E_c_*) and open (*E_o_*) states (**Figs. 6A-C**, **S54B**, **S60**; **Supplementary Texts 3**&**4**). Such a partially-open, intermediate conformation has also been proposed by recent single-molecule and small-angle X-ray scattering analyses (*80*, *81*). The suite of biochemical measurements obtained here provides support for the functional relevance of this state and critical new insights into its properties. To reproduce the observed incomplete inhibition and relationship between IC_50_ and 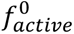, this intermediate state must: (1) possess residual catalytic activity (*r* · *k*, 0 < *r* < 1), and (2) bind drug more strongly than the closed state (*α* < 1) (**Supplementary Text 3**). We note that the model presented here represents a minimal model, and the true model may be yet more complicated (**Supplementary Text 5**).

**Figure 6.**
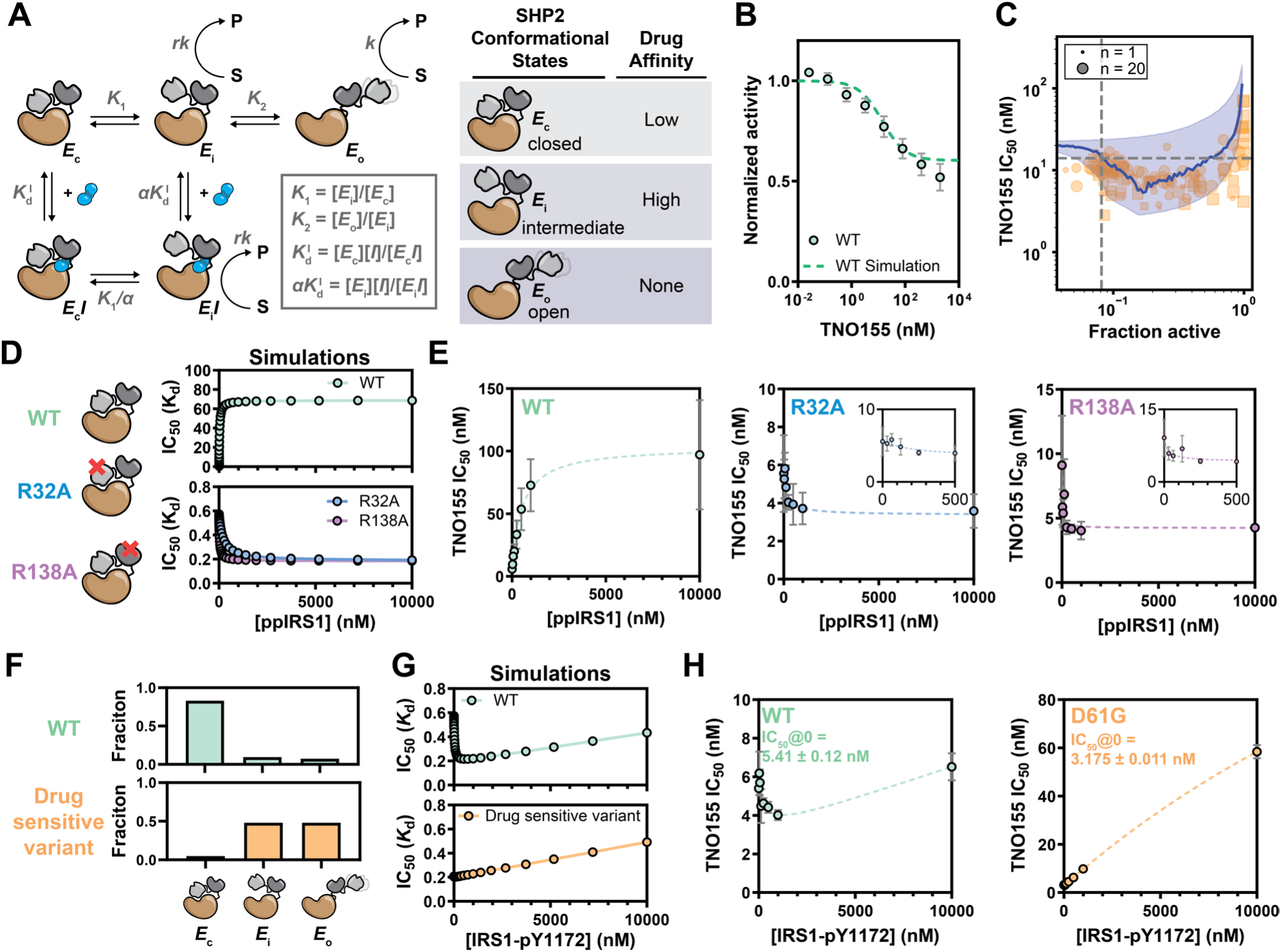
A partially active intermediate conformation is the primary target for clinical allosteric inhibitors. **(A)** Schematic of the three-conformational-state model of SHP2 conformational states and inhibition. **(B)** Simulated dose-response curve using the three-conformational-state model overlaid with experimental data for WT (Mean ± SEM). **(C)** Simulated IC_50_ versus 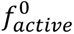 data from the three-conformational-state model (100,000 simulations). Blue line and shading: binned median and 95th percentile. Gold: TNO155 data for variants with mutations >15 Å from drug. **(D)** Simulated inhibitor IC_50_ dependence on ppIRS1 for WT, R32A, and R138A. **(E)** Experimental data of TNO155 IC_50_ dependence on ppIRS1 concentration for WT (left), R32A (center), and R138A (right). Data shown in mean ± SEM and fit to a general binding coupling model. **(F)** Simulated conformational state distribution of WT and a hypothetical drug sensitive variant. **(G)** Simulated inhibitor IC_50_ dependence on IRS1-pY1172 for WT (top) and a drug sensitive variant (bottom). **(H)** Experimental data of TNO155 IC_50_ dependence on IRS1-pY1172 concentration for WT (left) and D61G (right). Data shown in mean ± SEM and fit with a dual coupling model. See **Materials and Methods** for details of all simulations.

### Biasing SHP2 towards the intermediate state potentiates allosteric inhibition

The three-conformational-state inhibition model uniquely predicts that SHP2 will be: (1) more potently inhibited by allosteric drugs if shifted into the intermediate state, and (2) less potently inhibited if shifted into the open state. Prior single-molecule studies indicate that engaging a single SH2 domain with pTyr peptides stabilizes an intermediate conformation, whereas simultaneous engagement of both SH2 domains favors the fully open conformation. If this intermediate conformation corresponds to the highest drug-affinity state proposed in our three-conformational-state model, engagement of a single SH2 domain should enhance inhibition potency, while engagement of both should reduce it.

To test these predictions (**Figs. 6D**&**S61**), we mutated the key arginine in either nSH2 (R32A) or cSH2 (R138A) to impair binding to pTyr peptides so that the activator ppIRS1 is expected to only engage a single SH2 domain in each mutant. In both cases, increasing ppIRS1 concentration lowered the IC_50_ comparably (**Fig. 6E**), consistent with the prediction that engagement of a single SH2 domain populates an intermediate state with enhanced drug affinity. By contrast, when ppIRS1 could engage both SH2 domains (*i.e.* WT), the equilibrium shifted towards the open state, increasing IC_50_ (**Fig. 6E**). This increase saturated at high ppIRS1 concentrations, consistent with the three-conformational-state model prediction.

The three-conformational-state model further suggests that observed variant-specific increases in drug sensitivity (lower IC_50_ values) result from increased occupancy of the intermediate state (which has higher affinity for inhibitors) (**Fig. 6A**&**F**). The singly phosphorylated IRS1-pY1172 peptide binds nSH2 ∼250-fold more strongly than cSH2 (*82*). Therefore, for the predominantly closed WT SHP2, increasing IRS1-pY1172 concentration should initially only engage nSH2, stabilize the intermediate state, and lower IC_50_; as IRS1-pY1172 concentration increase further, additional engagement of cSH2 should drive SHP2 into the fully open state and increase IC_50_ (**Fig. 6G**). Consistent with these predictions, increasing IRS1-pY1172 initially decreased, and then increased, IC_50_ for WT SHP2 (**Fig. 6H**). By contrast, increasing IRS1-pY1172 concentration for a drug-sensitive mutant (*i.e.* D61G), which is expected to predominantly occupy the intermediate state, should only increase IC_50_ (**Fig. 6G**). As expected, increasing IRS1-pY1172 exclusively increased IC_50_ for D61G (**Fig. 6H**).

Notably, SHP099 (the first-in-class allosteric SHP2 inhibitor) behaved differently from TNO155, RMC-4630, and GDC-1971 (*26*, *32*). For SHP099, variants with a higher open fraction than WT consistently displayed higher IC_50_ (**Fig. S62A**). Thus, the model predicts that SHP099 does not preferentially bind the intermediate state over the closed state (*αα* ≥ 1) (**Supplementary Text 3**). Consequently, enrichment in the intermediate state via nSH2 engagement provides no advantage for SHP099 binding, and increasing IRS1-pY1172 should only increase SHP099 IC_50_ (**Fig. S62B**). Measured SHP099 IC_50_ values in the presence of increasing IRS1-pY1172 matched the prediction (**Fig. S62C**), providing additional evidence supporting the three-conformational-state inhibition model.

### Cellular evidence supports in vitro predictions of SHP2 inhibition

To test if *in vitro* drug profiling predicts SHP2 inhibition in cells, we treated U2OS cells containing transiently-transfected SHP2 variants with inhibitors and quantified SHP2 activity by assessing pERK levels via Western blot. Overexpression (typically >20-fold over the endogenous level) of SHP2 WT did not alter inhibitor sensitivity, whereas expression of the drug-resistant mutant E76K caused a substantial reduction of sensitivity, indicating that the assay can detect variant-induced shifts in sensitivity (**Figs. S63-S66**). We found that cellular IC_50_ values correlated with *in vitro* IC_50_ values measured under mildly activating conditions (0 or 5 nM ppIRS1) (Pearson *r* (log-log) = 0.84-0.93; **Figs. S67**&**S68**), though the variability of the cellular assay limited our ability to resolve statistically significant differences. As predicted, variants with intermediate 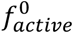 were more or similarly sensitive to TNO155, RMC-4630, and GDC-1971 compared to WT, consistent with stronger inhibitor binding to the intermediate state (**Fig. S69**).

## Discussion

The number of identified human allelic variants is growing rapidly (*1–4*), with most classified as VUS (*10*). Even for variants annotated as benign or pathogenic, we typically lack quantitative information about their biochemical consequences. As shared biochemical alterations drive similar clinical phenotypes (*63*), linking variants to their biochemical effects can improve genetic test interpretation, clinical management, and the development of targeted therapeutics (*83*, *84*), even for rare or previously unobserved variants. Here, we systematically quantify how 190 SHP2 allelic variants alter ten thermodynamic and kinetic parameters that describe SHP2 stability, catalysis, and regulation, yielding (to our knowledge) the largest quantitative biochemical dataset for allelic variants to date.

Quantifying variant effects across multiple distinct biochemical functions overcomes key limitations of current high-throughput experimental and computational approaches to profile variants. Multiplexed assays of variant effects (MAVEs) can evaluate thousands of variants in cells (*65*, *85*, *86*), and prior SHP2 deep mutational scanning yielded enrichment scores that correlate with our measured *k*_cat_/*K*_M_ values (**Fig. S70**) (*45*). However, the design of MAVEs often assumes that catalytic output is the primary predictor of pathology; for SHP2, our data show this assumption does not hold. Moreover, cellular assays report convolved outputs and lack the systematic control needed to directly quantify more complex properties such as autoinhibition and activator affinity, yet these proved to be the principal determinants of SHP2 pathogenesis. Beyond experimental assays, machine learning models such as AlphaMissense provide binary pathogenicity predictions but lack mechanistic interpretability (62, 63, 80), thus providing no biochemical understanding of why a given variant is pathogenic or ability to discriminate between pathogenic modalities. High-throughput biochemical data extend these approaches: defining the biochemical parameters most relevant to disease can inform the design of cellular assays that better approximate these parameters at greater scale, while quantitative biochemical datasets provide ground truth data to interpret existing and train next-generation computational models.

Systematic biochemical dissection of SHP2 variants enhances disease classification and establishes a foundation for disease mechanistic hypotheses. NS and NSML exhibit overlapping phenotypes that frequently confound clinical diagnosis (*39*, *40*); our data, consistent with prior reports but now expanded across a broader variant space (*25*, *44*), suggest that dysregulated autoinhibition drives their common phenotypes while deficits in intrinsic catalysis account for NSML-specific symptoms such as lentigines. In cancer cases, nearly all somatic missense variants—including those previously interpreted as consistent with a tumor-suppressive role (*15*, *33*, *45*)—exhibit dysregulated autoinhibition. Among these, variants with reduced catalytic activity are tolerated almost exclusively in solid cancers, suggesting hematologic malignancies may additionally require catalytic output at or exceeding wild-type levels. As dysregulated autoinhibition is the shared feature across all pathogenic classes regardless of catalytic activity, we speculate that SHP2 variants primarily promote disease through catalysis-independent scaffolding functions (*15*, *66–69*).

Beyond classification, thermodynamic and kinetic constants for hundreds of variants enabled systematic evaluation of mechanistic models for SHP2 function. Here, modeling of variant inhibition data revealed that three clinical-stage allosteric inhibitors preferentially bind a partially active intermediate state, whereas the first-generation inhibitor SHP099 does not. Although crystal structures depict all SHP2 inhibitors bound to the fully closed conformation (*32*, *34*, *35*), these static snapshots may not represent the complete conformational ensemble sampled under physiological conditions (*87–89*). The observed shift in binding properties from SHP099 to the more potent clinical-stage inhibitors illustrates how drug optimization can inadvertently shift the targeted conformation of allosteric inhibitors in ways not apparent from static structures. Integrating a panel of conformationally diverse variants into lead optimization can benchmark compound activity against mechanistic models and enable early identification of unintended shifts in conformational selectivity before they advance undetected into clinical development.

While allosteric inhibitors have the potential for greater selectivity and the ability to modulate (rather than block) activity (*90*), we highlight four challenges associated with allosteric SHP2 inhibition. First, many cancer variants preferentially adopt the open conformation, rendering them intrinsically less sensitive to compounds that bind the closed or intermediate conformations. Second, as allosteric inhibition relies on structure-wide conformational coupling, resistance-conferring mutations can arise throughout the protein (*90–92*). Third, preferential binding to an intermediate state with residual activity means that allosteric SHP2 inhibitors will not completely suppress activity, even at saturating concentrations. Fourth, drug binding to SHP2 competes with binding of endogenous activators, creating a conformational “tug-of-war”. This tug-of-war inverts the therapeutic window such that inhibitors exert weaker effects in cancer cells with elevated activators and stronger effects in normal tissues, potentially diminishing on-target potency while heightening the risk of on-target, off-tumor toxicity. As similar limitations have been observed in other drug targets including tyrosine kinases and Ras (*93–95*), these issues may reflect fundamental constraints of allosteric inhibition as a therapeutic strategy.

Restricting activator engagement to a single SH2 domain biases the conformational ensemble toward the intermediate state, enhancing potency for clinical allosteric inhibitors. This suggests a potential therapeutic strategy in which compounds that selectively block one SH2 domain could be combined with allosteric inhibitors to maximize potency in hyperactivated disease contexts. Alternatively, inhibitors that target the PTP domain could bypass this conformational tug-of-war entirely, or even exploit it synergistically if the target conformation is only accessible in the open state. Such inhibitors could act through orthosteric inhibition or by targeting allosteric sites within the PTP domain that operate independently of SH2-mediated autoinhibition (*96*, *97*); our data identified one such candidate site (**Fig. S13**).

We have demonstrated that systematic, multidimensional biochemical profiling of clinical SHP2 variants can expose profound and consequential mechanistic complexity—in both disease etiology and drug response—that remained elusive in one of the most studied protein tyrosine phosphatases in disease biology. SHP2 is unlikely to be exceptional in this regard. Extending this biochemistry-centered approach broadly across other clinically important proteins will be essential to transform the growing catalog of human variants from a challenge into a mechanistic and therapeutic opportunity for advancing precision medicine.

## Acknowledgements

We thank members of the Herschlag and Fordyce laboratories for discussions and review of the manuscript, and Dr. David A Stevenson and Dr. Bruce D Gelb for helpful discussions.

## Author contributions

A.A.L., D.A.M., D.H. and P.M.F. conceived the project. A.A.L. and D.A.M. designed experiments and collected and analyzed data. E.D.E. collected and analyzed data. A.A.L. performed modeling and simulations. S.C.B., D.H., and P.M.F. supervised the project. A.A.L., D.H., and P.M.F. wrote the manuscript, with input from all authors.

## Data and materials availability

Summary tables of all kinetic and thermodynamic parameters measured for each variant are included in the supplementary materials and available on the Fordyce Lab website (www.fordycelab.com/data). All data acquired in this study are available in a registered Open Science Foundation Repository (https://doi.org/10.17605/OSF.IO/YGHQV).

