## Supplementary information for "High-throughput biochemical phenotyping of SHP2 variants reveals the molecular basis of diseases and allosteric drug inhibition"

### Materials and Methods

#### SHP2 variant library generation

The C-terminally truncated human SHP2 isoform 1 (NP_002825.3) comprising residues 1-525 ("full length," $SHP2_{FL}$) and a further truncated construct lacking the tandem SH2 domains (residues 225-525; "catalytic domain," $SHP2_{CD}$) were fused to EGFP at the C-terminus via a flexible Ser/Gly linker (GGGSGGGGSG). Coding sequence for each SHP2-EGFP fusion was cloned into a linearized NEB PURExpress DHFR control vector with DHFR gene removed using HiFi DNA Assembly (New England Biolabs) following the manufacturer's instructions.

Variants in the library were identified from the ClinVar database (as of December 2022) from single missense mutation variants within residues of 1-525 of SHP2 isoform 1 (NM_002834.5). QuikChange site-directed mutagenesis was used to generate the variant library. Briefly, mutagenic forward and reverse oligonucleotide primers were designed using an in-house design script (available at https://github.com/FordyceLab/designQuikChangePrimers). Site-directed mutagenesis was performed in 96-well plate format with either $SHP2_{FL}$-EGFP or $SHP2_{CD}$-EGFP plasmids as templates using the QuikChange Lightning Site-Directed Mutagenesis Kit (Agilent), scaled to 1/5 of the recommended reaction volume (10 µL PCR). One µL of the resulting DpnI digested reaction solution was then used to transform 5 µL DH5alpha ultracompetent E. coli (New England Biolabs) in 96-well plate format, according to manufacturer's protocol. Transformed cultures were plated onto LB (Luria broth) agar with 100 µg/mL carbenicillin. Resulting transformants were cultured to saturation in 4 mL LB medium supplemented with carbenicillin (100 µg/mL). Plasmids were purified using QIAprep Spin Miniprep Kits (Qiagen) or ZymoPURE Plasmid Miniprep Kits (Zymo Research). All variants were confirmed by Sanger sequencing.

#### HT-MEK microfluidic device design and fabrication

Design files for HT-MEK microfluidic devices are available on the Fordyce Lab website (http://www.fordycelab.com/microfluidic-design-files). Silicon wafer molding masters and HT-MEK microfluidic polydimethylsiloxane (PDMS) devices were fabricated as described previously (*1*).

#### SHP2 variant library printing and device assembly

Enzyme plasmid microarray printing was performed similarly to previously reported (*1*). Briefly, each plasmid was diluted in a filter-sterilized solution of 5% (w/v) BSA, 60 mg/mL trehalose dihydrate, and 375 mM NaCl to a final concentration of 10-50 ng/µL plasmid in a solution of 1% (w/v) BSA, 12 mg/mL trehalose dihydrate, and 75 mM NaCl, and deposited into a 384-well plate.  Plasmid DNA solutions were printed onto epoxysilane-coated glass (Arrayit corporation) slides using a SciFlex Arrayer microarray printer (SCIENION AG) equipped with either a PDC50 or PDC70 nozzle. Plasmid DNA arrays were printed with 896 spots (for 896 of 1792 total chambers), to align with odd-rowed DNA chambers only within the HT-MEK device (skipping every other chamber). This is done to prevent cross contamination during assays and enable better quality control of measurements. The printed DNA array was dried under vacuum for at least 24 hours and stored under vacuum until further use. HT-MEK PDMS device was aligned to the printed DNA array. The aligned devices were then baked for 4 hours at 95ºC on a hotplate for bonding.

#### Imaging and pneumatics setups

Microfluidic devices were imaged using a Nikon Ti-S microscope and operated using a custom pneumatic manifold, which was synchronized with microscope acquisition through an in-house Python package (RunPack; https://github.com/FordyceLab/RunPack) that provides unified control of both the microscope and valve hardware. RunPack interfaces with the microscope via the pycromanager library (*2*), which uses open-source μManager microscope hardware package (*3*). The Nikon Ti-S Microscope is equipped with a motorized XY stage (Applied Scientific Instrumentation, MS-2000 XYZ stage), cMOS camera (Oxford Instruments, Andor Zyla 4.2), solid-state light source (Lumencor, SOLA SE Light Engine), and automated filter turret equipped with an eGFP filter set (Chroma Technology Corp., part no. 49002), DAPI filter set (Semrock Inc., catalog no. DAPI-1160B-NTE), and a custom "PBP" filter set (Semrock Inc., 427/10 bandpass excitation filter, catalog no. FF01-427/10-25; 470/22 bandpass emission filter, catalog no. FF01-470/22-25; 442 nm dichroic beamsplitter, catalog no. DI03-R442-T1- 25X36; mounted in a TE2000 filter cube, catalog no. NTE). All imaging was performed using a 4X objective (CFI Plan Apochromat $\lambda$ 4X NA 0.20, Nikon) at 2x2 binning (1024x1024 pixels).

**On-chip SHP2 variant library protein expression and purification**

HT-MEK devices aligned to plasmid DNA microarray prints were chemically patterned as previously described to deposit a surface-immobilized patch of biotinylated anti-GFP antibody (GFP Antibody (9F9.F9) [Biotin], Novus Biologicals, NB110-40670) beneath the Button valve. To express enzymes, we introduced a PURExpress in vitro transcription/translation mixture (New England Biolabs) into the device. Briefly, 12 μL of component A and 8 μL of component B were mixed and incubated on ice for 30 min. We then added 0.6 μL of 40 U/μL recombinant RNase inhibitor (Promega, N2515), 1 μL of 7.5 mM DTT, 1 μL of 3 mM TCEP·HCl, and 6.4 μL of UltraPure DNase/RNase-Free Distilled Water (Invitrogen). This reaction mixture was flowed into the device with the Neck valves closed, thereby preventing access to the DNA chambers. When most of the solution had passed through, the outlet valve was closed and the Neck valves were opened to dead-end fill the DNA chambers containing printed plasmid spots. Once the chambers were filled, the Neck valves were immediately closed to prevent cross-contamination.

For expression, devices were incubated on a hot plate at 30 °C for 1 h, followed by 1 h at room temperature to allow GFP maturation. All steps were performed in the dark to prevent photobleaching. During the expression period, reaction chambers were continuously flushed with reaction buffer (50 mM MOPS pH 7.0, 100 mM NaCl, 100 μM EDTA, 1 mM TCEP, 1 mg/mL BSA (ThermoFisher AM2616), 0.05% Tween-20 (w/v), and 3 mM DTT). Following expression, enzyme was bound to the surface-patterned Button valve by closing Sandwich valves and opening the Button and Neck valves. After sufficient binding, Button valves were closed, Sandwich valves opened, and excess reaction mixture was flushed from the device with reaction buffer for ~1 h or until no residual EGFP signal remained in the DNA chambers. Button fluorescence intensities were then measured via GFP-channel imaging. An on-chip standard curve relating summed button EGFP fluorescence to absolute EGFP concentration was used to calculate the effective concentration of each enzyme fusion construct, as previously described (*1*).

**Measuring SHP2 enzyme turnover with HT-MEK**

*DiFMU standard curve*

Fluorogenic leaving-group standard curves for DiFMUP were measured using 6,8-difluoro-4-methylumbelliferone (DiFMU; Invitrogen, D6566). Standard curves relating chamber fluorescence intensity to absolute product concentration were obtained by imaging a series of DiFMU solutions prepared in reaction buffer at the following concentrations: 0, 1, 2, 5, 10, 20, 50, 100 μM. Tubing lines loaded with each standard solution were inserted into the device inlet ports, and solutions were sequentially flowed through the device with Button valves closed to equilibrate all chambers for ~5 min. After each equilibration step, Sandwich valves were closed, Button valves were opened, and the device was imaged in the DAPI channel to quantify chamber fluorescence.

*DiFMUP hydrolysis Michaelis-Menten kinetics*

DiFMUP (Invitrogen, D6567) powder was dissolved in Milli-Q water to 100 mM, stored at –20 °C, and used within 3 months. A dilution series was prepared in reaction buffer to yield final substrate concentrations of 10, 20, 50, 100, 200, 500, 1000, 2000 μM. Each DiFMUP solution was sequentially flowed through the device with Button valves closed, thereby preventing enzyme access, for ~5 min to equilibrate all chambers. Reactions were initiated by closing the Sandwich valves and opening the Button valves. Product formation was monitored by time-series fluorescence imaging of the device in the DAPI channel. After 30–60 min, the Button valves were closed to protect the immobilized enzyme, and the device was equilibrated with the next higher substrate concentration.

*EGFRpY992 hydrolysis standard curve and Michaelis-Menten kinetics*

EGFRpY992 (Ac-DADEpYLIPQQG-amide) was custom synthesized by GenScript (≥95% purity). Hydrolysis of EGFRpY992 was measured using a fluorescent phosphate-binding protein (PBP; Phosphate Sensor, Thermo Fisher, PV4406). Binding of inorganic phosphate (Pi) by PBP produces an increase in fluorescence; excitation was 427/10 nm and emission was collected at 470 nm using a custom PBP filter set. Per-chamber standard curves were prepared in phosphate-free reaction buffer with [Pi] = 0, 2, 5, 10, 20, 50, and 100 µM and nominal [PBP] = 100 µM. Note that since commercially supplied BSA contains high levels of phosphate, we performed extensive buffer exchange to prevent phosphate contamination of the reaction buffer. PBP binding was modeled as a single-site binding isotherm relating median chamber fluorescence $I([Pi])$ to [Pi]:

$$I([Pi]) = 0.5A(K_D + [Pi] + [PBP] - \sqrt{(K_D + [Pi] + [PBP])^2 - 4[PBP][Pi]}) + I(0)$$

$I([Pi])$ is the median fluorescence at a given [Pi], [PBP] is the concentration of total PBP, $K_D$ is the PBP-Pi dissociation constant, and A is a fluorescence scaling factor. To convert measured fluorescence to [Pi], we performed a nonlinear least-squares fit of the measured standard intensities to the model, with A, I(0), [PBP], and $K_D$ as free parameters on a per-chamber basis. The fitted curve was then used as a standard curve to convert experimental fluorescence to [Pi] for the kinetic assays. We performed EGFRpY992 hydrolysis reactions typically with concentrations of 5, 10, 20, 50, 100, 200, 500, 1000 µM, supplemented with 100 µM PBP in reaction buffer. Assays were carried out using the same workflow as the DiFMUP hydrolysis measurements, with fluorescence detected using the PBP filter cube.

*Urea unfolding assays*

Assays were conducted using 100 µM DiFMUP in reaction buffer, with urea concentrations of 0, 0.25, 0.5, 1, 1.5, 2, 2.5, and 3 M. Each urea concentration was introduced sequentially from lowest to highest, allowed to equilibrate for 5–7 minutes, and then imaged for approximately 30 minutes. Following each change in urea concentration, a 0 M urea solution was flowed through the device, and three consecutive equilibration-and-imaging cycles (5–7 minutes equilibration followed by ~30 minutes imaging) were performed to reestablish the baseline before proceeding to the next condition.

*ppIRS1 activation assays*

ppIRS1 (SLNpYIDLDLVK-dPEG8-LSTpYASINFQK-amide) was custom synthesized by GenScript (≥95% purity). ppIRS1 assays were performed using 100 µM DiFMUP in reaction buffer, with ppIRS1 concentrations of 0, 0.0256, 0.128, 0.64, 3.2, 16, 80, 400, and 2000 nM, flowed from low to high. For each ppIRS1 concentration, we performed three consecutive equilibration–imaging cycles (5–7 minutes equilibration followed by ~30 minutes imaging). The initial rate from the third cycle was recorded to ensure >1 hour of equilibration for each condition.

*Inhibition assays*

TNO155 (Batoprotafib; CAS No. 1801765-04-7), RMC-4630 (Vociprotafib; CAS No. 2172652-48-9), and GDC-1971 (RLY-1971; CAS No. 2377352-49-1) were purchased from Selleck chemicals. Inhibition assays were performed using 100 µM DiFMUP in reaction buffer, with inhibitor concentrations of 0, 0.0256, 0.128, 0.64, 3.2, 16, 80, 400, and 2000 nM. For each inhibitor concentration, we carried out three consecutive equilibration–imaging cycles (5–7 minutes equilibration followed by ~30 minutes imaging), and the initial rate from the third cycle was recorded to ensure >1 hour of equilibration for each condition.

*Image processing and kinetic fitting*

Image stitching and initial processing was performed via an in-house Python package (ImageStitcher and ProcessingPack) previously reported (*1*). Initial rate fitting of extracted chamber intensities and downstream processing was performed in Python, with fitting algorithms and procedures as previously reported and re-implemented in Python (*1*), with minor adjustments. Briefly, to further ensure high-quality kinetic fits, we applied adaptive goodness-of-fit thresholds based on the fraction of substrate consumed at each timepoint. For very early timepoints, where the signal is low and noise can disproportionately influence fits, we required only a modest goodness-of-fit ($r^2 \geq 0.9$ for data points corresponding to <2% substrate consumption). As the reaction progressed and signal-to-noise ratio improved, stricter criteria were applied: $r^2 \geq 0.99$ for <5% substrate consumption, $r^2 \geq 0.999$ for <10% substrate consumption, and $r^2 \geq 0.9999$ for all remaining data points. Using these criteria, all initial rate fits included more than three data points and achieved at least $r^2 \geq 0.9$ for <10% substrate consumption, ensuring robust estimates of initial rates.

*Fitting Michaelis-Menten parameters*

The parameters $k_{cat}$, $K_M$, and $k_{cat}/K_M$ were determined from a non-linear least-squares fit of the initial rates at each substrate concentration to the Michaelis-Menten equation:

$$v_i = \frac{k_{cat}[E][S]}{K_M + [S]}$$

To ensure removal of datapoints with problematic measurements due to inhomogeneous flow or incomplete substrate equilibration, we applied criteria to exclude certain substrate concentrations from the fit. Specifically, any substrate concentration was excluded if the measured initial rate was negative (except the first concentration, which can sometimes be near zero), or if the rate was lower than 90% of any preceding (lower) substrate concentrations in the series. Chambers with less than four valid points were flagged and excluded from downstream analysis.

Accurate determination of $K_M$ requires measurements of initial rates at concentrations significantly above and below the true $K_M$ value. For cases with fitted $K_M$ values exceeded the highest substrate concentration used, we set $K_M$ to this maximum concentration and flagged the value as a lower limit. Conversely, cases with fitted $K_M$ values below the lowest substrate concentration would have been flagged as upper limits; however, no measurements in this dataset required this adjustment.

*Fitting stability parameters*

Urea-induced unfolding of each chamber was analyzed by calculating the fraction of native protein at each urea concentration. For each chamber, initial rates measured at different urea concentrations were used to compute native fractions relative to a reference condition (determined by the initial rate at 0 M urea measured immediately before each urea concentration):

$$f_{native} = \frac{v_i}{v_0}$$

To minimize artifacts, urea concentrations were excluded if the native fraction was negative or if it increased by more than 10% relative to the previous lower urea concentration, indicating potential measurement errors or noise. Chambers with less than four valid points were flagged and excluded from downstream analysis. The remaining valid points were then used to fit a two-state linear extrapolation model:

$$f_{native} = \frac{(Y_{max} - Y_{min})e^{-(\Delta G^0 + m[urea])/RT}}{1 + e^{-(\Delta G^0 + m[urea])/RT}} + Y_{min}$$

Where $Y_{max}$ and $Y_{min}$ are the upper and lower initial rates baseline, $\Delta G^0$ is the free energy of active site folding in the absence of urea, *m* is the urea dependence, R is the gas constant, and T is the temperature. Curve fitting was performed using non-linear least-squares regression. For cases with fitted $\Delta G^0$ values more than 0 (more than 50% non-native at 0 M urea), we flagged that measurement as with high uncertainty.

*Fitting activation parameters*

To characterize the dose–response behavior of each chamber, initial rates were fit to a standard Hill equation with a fixed Hill slope of 1:

$$v_i = Bottom + \frac{(Top - bottom)[A]}{[A] + EC_{50}}$$

where [S] is the activator concentration, *Bottom* and *Top* are the minimum and maximum response levels, and $EC_{50}$ is the concentration yielding half-maximal response. For each chamber, the list of measured initial rates and corresponding activator concentrations was extracted. Any data points with negative initial rates were excluded. Chambers with less than four valid points were flagged and excluded from downstream analysis. Initial rates were normalized to the maximum value within each chamber to improve numerical stability during fitting. Curve fitting was performed using non-linear least-squares regression, with initial guesses for *Bottom*, *Top*, and $EC_{50}$ set based on the minimum, maximum, and median values of the data. Parameter bounds were applied to constrain physically reasonable values.

*Fitting inhibition parameters*

To determine the inhibitory potency of compounds in each chamber, initial rates were fit to a standard inhibition curve with a fixed Hill slope of 1:

$$v_i = Bottom + \frac{(Bottom - Top)}{1 + [I]/IC_{50}}$$

where [I] is the inhibitor concentration, *Bottom* and *Top* are the minimum and maximum responses, and $IC_{50}$ is the inhibitor concentration that reduces the response by 50%. For each chamber, measured initial rates and corresponding inhibitor concentrations were extracted. Data points with negative concentrations were excluded from the fit. Non-linear least-squares regression was used to fit the $IC_{50}$ curve, with initial guesses for *Bottom*, *Top*, and $IC_{50}$ based on the observed data.

*Quality control*

To eliminate chambers with levels of expressed SHP2 below the limit of accurate on-chip quantification of enzyme concentration, we automatically eliminated any chambers containing a calculated [E] <0.5 nM. Chambers were also manually inspected and culled if they contained dust or aggregates to eliminate high-intensity spot artifacts.

For activity experiments, ppIRS1-dependent activation experiments, and inhibition experiments, we excluded chambers with fitted initial rates less than twofold above adjacent empty chambers in the same column (to remove potentially contaminated and/or too-slow reactions), or with curve fit $r^2 < 0.9$.

For urea-dependent reaction rate experiments, we excluded chambers with fitted initial rates less than 1.5-fold above adjacent empty chambers in the same column, or with curve fit $r^2 < 0.9$.

*Normalization and statistical tests*

All statistical analyses were performed in Python using custom scripts. We normalized data between experiments prior to aggregation using a linear normalization strategy as described previously. We estimate the uncertainty of measured biochemical parameters for each variant using bootstrap resampling. For each variant, the log-transformed data were resampled with replacement for 10,000 iterations, and the median was computed for each bootstrap sample. The 95% confidence interval (CI) for the mutant median was defined by the 2.5th and 97.5th percentiles of the resulting bootstrap distribution.

To test for significance of the observed differences between variant and WT values, a two-sample bootstrap test was performed on the log-transformed data. The test statistic was defined as the absolute difference between the mutant and WT medians in log space. Under the null hypothesis of no difference between groups, a bootstrap null distribution was generated by pooling mutant and WT log-transformed measurements and resampling with replacement to generate synthetic mutant and WT samples of the same sizes as the original data. For each of 10,000 bootstrap iterations, the absolute difference in medians was recalculated. Two-tailed p-values were computed as the fraction of bootstrap iterations in which the bootstrap statistic exceeded the observed statistic.

To quantify uncertainty in fraction active (i.e., $\left(\frac{k_{cat}}{K_M}\right)_{FL} / \left(\frac{k_{cat}}{K_M}\right)_{CD}$), we computed the standard error of the fraction active using bootstrap resampling. For each variant, the log-transformed data were resampled with replacement independently for the FL and CD measurements for 10,000 iterations. In each iteration, the medians of the resampled FL and CD datasets were computed, and their ratio was recorded. The standard error of the fraction active was taken as the standard deviation of the resulting bootstrap distribution. If the variant has its mutation in the SH2 domain, its CD activity is equivalent to WT CD activity, and thus assigned WT CD distribution for calculation

To test for significance of the observed differences in fraction active between variant and WT, we performed a two-tailed bootstrap hypothesis test on FL/CD ratios in log space. For each group of data ($\left(\frac{k_{cat}}{K_M}\right)_{FL,mut}$, $\left(\frac{k_{cat}}{K_M}\right)_{CD,mut}$, $\left(\frac{k_{cat}}{K_M}\right)_{FL,WT}$, $\left(\frac{k_{cat}}{K_M}\right)_{CD,WT}$), the median of the log-transformed data was computed. The test statistic was defined as the difference between mutant and WT log-ratios:

$$\Delta = \left[median\left(\log_{10}\left(\frac{k_{cat}}{K_M}\right)_{FL,mut}\right) - median\left(\log_{10}\left(\frac{k_{cat}}{K_M}\right)_{CD,mut}\right)\right] - \left[median\left(\log_{10}\left(\frac{k_{cat}}{K_M}\right)_{FL,WT}\right) - median\left(\log_{10}\left(\frac{k_{cat}}{K_M}\right)_{CD,WT}\right)\right]$$

To generate the null distribution, bootstrap resampling was performed independently within each of the four datasets ($\left(\frac{k_{cat}}{K_M}\right)_{FL,mut}$, $\left(\frac{k_{cat}}{K_M}\right)_{CD,mut}$, $\left(\frac{k_{cat}}{K_M}\right)_{FL,WT}$, $\left(\frac{k_{cat}}{K_M}\right)_{CD,WT}$). For each of 10,000 bootstrap iterations, samples were drawn with replacement from each dataset, and the bootstrap estimate of Δ was recalculated using medians as above. The resulting bootstrap distribution of Δ values was centered by subtracting its median, enforcing a null hypothesis of zero difference in log-ratios. Two-tailed p-values were calculated as the fraction of bootstrap samples for which the absolute value of the centered bootstrap statistic was greater than or equal to the absolute value of the observed statistic.

**Measuring SHP2 enzyme turnover with plate reader**

*Protein purification*

SHP2 variants in frame with a C-terminal Strep-tag II were cloned into a pET22b plasmid and transformed into *E. coli* BL21(DE3) expression cells. Transformants were grown overnight at 37 °C in LB medium supplemented with carbenicillin (50 µg/mL). The overnight culture was diluted 1:100 (v/v) into fresh LB medium with carbenicillin (50 µg/mL) and grown at 37 °C to an $OD_{600}$ of 0.6-0.8. Cultures were cooled to room temperature, and protein expression was induced with 0.1 mM isopropyl β-D-1-thiogalactopyranoside (IPTG) at 25 °C with shaking (200 rpm) overnight. Cells were harvested by centrifugation at 4,000 rpm. Pellets were either frozen for later purification or resuspended in column buffer (100 mM Tris pH 8.0, 150 mM NaCl, 1 mM TCEP) and lysed by three passes through an EmulsiFlex-C5 homogenizer (Avestin). Lysates were clarified by centrifugation at 20,000 rpm and filtered through a 0.45 µm membrane. The clarified lysate was applied to a gravity column packed with Strep-Tactin Sepharose resin (Cytiva). The resin was washed with 6 column volumes of column buffer, and bound protein was eluted with 2.5 mM desthiobiotin in column buffer. Fractions were analyzed by sodium dodecyl sulfate-polyacrylamide gel electrophoresis (SDS-PAGE) with Coomassie staining. Fractions containing >95% pure SHP2 were pooled and buffer exchanged into storage buffer (50 mM MOPS (pH 7.0), 100 mM NaCl, 100 µM EDTA, 1 mM TCEP) using 10 kDa MWCO Amicon Ultra-15 centrifugal filter units (MilliporeSigma). The concentration of protein was determined by measuring $OD_{280}$ (ext. coefficient 73800 $M^{-1}cm^{-1}$) using nanodrop, and the protein solution was flash-frozen in liquid nitrogen and stored at -80ºC.

*IVTT protein expression*

In vitro transcription translation of SHP2-EGFP constructs were performed with the same mutant plasmids expressed on chip. Briefly, 12 µL of component A and 8 µL of component B were mixed and incubated on ice for 30 min. We then added 0.6 µL of 40 U/µL recombinant RNase inhibitor (Promega, N2515), 1 µL of 7.5 mM DTT, 1 µL of 3 mM TCEP·HCl, and 6.4 µL of UltraPure DNase/RNase-Free Distilled Water (Invitrogen). Then, 1 µL of SHP2 variant plasmid at 50-100 ng/µL was added to the mixture. Expression was performed in PCR strips incubated in a thermocycler, holding for 1 hour at 30 °C, then 1 hour at 22 °C. Enzyme concentration was measured by diluting the raw IVTT products 1:100 in PBS and quantifying fluorescence with a Denovix fluorometer/spectrophotometer (model DS-11 FX), reading fluorescence (emission 514–567 nm) between from blue-light excitation at ~470 nm and converted to eGFP concentration using a standard curve prepared using a commercial eGFP sample (Biovision, 4999). Enzymes were then diluted 1:200-fold into reaction buffer before assay.

*Plate reader assay setup*

Enzyme turnover was measured using a Tecan Infinite M200 plate reader. All assays were performed in 384-well solid black polystyrene microplates (Corning, 3820). DiFMUP hydrolysis and production of DiFMU was measured with excitation at 358 nm and emission at 455 nm. For EGFRpY992 hydrolysis assays, inorganic phosphate binding by PBP was measured with excitation at 430 nm and emission at 450 nm.

*DiFMU standard curve*

Standard curves relating chamber fluorescence intensity to absolute product concentration were obtained by measuring the fluorescence of a series of DiFMU solutions prepared in reaction buffer at the following concentrations: 0, 1, 2, 5, 10, 20, 50, 100 µM.

*DiFMUP hydrolysis Michaelis-Menten kinetics*

Enzymes were prepared in reaction buffer at 2X the desired final assay concentration (0.1-10 nM in the final reaction). Assays were initiated by mixing equal volumes of the 2X enzyme solution with 2X DiFMUP substrate stocks in reaction buffer. This yielded final (1X) DiFMUP concentrations of 10, 20, 50, 100, 200, 500, 1000, 2000 µM. After mixing, reactions proceeded at room temperature for 2 hours, and fluorescence was monitored every minute to obtain initial reaction velocities.

*Urea unfolding assays*

Urea unfolding assays were performed using 100 µM DiFMUP in reaction buffer across a urea concentration series of 0, 0.25, 0.5, 1.0, 1.5, 2.0, 2.5, and 3.0 M. SHP2 was preincubated with the indicated urea concentrations for 30 minutes (or for the specified time). Reactions were initiated by mixing 10 µL of SHP2 urea solution with 10 µL of 200 µM DiFMUP solution prepared in reaction buffer containing the same urea concentration as the corresponding SHP2 incubation condition, thereby maintaining constant urea levels throughout the assay.

*ppIRS1 activation assays*

ppIRS1 activation assays were performed using 100 µM DiFMUP in reaction buffer, with ppIRS1 concentrations of 0, 0.0256, 0.128, 0.64, 3.2, 16, 80, 400, and 2000 nM. SHP2 was preincubated with the indicated ppIRS1 concentrations for 30 minutes. Reactions were initiated by mixing 10 µL of SHP2 ppIRS1 mixture solution with 10 µL of 200 µM DiFMUP solution prepared in reaction buffer containing the same ppIRS1 concentration as the corresponding SHP2 incubation condition.

*Inhibition assays*

Inhibition assays were performed using 100 µM DiFMUP in reaction buffer, with inhibitor concentrations of 0, 0.0256, 0.128, 0.64, 3.2, 16, 80, 400, and 2000 nM. SHP2 was preincubated with the indicated inhibitor concentrations for 30 minutes. Reactions were initiated by mixing 10 µL of SHP2 inhibitor mixture solution with

10 µL of 200 µM DiFMUP solution prepared in reaction buffer containing the same inhibitor concentration as the corresponding SHP2 incubation condition.

*Inhibition assays in the presence of activating peptide*

Inhibition assays were performed using 100 µM DiFMUP in reaction buffer, with inhibitor concentrations of 0, 0.0256, 0.128, 0.64, 3.2, 16, 80, 400, and 2000 nM, and varying concentrations of the IRS1-pY1172 peptide (SLNpYIDLDLVK; custom synthesized by GenScript at ≥95% purity) or ppIRS1 (0, 31.25, 62.5, 125, 250, 500, 1000,10000 nM). SHP2 was preincubated with the indicated inhibitor concentrations for 30 minutes. Reactions were initiated by mixing 10 µL of SHP2 inhibitor/activator mixture solution with 10 µL of 200 µM DiFMUP solution prepared in reaction buffer containing the same inhibitor/activator concentration as the corresponding SHP2 incubation condition.

*Curve fitting*

Michaelis–Menten parameters, activation dose-response curves, inhibition dose-response curves, and urea unfolding curves from plate reader data were fit using the same models described in the HT-MEK section above.

For fitting inhibition $IC_{50}$ dependence on ppIRS1 concentration (**Fig. 6E**), $IC_{50}$ data were fit to a general binding coupling model:

$$IC_{50} = IC_{50}^{0}(1 + \frac{[A]}{K_d^A})/(1 + \frac{\alpha[A]}{K_d^A})$$

where $IC_{50}^0$ is the $IC_{50}$ in the absence of activator, $K_d^A$ is the apparent dissociation constant for ppIRS1 binding, and $\alpha$ is the coupling coefficient describing the fold-change in $IC_{50}$ at saturating activator. For IRS1-pY1172 (**Fig. 6H**), which can independently engage two SH2 domains, $IC_{50}$ data were fit to a dual-site coupling model:

$$IC_{50} = IC_{50}^{0}\frac{(1 + \frac{[A]}{K_d^{A1}})(1 + \frac{[A]}{K_d^{A2}})}{(1 + \frac{\alpha_1[A]}{K_d^{A1}})(1 + \frac{\alpha_2[A]}{K_d^{A2}})}$$

where $K_d^{A1}$ and $K_d^{A2}$ are the apparent dissociation constants for binding to each SH2 domain. Curve fitting was performed using non-linear least-squares regression.

**Simulations**

*SHP2 two-conformational-state model with inhibition*

The two-conformational-state model (**Fig. S36A**) was simulated by solving the system of equations:

$K = [E_o]/[E_c]$

$[I]/K_d = [E_cI]/[E_c]$

$[E_o] + [E_c] + [E_cI] = 1$

For the simulation of WT dose-response curve in **Fig. S54A**, $K = 0.1$ was used based on our experimental data.

Enzyme activity was modeled as proportional to the open-state population as $\mathrm{k}[\mathrm{E_o}]$, which can also be expressed as:

$$k[E_o] = k\frac{K}{1 + K + \frac{[I]}{K_d}}$$

$k[E_o]$ was plotted as a function of $[I]/K_d$ to get a dose-response curve, such that the x-axis is expressed in units of $K_d$, eliminating the need for an arbitrary choice of $K_d$.

Under the two-conformational-state model, fraction active ($f^0_{active}$) equals $K/(1 + K)$, and the apparent $IC_{50}$ is related to $f^0_{active}$ by:

$$IC_{50} = (\frac{1}{1 - f^0_{active}})K_d$$

To overlay the two-state model prediction with experimental data, the curve was anchored to the WT data point by solving for $K_d$:

$$K_d = IC^{WT}_{50}(1 - f^0_{active,WT})$$

*SHP2 three-conformational-state model with inhibition*

The three-conformational-state model (**Fig. 6A**) was simulated by solving the system of equations:

$K_1 = [E_i]/[E_c]$

$K_2 = [E_o]/[E_i]$

$[I]/K_d = [E_cI]/[E_c]$

$[I]/\alpha K_d = [E_iI]/[E_i]$

$[E_o] + [E_i] + [E_c] + [E_cI] + [E_iI] = 1$

For the simulations in **Fig. 6C**, we used the following parameters:

$K_1 = randomly\ sampled\ from\ a\ log - uniform\ distribution\ between\ 0.011\ and\ 11$

$K_2 = randomly\ sampled\ from\ a\ log - uniform\ distribution\ between\ 0.085\ and\ 85$

$K_d = 24\ nM$

$[I] = 10^{-2+\frac{i}{5}} K_d,\ i = 0, 1, 2, \ldots, 30.$ (31 logarithmically spaced values from 0.01 to 10000)

$r = 0.1$

$\alpha = 0.1$

The rationale for the selection of parameters is detailed in **Supplementary Text 4**.

The exact fractional distribution of all five enzyme states was solved analytically using the SymPy library in Python.

We generated an ensemble of 100,000 theoretical variants, each with a randomly chosen $K_1$ and $K_2$. For each variant and for each $[I]$, the activity is calculated by:

$$Activity([I]) = k \times [E_o] + r \times k \times ([E_i] + [E_iI])$$

To extract the $IC_{50}$ for each simulated variant, the resulting dose-response data points were fitted to a standard single-site inhibition model using the lmfit Python package:

$$Activity([I]) = Top - \frac{(Top - Bottom)[I]}{IC_{50} + [I]}$$

Simulations that yielded increases in activity or lacked valid numerical solutions were excluded.

The intrinsic fraction active was then calculated by:

$$f_{active}^{0} = \frac{K_1K_2 + rK_1}{1 + K_1 + K_1K_2}$$

For parameter sensitivity analysis (**Fig. S60**), we repeated the simulations by independently scaling $r$ and $\alpha$ twofold higher and lower than their baseline values. This resulted in four additional 10,000 variant simulations to evaluate the system dynamics at $r = 0.05$, $r = 0.2$, $\alpha = 0.05$, $\alpha = 0.2$, holding all other baseline parameters constant.

For the simulation of WT dose-response curves in **Fig. 6B** and **Fig. S54B**, we used the following parameters:

$K_1 = 0.11$

$K_2 = 0.85$

$K_d = 24\ nM$

$[I] = 10^{-2+\frac{i}{5}} K_d$, $i = 0, 1, 2, \ldots, 30$. (31 logarithmically spaced values from 0.01 to 10000)

$r = 0.1$

$\alpha = 0.1$

*SHP2 inhibition with simultaneous allosteric activation*

To model inhibition alongside allosteric activation by ligands, a thermodynamic equilibrium model was simulated by solving the following system of equations:

$K_1 = [I]/[C]$

$K_2 = [O]/[I]$

$\frac{[A]}{K_d^{A1}} = \frac{[AI]}{[I]} = \frac{[AO]}{[O]}$

$\frac{[A]}{K_d^{A2}} = \frac{[IA]}{[I]} = \frac{[OA]}{[O]}$

$[OAA] = \frac{[OA]C_{eff}}{K_d^{A1}} + \frac{[AO]C_{eff}}{K_d^{A2}}$

$\frac{[D]}{\alpha K_d} = \frac{[ID]}{[I]} = \frac{[IAD]}{[IA]} = \frac{[AID]}{[AI]}$

$\frac{[D]}{K_d} = \frac{[CD]}{[C]}$

$[C] + [I] + [O] + [AI] + [IA] + [AO] + [OA] + [AOA] + [CD] + [ID] + [AID] + [IAD] = 1$

Here, *C*, *I*, and *O* represent the apo closed, intermediate, and open enzyme states, respectively. States bound to the activator (*A*) are denoted by the addition of *A*. Binding to the nSH2 domain is represented by an *A* in front of the state (e.g., *AI*, *AO*), while binding to the secondary site (e.g., cSH2) is represented by an *A* following the state (e.g., *IA*, *OA*). Because the bisphosphorylated ligand is bivalent, the second binding event occurs intramolecularly and was modeled using an effective concentration ($C_{eff}$) rather than the free ligand concentration ($[A]$). The open state with both nSH2 and cSH2 bound to ligand is denoted as *AOA*. States bound to the inhibitor (*D*) are denoted by the addition of *D* (e.g., *CD*, *ID*, *AID*, *IAD*).

For simulating WT inhibition in the presence of ppIRS1 (**Fig. 6D**), we used the following parameters:

$K_1 = 0.11$

$K_2 = 0.85$

$[D] = 10^{-5+\frac{15i}{99}} K_d$, $i = 0, 1, 2, \ldots, 99$. (100 logarithmically spaced values from $10^{-5}$ to $10^{10}$)

$[A] = 10^{-3+\frac{j}{7}} K_d$, $j = 0, 1, 2, \ldots, 49$. (50 logarithmically spaced values from $10^{-3}$ to $10^{5}$)

$K_d^{A1} = 14\, nM$

$K_d^{A2} = 3400\, nM$

$C_{eff} = 10^5\, nM$

$r = 0.1$

$\alpha = 0.1$

The inhibitor concentrations were varied as multiples of $K_d$. Consequently, the simulated $IC_{50}$ values were expressed as multiples of $K_d$ as well. By expressing concentrations relative to $K_d$, the simulation effectively uses dimensionless values (i.e., $[I]/K_d$). This normalization was chosen to keep the simulations general and independent of an assumed $K_d$, which is not experimentally determined and need not be arbitrarily assigned. The $K_d^{A1}$ and $K_d^{A2}$ values were obtained from prior measurements reported in the literature (*4*). WT $K_1$ and $K_2$ values were chosen based on estimates from published experimental data (*5*).

For each fixed concentration of $[A]$, the activity of the system was simulated across the entire defined range of inhibitor concentrations ($[D]$). The fractional distribution of all twelve enzyme states satisfying the system of equations was solved numerically using the fsolve function from the SciPy library in Python. Activity was calculated by summing the fully active open states and applying a residual activity coefficient ($r = 0.1$) to the intermediate states:

$$Activity([D]) = ([O] + [AO] + [OA] + [AOA]) + r([I] + [AI] + [IA] + [ID] + [AID] + [IAD])$$

To extract the $IC_{50}$ for each simulated condition, dose-response data were fitted to a standard single-site inhibition model using the lmfit Python package:

$$Activity([D]) = Top - \frac{(Top - Bottom)[D]}{IC_{50} + [D]}$$

Two additional parameter sets were simulated: (1) with $K_d^{A1} = \infty$ to simulated R32A mutation with nSH2 binding impaired, and 2) with $K_d^{A2} = \infty$ to simulate R138A mutation with cSH2 binding impaired, while fixing the other parameters (**Fig. 6D**).

*SHP2 inhibition with simultaneous allosteric activation by singly phosphorylated ligands*

To model inhibition alongside allosteric activation by IRS1-pY1172 (**Fig. 6G**), a thermodynamic model was simulated by solving the following system of equations:

$K_1 = [I]/[C]$

$K_2 = [O]/[I]$

$\frac{[A]}{K_d^{A1}} = \frac{[AI]}{[I]} = \frac{[AO]}{[O]}$

$\frac{[A]}{K_d^{A2}} = \frac{[IA]}{[I]} = \frac{[OA]}{[O]}$

$[OAA] = \frac{[OA][A]}{K_d^{A1}} + \frac{[AO][A]}{K_d^{A2}}$

$\frac{[D]}{\alpha K_d} = \frac{[ID]}{[I]} = \frac{[IAD]}{[IA]} = \frac{[AID]}{[AI]}$

$\frac{[D]}{K_d} = \frac{[CD]}{[C]}$

$[C] + [I] + [O] + [AI] + [IA] + [AO] + [OA] + [AOA] + [CD] + [ID] + [AID] + [IAD] = 1$

Here, *C*, *I*, and *O* represent the apo closed, intermediate, and open enzyme states, respectively. States bound to the activator (*A*) are denoted by the addition of *A*. Binding to the nSH2 domain is represented by an *A* in front of the state (e.g., *AI*, *AO*), while binding to the secondary site (e.g., cSH2) is represented by an *A* following the state (e.g., *IA*, *OA*). The fully occupied open state is denoted as *AOA*. States bound to the inhibitor (*D*) are denoted by the addition of *D* (e.g., *CD*, *ID*, *AID*, *IAD*).

For simulating WT inhibition in the presence of IRS1-pY1172, we used the following parameters:

$K_1 = 0.11$

$K_2 = 0.85$

$[D] = 10^{-5+\frac{15i}{99}} K_d,\ i = 0, 1, 2, \dots, 99$. (100 logarithmically spaced values from $10^{-5}$ to $10^{10}$)

$[A] = 10^{-3+\frac{j}{7}} K_d,\ j = 0, 1, 2, \dots, 49$. (50 logarithmically spaced values from $10^{-3}$ to $10^{5}$)

$K_d^{A1} = 14\ nM$

$K_d^{A2} = 3400\ nM$

$r = 0.1$

$\alpha = 0.1$

The inhibitor concentrations were varied as multiples of $K_d$. The $K_d^{A1}$ and $K_d^{A2}$ values were obtained from prior measurements reported in the literature (*4*). WT $K_1$ and $K_2$ values were chosen based on estimates from published experimental data (*5*).

The fractional distribution of all twelve enzyme states satisfying the system of equations was solved numerically using the fsolve function from the SciPy library in Python. Activity was calculated by summing the fully active open states and applying a residual activity coefficient ($r = 0.1$) to the intermediate states:

$$Activity([D]) = ([O] + [AO] + [OA] + [AOA]) + r([I] + [AI] + [IA] + [ID] + [AID] + [IAD])$$

To extract the $IC_{50}$ for each simulated condition, dose-response data were fitted to a standard single-site inhibition model using the lmfit Python package:

$$Activity([D]) = Top - \frac{(Top - Bottom)[D]}{IC_{50} + [D]}$$

To explore variants with altered conformational states (the hypothetical drug sensitive variant in **Fig. 6F**), an additional parameter set was simulated with $K_1 = 10$ and $K_1 = 1$, while fixing the other parameters (**Fig. 6G**).

### Cellular SHP2 inhibition assay

*Cell culture and inhibitor treatment*

U2OS cells were cultured in McCoy's 5A media (ATCC, 30-2007) supplemented with 10% fetal bovine serum (FBS) (GeminiBio, 100-106) and 1% penicillin-streptomycin (Gibco, 15140122). They were maintained at 37°C with 5% carbon dioxide and routinely checked for mycoplasma contamination. Cells were transfected in 24-well dishes using 150 ng of CMV promoter-driven SHP2 plasmid DNA and 0.6 ul FuGENE HD Transfection Reagent (Promega, E2311) per well. After 24 hours, the media was replaced with fresh media containing 0.1% dimethyl sulfoxide (DMSO) and varying concentrations of TNO-155 (MedChem Express, HY-136173), RMC-4630 (MedChem Express, HY-141523), or GDC-1971 (Cayman Chemical Company, 39884). Cells were returned to the incubator for 2 hours before a quick wash with cold DPBS (Corning, 21-031-CV) followed by lysis in 100 ul 1x LDS buffer containing 25 mM dithiothreitol along with protease (Millipore Sigma, 11836170001) and phosphatase (Millipore Sigma, 4906837001) inhibitors.

*Western blotting*

Frozen cell lysates were thawed then boiled at 95°C for 5 minutes, cooled on ice, then sonicated at 15% for 2 pulses of 15 seconds each at 4°C in a Qsonica Q800R instrument. Lysates were loaded on 4-20% polyacrylamide gels (BioRad, 5671095), run in Tris-Glycine SDS buffer (Thermo Fisher Scientific, LC26755), then transferred onto nitrocellulose membranes (Cytiva, 10600001). Membranes were then blocked with 5% bovine serum albumin (BSA) in Tris-buffered saline with 0.25% Tween-20 (TBST) at room temperature for at least 30 min then incubated with primary antibodies against pERK and β-tubulin (Cell Signaling Technology, 9101 and 86298, respectively) diluted 1:1000 in 5% BSA in TBST overnight at 4°C. Membranes were washed 3 times for 5 minutes each in TBST then incubated with anti-mouse IR680 and anti-rabbit IR800 antibodies (LI-COR, 92668070 and 92632213, respectively) diluted 1:20000 in 5% BSA in TBST for 45 minutes at room temperature. After 3 more washes in TBST, membranes were imaged on a LI-COR Odyssey Fc instrument. Membranes were then reprobed with primary antibodies against ERK and SHP2 (Cell Signaling Technology, 9102 and 3397, respectively) in the same manner. Band intensities were quantified using Image Studio Lite software (LI-COR). The pERK signals were divided by their respective β-tubulin signals then each SHP2 variant's DMSO control value was set to 100.

*Curve fitting*

Normalized pERK signals were plotted as a function of inhibitor concentration and fit to a one-site dose-response equation:

$$pERK = Top\frac{IC_{50}}{X + IC_{50}}$$

Each independent experiment was fit separately, and the $IC_{50}$ values reported were calculated from the mean of three independent experiments.

**Statistical analysis**

*General statistical methods.*

Bootstrap resampling, significance testing, and normalization procedures for HT-MEK measurements are described in the **Measuring SHP2 enzyme turnover with HT-MEK** section above. All statistical analyses were performed in Python.

*Identification of substrate-dependent mutational effects*

$k_{cat}/K_M$ for DiFMUP and EGFRpY992 substrates were log-transformed, and a linear regression was fit to the transformed values. Goodness of fit was assessed by $r^2$. Outliers were identified as data points with regression residuals exceeding a *z*-score threshold of 3 (**Fig. 2G**).

*Identification of variants with altered activator affinity*

The product of fraction active and ppIRS1 $EC_{50}$ is predicted to be constant across variants (**Eqn. 1**). A global constant was estimated by computing the mean of $\log_{10}$(fraction active × $EC_{50}$) across all variants. Per-variant deviations from this global constant were assessed using a bootstrap hypothesis test: for each variant, fraction active and $EC_{50}$ measurements were independently resampled with replacement (10,000 iterations), and the resulting null distribution was used to compute a two-tailed p-value. Variants were classified as significantly deviated if they met all three criteria: bootstrap $p < 0.01$, greater than 3-fold deviation from the global fit, and at least 3 replicate measurements (**Fig. 3K**).

*Comparison of biochemical parameters between disease types*

Effect sizes were quantified using Cliff's delta, a non-parametric measure computed as the proportion of all pairwise comparisons between two groups in which group A exceeds group B, minus the proportion in which group B exceeds group A, yielding values ranging from −1 to +1, where $|\delta| = 1$ indicates complete separation. Statistical significance was assessed by a two-sided bootstrap permutation test: group labels were randomly shuffled and the difference in medians recomputed over 10,000 iterations to generate a null distribution, with *p*-values calculated as the fraction of permuted differences equal to or exceeding the observed difference in absolute value. Permutation was performed at the level of individual cases.

*Identification of drug sensitivity outliers*

Median $IC_{50}$ values were $log_{10}$-transformed, and a linear regression was fit to the transformed values across all variants. For each variant with ≥5 replicates per drug, $IC_{50}$ values were independently resampled with replacement (10,000 iterations) and median residuals from the global regression recalculated to generate a null distribution. Two-tailed p-values were corrected using the Benjamini–Hochberg method. Variants were classified as drug-selectivity outliers if they met both criteria: adjusted $p < 0.05$ and ≥3-fold deviation between observed and regression-predicted $IC_{50}$ (**Fig. S52**).

### Supplementary Text

#### S1. Quantifying reversible folding for the catalytic domain of SHP2

#### Measuring reversible folding in the presence of irreversible $SHP2_{CD}$ inactivation on HT-MEK.

When assayed on a plate reader, urea-dependent loss of $SHP2_{CD}$ activity reached equilibrium within 1 min and was largely reversible (>80% activity recovery) even after a 48 hr incubation with urea (**Fig. S14B**). These properties enabled estimation of native-state stability by fitting the urea-dependent activity curve to a two-state folding–denaturation model. Since denaturation was monitored through loss of enzymatic activity, this analysis reports the apparent stability of the catalytically competent native ensemble relative to inactive, non-native conformations, rather than a strictly defined native–unfolded structural transition. The fitted m-value of 1.38 kcal/mol/M (95% CI: 1.36–1.40 kcal/mol/M) was lower than the theoretical value (3.5±1.6 kcal/mol/M), consistent with the presence of partially unfolded intermediates (**Fig. S71**). While non-linearities in linear extrapolation can bias stability estimates in some cases with partially unfolded intermediates, such effects are expected to be minimal here given that SHP2 is marginally stable and the extrapolation distance is short. Here, we acknowledge these complications by reporting values as the apparent native state stability ($\Delta G_{native}^{app}$).

On HT-MEK devices, urea-dependent loss of activity did not reach an apparent equilibrium and prolonged incubation with 1 M urea led to continued loss of activity over time (**Fig. S72A**). Furthermore, SHP2 variants only partially recovered activity upon transfer from urea-containing to urea-free conditions on HT-MEK (**Fig. S72A**). These observations suggest that some fraction of SHP2 molecules undergo irreversible inactivation under denaturing conditions within HT-MEK devices, likely due to interactions between unfolded protein and the device surface. Notably, SHP2 variants do not exhibit loss of activity under native conditions over the typical time required to perform experiments (≤8 hr) (**Fig. S72B**), indicating that irreversible inactivation arises specifically from the unfolded state.

To account for these processes and enable high-throughput recovery of thermodynamic stability information via HT-MEK, we developed a kinetic model describing these processes, where: (1) $E_F$ and $E_U$ represent the folded and unfolded states of the enzyme, respectively, (2) $k_F$ is the folding rate, (3) $k_U$ is the unfolding rate, and (4) $k_{decay}$ is the rate of irreversible inactivation (**Fig. S72C**).

Reversible folding kinetics can be estimated from plate reader assays. Since reactions require $3\tau$ in time to reach > 95% completion and the urea-dependent loss of SHP2 activity reaches equilibrium within 1 min (**Fig. S14B&S72H**), we estimate $3\tau_{relax} < 1\,min$. Thus, $\tau_{relax} < 1/3\,min$, and the relaxation rate $k_{relax} = k_F + k_U > 3\,min^{-1}$. From plate reader assays, the equilibrium native fraction at 1M urea is $f_F^{eq} = 0.61\ \pm 0.03$, allowing the estimation of the parameters $k_F$ and $k_U$:

$$k_F = k_{relax} \times f_F^{eq} > 1.83\,min^{-1}$$

$$k_U = k_{relax} \times (1 - f_F^{eq}) > 1.17\,min^{-1}$$

Irreversible inactivation kinetics were estimated by fitting a single exponential equation, $y = Ae^{-kt}$, to the loss of $SHP2_{CD}$ WT activity at 1M urea over time on HT-MEK (**Fig. S72D**), yielding $k_{obs} = 0.003 \pm 0.001\,min^{-1}$. Since $k_{obs} = k_{decay} \times f_U$, we estimate $k_{decay} \approx k_{obs}/(1 - f_F^{eq}) = 0.008\,min^{-1}$.

Assuming that reversible unfolding kinetics are similar on the plate reader and on-chip, we concluded that $k_F + k_U \gg k_{decay}$. Therefore, the initial drop in activity observed upon urea addition on HT-MEK primarily reflects reversible unfolding, with only a minor contribution from slow irreversible inactivation. To illustrate this, we simulated the time-dependent loss of SHP2 activity in the presence of 1M urea using a two-state unfolding model with irreversible decay. The kinetics are governed by the following ordinary differential equations:

$$\frac{dE_f}{dt} = -k_u E_f + k_f E_u$$

$$\frac{dE_u}{dt} = k_u E_f - k_f E_u - k_{decay} E_u$$

$$\frac{d\emptyset}{dt} = k_{decay} E_u$$

Simulation from an initial condition of $E_F = 1$ and $E_U = 0$ show that $E_F$ rapidly decays to the equilibrium folded fraction $f_F^{eq}$, as expected, followed by a slower decay due to irreversible inactivation (**Fig. S72E**). Consequently, measuring the initial rate of SHP2 activity in the presence of urea primarily reports the effects of reversible unfolding. In our assay, measuring SHP2 activity for 30 min is expected to yield native fraction estimates within 10% of the true value based on comparing the model-simulated $E_F$ at 30 min and $f_F^{eq}$. Additionally, by resetting the baseline using activity measurements in the absence of urea (i.e., interspersing 0 M urea baseline measurements between each incremental step in urea concentration), the remaining total enzyme ($E_{tot}$) can be reassessed after each urea assay. This approach allows reliable measurement of $f_U^{eq}$ repeatedly within the same chamber of HT-MEK devices, even in the presence of irreversible inactivation. As an example, the residual activity observed at 1 M urea relative to the prior 0 M urea measurement remains stable over multiple iterations and agrees with results from the plate reader assay for both $SHP2_{CD}$ and $SHP2_{FL}$ when processed in this way (**Fig. S72F&G**).

**Estimating folding stability of the catalytic PTP domain of $SHP2_{FL}$**

Since SHP2 is a multidomain protein, the addition of urea can perturb multiple equilibria, not just a single folding transition. At minimum, we expect four equilibria to be perturbed:

1. Folding of the PTP domain.
2. Folding of the nSH2 domain.
3. Folding of the cSH2 domain.
4. The autoinhibition interaction in which SH2 domains contact the PTP domain.

Thus, while the urea-dependent activity of $SHP2_{FL}$ fits well to a two-state model and is both reversible and stable over time (**Fig. S72H&I**), the curve is unlikely to reflect a meaningful thermodynamic quantity for SHP2. To identify mutations that likely impact $SHP2_{FL}$ thermodynamic stability, we quantified activity loss at 1 M urea, a concentration at which prior literature measurements suggest the SH2 domains remain folded but our $SHP2_{CD}$ measurements reveal destabilizes the PTP domain (*6*). We also expect the SH2–PTP autoinhibitory interaction to be minimally perturbed at 1 M urea, as its denaturant sensitivity (m-value) should be much smaller due to the relatively modest change in buried hydrophobic surface area upon dissociation.

To estimate the native fraction of the catalytic PTP domain of $SHP2_{FL}$ variants in the absence of denaturant (0 M urea), we used the measured native fraction at 1 M together with a fixed m-value of 1.4 kcal mol. This m-value was determined from $SHP2_{CD}$ variants (**Fig. S71**) and is expected to be unchanged, since it reflects the intrinsic folding properties of the same catalytic domain regardless of whether it is expressed in isolation or within the full-length protein.

The fraction native at a given urea concentration is given by the standard two-state relationship:

$$f_F^{eq}([urea]) = \frac{1}{1 + \exp\left(-\frac{\Delta G^0 - m[urea]}{RT}\right)}$$

Where $\Delta G^0$ is the folding energy in the absence of urea, $m$ is the denaturant dependence, and $RT$ is the thermal energy.

Rearranging the equation at 1 M urea allows $\Delta G^0$ to be expressed in terms of the measured $f_F^{eq}(1)$:

$$\Delta G^0 = m + RT ln\left(\frac{f_F^{eq}(1)}{1 - f_F^{eq}(1)}\right)$$

Substituting this expression into the equation for 0 M urea yields:

$$f_F^{eq}(0) = \frac{1}{1 + \frac{1 - f_F^{eq}(1)}{f_F^{eq}(1)} e^{-m/RT}}$$

### S2. Derivation of the relationship between conformational equilibrium constants and sensitivity to allosteric activator and inhibitor.

#### Derivation of $EC_{50}$ for SHP2 allosteric activator that stabilizes the active state

To describe the effect of an allosteric activator that selectively binds the open (active) conformation of SHP2, we consider a two-state conformational equilibrium between a closed, inactive state and the open state (**Fig. S73**).

The equilibrium constant for this conformational transition is defined as:

$$K = \frac{[E_o]}{[E_c]}$$

The activator binds exclusively to the open state with the dissociation constant $K_d^A$ defined as:

$$K_d^A = \frac{[E_o][A]}{[E_oA]}$$

In these two equations, $[E_o]$ is the concentration of free enzyme in the open state, $[E_c]$ is the concentration of free enzyme in the closed state, $[E_oA]$ is the concentration of enzyme in the activator bound state, and $[A]$ is the activator concentration. Thus, the total enzyme population comprises $[E_c]$, $[E_o]$, and the activator-bound species $[E_oA]$, with only the latter two contributing to observed activity with rate constant $k$.

In all derivations, we made the following assumptions: (1) SHP2 conformational change and activator binding is at equilibrium, (2) substrate is subsaturating and so does not influence activator binding or cause a change in the SHP2 conformational equilibrium, and (3) catalytic activity is proportional to the total population in the active states.

This framework allows us to derive an expression for the observed activity as a function of activator concentration, and from it, the relationship between the half-maximal activating concentration $EC_{50}$, the dissociation constant $K_d^A$, and the conformational equilibrium constant $K$.

The total enzyme is:

$$[E_{tot}] = [E_c] + [E_o] + [E_oA]$$

From the equilibria, we can write everything in terms of $[E_o]$:

$[E_c] = \frac{[E_o]}{K}$;      $[E_oA] = \frac{[E_o][A]}{K_d^A}$

Thus,

$$[E_{tot}] = [E_o]\left(\frac{1}{K} + 1 + \frac{[A]}{K_d^A}\right).$$

Only the open species $[E_o]$ and $[E_oA]$ are catalytically competent, so the fraction active is:

$$f_{active}([A]) = \frac{[E_o] + [E_oA]}{[E_{tot}]} = \frac{1 + \frac{[A]}{K_d^A}}{\frac{1}{K} + 1 + \frac{[A]}{K_d^A}},$$

and the intrinsic fraction open is:

$$f_{active}^0 = \frac{[E_o]}{[E_o] + [E_c]} = \frac{1}{\frac{1}{K} + 1} = \frac{K}{1 + K}.$$

The observed activity is proportional to the active fraction:

$$k_{obs}([A]) = f_{active}([A]) \times k = \frac{1 + \frac{[A]}{K_d^A}}{\frac{1}{K} + 1 + \frac{[A]}{K_d^A}} \times k$$

Define $EC_{50}$ as the activator concentration where the activity is halfway between baseline and saturation:

$$k_{obs}(EC_{50}) = \frac{k_{obs}(0) + \lim_{[A]\to\infty} k_{obs}([A])}{2}$$

Since baseline (no activator) activity is

$$k_{obs}(0) = f_{active}^0 \times k = \frac{K}{1 + K} \times k,$$

maximal (saturating activator) activity is

$$\lim_{[A]\to\infty} k_{obs}([A]) = k,$$

And the activity halfway between baseline and saturation is:

$$\frac{k_{obs}(0) + \lim_{[A]\to\infty} k_{obs}([A])}{2} = \frac{\frac{K}{1 + K} \times k + k}{2} = \frac{(1 + 2K)k}{2(1 + K)}$$

The activity with activator concentration equal to $EC_{50}$ is:

$$k_{obs}(EC_{50}) = \frac{1 + \frac{EC_{50}}{K_d^A}}{\frac{1}{K} + 1 + \frac{EC_{50}}{K_d^A}} \times k = \frac{K + K\frac{EC_{50}}{K_d^A}}{1 + K + K\frac{EC_{50}}{K_d^A}} \times k$$

Plugging in these values to solve for the value of $EC_{50}$ that satisfies $k_{obs}(EC_{50}) = (k_{obs}(0) + \lim_{[A]\to\infty} k_{obs}([A]))/2$ gives:

$$\frac{K + K\frac{EC_{50}}{K_d^A}}{1 + K + K\frac{EC_{50}}{K_d^A}} \times k = \frac{(1 + 2K)k}{2(1 + K)}$$

$$\Rightarrow \frac{K + K\frac{EC_{50}}{K_d^A}}{1 + K + K\frac{EC_{50}}{K_d^A}} = \frac{(1 + 2K)}{2(1 + K)}$$

$$\Rightarrow 1 - \frac{1}{1 + K + K\frac{EC_{50}}{K_d^A}} = 1 - \frac{1}{2(1+K)}$$

$$\Rightarrow \frac{1}{1 + K + K\frac{EC_{50}}{K_d^A}} = \frac{1}{2(1+K)}$$

$$\Rightarrow 1 + K + K\frac{EC_{50}}{K_d^A} = 2 + 2K$$

$$\Rightarrow K\frac{EC_{50}}{K_d^A} = 1 + K$$

$$\Rightarrow EC_{50} = \frac{1+K}{K}K_d^A$$

Since $f_{active}^0 = K/(1+K)$, the result is equivalently:

$$EC_{50} = \frac{K_d^A}{f_{active}^0}$$

**Derivation of IC$_{50}$ for an allosteric inhibitor stabilizing the closed state**

We now consider the case where a small-molecule inhibitor (I) binds selectively to the closed state of the enzyme (**Fig. S74A**).

The equilibrium constant for the conformational transition is defined as:

$$K = \frac{[E_o]}{[E_c]}$$

The inhibitor binds exclusively to the closed state, and the dissociation constant $K_d^I$ is defined as:

$$K_d^I = \frac{[E_c][I]}{[E_cI]}$$

Where $[E_o]$ is the concentration of enzyme in the open state, $[E_c]$ is the concentration of enzyme in the closed state, $[E_cI]$ is the concentration of enzyme in the inhibitor bound state, and $[I]$ is the inhibitor concentration.

To derive the relationship between $IC_{50}$, the dissociation constant $K_d^I$, and the conformational equilibrium constant $K$, we made the following assumptions: (1) SHP2 conformational change and inhibitor binding is at equilibrium, (2) catalysis is under subsaturating substrate conditions such that catalysis does not influence activator binding or cause a change in the SHP2 conformational equilibrium, and (3) catalytic activity is proportional to the population in active state.

The total enzyme is:

$$[E_{tot}] = [E_c] + [E_o] + [E_cI]$$

From the equilibria, we can write everything in terms of $[E_c]$:

$[E_o] = K[E_c]$;  $[E_cI] = \frac{[E_c][I]}{K_d^I}$

Thus,

$$[E_{tot}] = [E_c]\left(1 + K + \frac{[I]}{K_d^I}\right)$$

Only $[E_o]$ is catalytically competent, so the fraction active is

$$f_{active}([A]) = \frac{[E_o]}{[E_{tot}]} = \frac{K}{1 + K + \frac{[I]}{K_d^I}}$$

and the intrinsic fraction open is

$$f_{active}^0 = \frac{[E_o]}{[E_o] + [E_c]} = \frac{1}{\frac{1}{K} + 1} = \frac{K}{1 + K}$$

The observed activity is proportional to the active fraction:

$$k_{obs}([I]) = f_{active}([I]) \times k = \frac{K}{1 + K + \frac{[I]}{K_d^I}} \times k$$

Define $IC_{50}$ as the inhibitor concentration where the activity is halfway between baseline and saturation:

$$k_{obs}(IC_{50}) = \frac{k_{obs}(0) + \lim_{[I]\to\infty} k_{obs}([I])}{2}$$

Since baseline (no inhibitor) activity is

$$k_{obs}(0) = f_{active}^0 \times k = \frac{K}{1 + K} \times k,$$

and activity at saturating inhibitor is

$$\lim_{[I]\to\infty} k_{obs}([I]) = 0,$$

the activity halfway between baseline and saturation is:

$$\frac{k_{obs}(0) + \lim_{[I]\to\infty} k_{obs}([I])}{2} = \frac{\frac{K}{1 + K} \times k}{2} = \frac{K \times k}{2(1 + K)}$$

The activity with inhibitor concentration equal to $IC_{50}$ is:

$$k_{obs}(IC_{50}) = \frac{K}{1 + K + \frac{IC_{50}}{K_d^I}} \times k$$

Plugging in these values to solve for the value of $IC_{50}$ that satisfies $k_{obs}(IC_{50}) = (k_{obs}(0) + \lim_{[I]\to\infty} k_{obs}([I]))/2$ gives:

$$\frac{K}{1 + K + \frac{IC_{50}}{K_d^I}} \times k = \frac{K \times k}{2(1 + K)}$$

$$\Rightarrow \frac{1}{1 + K + \frac{IC_{50}}{K_d^I}} = \frac{1}{2(1 + K)}$$

$$\Rightarrow 1 + K + \frac{IC_{50}}{K_d^I} = 2 + 2K$$

$$\Rightarrow \frac{IC_{50}}{K_d^I} = 1 + K$$

$$\Rightarrow IC_{50} = (1 + K)K_d^I$$

Since

$$f_{active}^0 = \frac{K}{1 + K}$$

and

$$1 - f_{active}^0 = 1 - \frac{K}{1 + K} = \frac{1}{1 + K}$$

the result is equivalent to:

$$IC_{50} = \frac{K_d^I}{1 - f_{active}^0}$$

**S3. Assessing alternative models of SHP2 conformational states to explain inhibition data.**

The prevailing model for SHP2 inhibition (**Fig. S74A**), hereafter referred to as *Model 1*, is based on the following observations. First, only two SHP2 conformational states have been crystallized to date: a closed state and an open state. In the closed state, the catalytic site is occluded, whereas in the open state the active site is accessible. Second, the crystal structure of SHP2 bound to an allosteric inhibitor is nearly identical to the closed-state structure. Together, these observations support the simplest mechanistic model *(Model 1)*: SHP2 exists in an equilibrium between a closed/inactive and an open/active conformation, and the inhibitor binds selectively to the closed state, thereby stabilizing the inactive form.

However, *Model 1* fails to explain two key features of the experimental data presented here: (i) inhibition remains incomplete even at saturating inhibitor concentrations, and (ii) many variants with increased open-state propensity exhibit greater inhibitor sensitivity.

To identify the minimal model capable of reproducing these features, we systematically tested progressively more complex alternative models. Each model was evaluated based on whether simulations recapitulated (i) incomplete inhibition at saturating inhibitor concentration and (ii) the observed relationship between $IC_{50}$ and basal fraction active.

In all simulations, we made the following assumptions: (1) drug binding and SHP2 conformational change is at equilibrium, (2) catalysis does not influence drug binding or the SHP2 conformational change equilibrium, and (3) catalytic activity is proportional to the fraction of the enzyme population in active states. All simulations were performed using Python scripts. To generalize the results, all simulations were carried out in dimensionless units.

*Model 1*

We first simulated *Model 1* to compare with the analytical solutions (**Supplementary text S2**). The concentration of each species was simulated by solving for $[E_o]$, $[E_c]$ , and $[E_cI]$ from the following equations:

$K = [E_o]/[E_c]$

$[I]/K_d = [E_cI]/[E_c]$

$[E_o] + [E_c] + [E_cI] = 1$

Where $[E_o]$ is the fraction of enzyme in the open state, $[E_c]$ is the fraction of enzyme in the closed state, $[E_cI]$ is the fraction of enzyme in the inhibitor bound state, and $[I]$ is the inhibitor concentration.

Using the following parameters:

$K = 10^{-2+\frac{i}{5}}, i = 0, 1, 2, \ldots, 20$. (21 logarithmically spaced values from 0.01 to 100)

$[I] = 10^{-2+\frac{i}{5}} K_d, i = 0, 1, 2, \ldots, 30$. (31 logarithmically spaced values from 0.01 to 10000)

The total enzyme concentration was normalized to unity, such that each species is expressed as a fraction of the total enzyme population. If desired, these values can be converted to absolute concentrations by multiplying by the total enzyme concentration.

The inhibitor concentrations were varied as multiples of $K_d$. Consequently, the simulated $IC_{50}$ values were expressed as multiples of $K_d$ as well. By expressing concentrations relative to $K_d$, the simulation effectively uses dimensionless values (i.e., $[I]/K_d$). This normalization was chosen to keep the simulations general and independent of an assumed $K_d$, which is not experimentally determined and need not be arbitrarily assigned.

After solving $[E_o]$, $[E_c]$ , and $[E_cI]$ at a range of inhibitor concentrations $[I]$ for a given value of K, we calculated the enzyme activity at each condition as:

$$Activity = k \times [E_o]$$

where $k$ is the rate constant. The resulting activity versus $[I]$ data were then fitted to a standard inhibitory dose–response function to extract the $IC_{50}$ for that given K:

$$f(I) = Top - (Top - Bkg) \times \frac{[I]}{[I] + IC_{50}}$$

To visualize how $IC_{50}$ depends on differences in enzyme conformational equilibrium (i.e., *K* values altered by mutations), $IC_{50}$ was plotted as a function of the fraction of open enzyme in the absence of inhibitor, calculated as:

$$f^0_{active} = \frac{[E_o]}{[E_o] + [E_c]} = \frac{K}{1 + K}$$

This allowed us to compare inhibitor sensitivity across different intrinsic enzyme conformational equilibria (i.e., different $K$values). To provide a reference point, we labeled K=0.1 as WT, consistent with experimentally determined estimates. Simulations of *Model 1* show complete inhibition at saturating inhibitor concentrations, consistent with analytical solutions (**Fig. S74B**). Furthermore, the $IC_{50}$ plotted as a function of the fraction of active enzyme in the absence of inhibitor exactly matched the analytically derived relationship (**Fig. S74C**), confirming the validity of the simulation. However, both observations are inconsistent with experimental data, confirming that *Model 1* fails to capture key features of the system.

**Two-conformational-state models**

We next examined modifications of *Model 1* without introducing additional conformational states. The assumptions of Model 1 are: (1) The closed state is completely inactive, and (2) the inhibitor binds exclusively to the closed state. We systematically relaxed these assumptions in *Model 1*.

*Model 1-1*

In *Model 1-1*, the closed state was allowed to retain partial catalytic activity (**Fig. S74D**). Although this assumption is structurally inconsistent with the closed-state crystal structure, where the active site is occluded, we still explored this possibility for completeness. The rationale was to be rigorous and to account for the possibility of mechanisms that are not yet understood.

The concentration of $[E_o]$, $[E_c]$ , and $[E_cI]$ is determined by the following system of equations:

$K = [E_o]/[E_c]$

$[I]/K_d = [E_cI]/[E_c]$

$[E_o] + [E_c] + [E_cI] = 1$

Where we normalize the total enzyme concentration to 1.

To simplify, we express everything in $[E_c]$:

$[E_o] = K[E_c]$

$[E_cI] = \frac{[I]}{K_d}[E_c]$

Substitute these into the mass balance equation to solve for $[E_c]$:

$$[E_o] + [E_c] + [E_cI] = 1$$

$$\Rightarrow [E_c](K + 1 + \frac{[I]}{K_d}) = 1$$

$$\Rightarrow [E_c] = \frac{1}{K + 1 + \frac{[I]}{K_d}}$$

The activity can then be expressed as:

$$k_{obs}([I]) = k([E_o] + r([E_c] + [E_cI])) = k[E_c](K + r\left(1 + \frac{[I]}{K_d}\right)) = k\frac{K + r\left(1 + \frac{[I]}{K_d}\right)}{K + 1 + \frac{[I]}{K_d}}$$

Here, *r* represents the fraction of catalytic activity retained by the enzyme in the closed state.

To determine if the enzyme retains activity at saturating inhibitor concentrations, we evaluate the limit of $k_{obs}([I])$ as $[I]$ approaches infinity.

$$\lim_{[I]\to\infty} k_{obs}([I]) = \lim_{[I]\to\infty} k\frac{K + r\left(1 + \frac{[I]}{K_d}\right)}{K + 1 + \frac{[I]}{K_d}} = \lim_{[I]\to\infty} \frac{rk\frac{[I]}{K_d}}{\frac{[I]}{K_d}} = rk$$

Thus, in *Model 1-1*, there will be residual activity at saturating inhibitor concentrations.

To determine if variants with higher $f^0_{active}$ can yield a lower $IC_{50}$, we must examine whether $f^0_{active}$ and $IC_{50}$ can move inversely with respect to $K$. We examine the partial derivatives of both functions in the domain $K > 0$ and $r \in [0,1]$.

Note that in this model, the closed state is also catalytically competent, so the true intrinsic fraction open does not equal the experimental measurable $f^0_{active}$. Rather, $f^0_{active}$ represents the proportion of catalytic capacity realized at basal conditions compared to the maximal possible activity:

$$f^0_{active} = \frac{[E_o] + r * [E_c]}{[E_c] + [E_o]} = \frac{K + r}{K + 1}$$

Since the direct experimental measurable is $f^0_{active}$, we analyzed the relationship between $f^0_{active}$ and $IC_{50}$ for consistency.

Define $IC_{50}$ as the inhibitor concentration where the activity is halfway between baseline and saturation:

$$k_{obs}(IC_{50}) = \frac{k_{obs}(0) + \lim_{[I]\to\infty} k_{obs}([I])}{2} = \frac{k\frac{K + r}{K + 1} + rk}{2}$$

Solving for $IC_{50}$ yields:

$$IC_{50} = K_d(K+1)$$

We now examine how $IC_{50}$ and $f^0_{active}$ move with respect to the equilibrium constant $K$. The derivatives are:

$$\frac{\partial f^0_{active}}{\partial K} = \frac{(K+1)-(K+r)}{(K+1)^2} = \frac{1-r}{(K+1)^2} > 0$$

$$\frac{\partial IC_{50}}{\partial K} = K_d > 0$$

Since both $f^0_{active}$ and $IC_{50}$ are strictly increasing functions of $K$, a variant with higher $f^0_{active}$ will strictly result in a higher $IC_{50}$ according to _Model 1-1_.

We simulated _Model 1-1_ by solving for $[E_o]$, $[E_c]$ , and $[E_cI]$ from the following system of equations with the provided parameters:

$K = [E_o]/[E_c]$

$[I]/K_d = [E_cI]/[E_c]$

$[E_o] + [E_c] + [E_cI] = 1$

$K = 10^{-2+\frac{i}{5}}, i = 0, 1, 2, \ldots, 20.$ (21 logarithmically spaced values from 0.01 to 100)

$[I] = 10^{-2+\frac{i}{5}} K_d, i = 0, 1, 2, \ldots, 30.$ (31 logarithmically spaced values from 0.01 to 10000)

$r = 0.1, 0.5, 0.9.$

After solving $[E_o]$, $[E_c]$ , and $[E_cI]$ at a range of inhibitor concentrations $[I]$ for a given value of $K$, we calculated the enzyme activity at each condition as:

$$Activity = k \times [E_o] + r \times k \times ([E_cI] + [E_c])$$

The intrinsic fraction active is:

$$f^0_{active} = \frac{[E_o] + r * [E_c]}{[E_c] + [E_o]} = \frac{K+r}{K+1}$$

Simulations across three different potential values of *r* (*r* = 0.1, 0.5, and 0.9) revealed that Model 1-1 produces incomplete inhibition but fails to reproduce the lower $IC_{50}$ values observed for more “open” (higher $f^0_{active}$) variants (**Fig. S74E-S74J**).

_Model 1-2_

Here, the inhibitor was permitted to bind the open state with affinity $\alpha K_d$ (α > 1) (**Fig. S74K**). For α ≤ 1, the ligand would not act as an inhibitor. This introduces a new species $[E_oI]$. The concentrations of $[E_o]$, $[E_c]$ , $[E_cI]$, and $[E_oI]$ are determined by the following system of equations:

$K = [E_o]/[E_c]$

$[I]/K_d = [E_cI]/[E_c]$

$[I]/\alpha K_d = [E_oI]/[E_o]$

$[E_o] + [E_c] + [E_cI] + [E_oI] = 1$

Where we normalize the total enzyme concentration to 1.

To simplify, we express everything in $[E_c]$:

$[E_o] = K[E_c]$

$[E_cI] = \frac{[I]}{K_d}[E_c]$

$[E_oI] = \frac{K[I]}{\alpha K_d}[E_c]$

Substitute these into the mass balance equation to solve for $[E_c]$:

$$[E_o] + [E_c] + [E_cI] + [E_oI] = 1$$

$$\Rightarrow [E_c](K + 1 + \frac{[I]}{K_d} + \frac{K[I]}{\alpha K_d}) = 1$$

$$\Rightarrow [E_c] = \frac{1}{K + 1 + \frac{[I]}{K_d} + \frac{K[I]}{\alpha K_d}}$$

The activity can then be expressed as:

$$k_{obs}([I]) = k([E_o] + [E_oI]) = k[E_c](K + \frac{K[I]}{\alpha K_d}) = k\frac{K + \frac{K[I]}{\alpha K_d}}{K + 1 + \frac{[I]}{K_d} + \frac{K[I]}{\alpha K_d}}$$

To determine if the enzyme retains activity at saturating inhibitor concentrations, we evaluate the limit of $k_{obs}([I])$ as $[I]$ approaches infinity.

$$\lim_{[I]\to\infty} k_{obs}([I]) = \lim_{[I]\to\infty} k\frac{\frac{K[I]}{\alpha K_d}}{\frac{[I]}{K_d} + \frac{K[I]}{\alpha K_d}} = \frac{kK}{\alpha + K}$$

Thus, in *Model 1-2*, there will be residual activity at saturating inhibitor concentrations.

To determine if variants with higher $f_{active}^0$ can yield a lower $IC_{50}$, we must examine whether $f_{active}^0$ and $IC_{50}$ can move inversely with respect to $K$. We examine the partial derivatives of both functions in the domain $K > 0$ and $\alpha \in [0,1]$.

In this model, the intrinsic fraction active is:

$$f_{active}^0 = \frac{[E_o]}{[E_c] + [E_o]} = \frac{K}{K + 1}$$

Define $IC_{50}$ as the inhibitor concentration where the activity is halfway between baseline and saturation:

$$k_{obs}(IC_{50}) = \frac{k_{obs}(0) + \lim_{[I]\to\infty} k_{obs}([I])}{2} = \frac{\frac{kK}{1 + K} + \frac{kK}{\alpha + K}}{2}$$

Solving for $IC_{50}$ yields:

$$IC_{50} = K_d\frac{\alpha(1 + K)}{\alpha + K}$$

We now examine how $IC_{50}$ and $f_{active}^0$ move with respect to the equilibrium constant $K$. The derivatives are:

$$\frac{\partial f_{active}^0}{\partial K} = \frac{1}{(K + 1)^2} > 0$$

$$\frac{\partial IC_{50}}{\partial K} = K_d \frac{\alpha(\alpha-1)}{(\alpha+K)^2} > 0$$

Since both $f_{active}^{0}$ and $IC_{50}$ are strictly increasing functions of $K$, a variant with higher $f_{active}^{0}$ will strictly result in a higher $IC_{50}$ according to *Model 1-2*.

We simulated *Model 1-2* by solving for $[E_o]$, $[E_c]$ , $[E_cI]$, and $[E_oI]$ from the following system of equations with the provided parameters:

$K = [E_o]/[E_c]$

$[I]/K_d = [E_cI]/[E_c]$

$[I]/\alpha K_d = [E_oI]/[E_o]$

$[E_o] + [E_c] + [E_cI] + [E_oI] = 1$

$K = 10^{-2+\frac{i}{5}}$, $i = 0, 1, 2, \ldots, 20$. (21 logarithmically spaced values from 0.01 to 100)

$[I] = 10^{-2+\frac{i}{5}} K_d$, $i = 0, 1, 2, \ldots, 30$. (31 logarithmically spaced values from 0.01 to 10000)

$\alpha = 10, 100, 1000$

After solving $[E_o]$, $[E_c]$ , and $[E_cI]$ at a range of inhibitor concentrations $[I]$ for a given value of $K$ and $\alpha$, the enzyme activity at each condition was calculated as:

$$Activity = k \times ([E_o] + [E_oI])$$

The intrinsic fraction active was then calculated by:

$$f_{active}^{0} = \frac{[E_o]}{[E_o] + [E_c]} = \frac{K}{1+K}$$

Simulations of *Model 1-2* with multiple α values (α = 2, 10, or 100) also generated incomplete inhibition but, as with *Model 1-1*, did not capture the experimentally observed shift in $IC_{50}$ for more open variants (**Figs. S74L-S74Q**).

**Three-state models**

Since all possible variations of the two-state framework failed to explain the experimental observations, we next considered the simplest higher-order alternative: a three-state model. In this framework, SHP2 interconverts between a closed, an intermediate, and an open conformation. Two transition schemes are possible: (1) a sequential pathway (closed ↔ intermediate ↔ open), or (2) free interconversion among all three states. Since our analysis is based on equilibrium thermodynamics, the two schemes are thermodynamically equivalent. For simplicity and based on prior single-molecule studies, we adopted the sequential model for further analysis. To provide a reference point, we estimated the relative population of the three states for WT based on literature data.

*Model 2-1*

The simplest version of the three-state model assumes that the inhibitor binds exclusively to the closed state (**Fig. S75A**). In *Model 2-1*, $[E_o]$ is the fraction of enzyme in the open state, $[E_i]$ is the fraction of enzyme in the intermediate state, $[E_c]$ is the fraction of enzyme in the closed state, $[E_cI]$ is the fraction of enzyme in the inhibitor bound state, and $[I]$ is the inhibitor concentration. *r* represents the fractional activity of the intermediate state relative to the open state. The concentration of each species is determined by the following system of equations:

$K_1 = [E_i]/[E_c]$

$K_2 = [E_o]/[E_i]$

$[I]/K_d = [E_cI]/[E_c]$

$[E_o] + [E_i] + [E_c] + [E_cI] = 1$

Where we normalize the total enzyme concentration to 1.

To simplify, we express everything in $[E_c]$:

$[E_i] = K_1[E_c]$

$[E_o] = K_2[E_i] = K_1K_2[E_c]$

$[E_cI] = \frac{[I]}{K_d}[E_c]$

Substitute these into the mass balance equation to solve for $[E_c]$:

$$[E_o] + [E_i] + [E_c] + [E_cI] = 1$$

$$\Rightarrow [E_c](K_1K_2 + K_1 + 1 + \frac{[I]}{K_d}) = 1$$

$$\Rightarrow [E_c] = \frac{1}{K_1K_2 + K_1 + 1 + \frac{[I]}{K_d}}$$

The activity can then be expressed as:

$$k_{obs}([I]) = k([E_o] + r[E_i]) = k[E_c](K_1K_2 + rK_1) = k\frac{K_1K_2 + rK_1}{K_1K_2 + K_1 + 1 + \frac{[I]}{K_d}}$$

To determine if the enzyme retains activity at saturating inhibitor concentrations, we evaluate the limit of $k_{obs}([I])$ as $[I]$ approaches infinity.

$$\lim_{[I]\to\infty} k_{obs}([I]) = \lim_{[I]\to\infty} \frac{K_1K_2 + rK_1}{K_1K_2 + K_1 + 1 + \frac{[I]}{K_d}} = 0$$

Thus, in _Model 2-1_, there will be no residual activity at saturating inhibitor concentrations.

To determine if a variant with higher $f_{active}^0$ can yield a lower $IC_{50}$, we must examine whether $f_{active}^0$ and $IC_{50}$ can move inversely with respect to the underlying parameters $K_1$ and $K_2$. We examine the partial derivatives of both functions in the domain $K_1$, $K_2 > 0$ and $r \in [0,1]$.

The intrinsic fraction active is:

$$f_{active}^0 = \frac{k([E_o] + r[E_i])}{k([E_o] + [E_i] + [E_c])} = \frac{K_1K_2 + rK_1}{K_1K_2 + K_1 + 1}$$

Define $IC_{50}$ as the inhibitor concentration where the activity is halfway between baseline and saturation:

$$k_{obs}(IC_{50}) = \frac{k_{obs}(0) + \lim_{[I]\to\infty} k_{obs}([I])}{2} = \frac{k(K_1K_2 + rK_1)}{2(K_1K_2 + K_1 + 1 + \frac{[I]}{K_d})}$$

Solving for $IC_{50}$ yields:

$$IC_{50} = K_d(K_1K_2 + K_1 + 1)$$

We analyzed the gradient of $f^0_{active}$ and $IC_{50}$ with respect to $K_1$, $K_2$:

$$\frac{\partial f^0_{active}}{\partial K_1} = \frac{(K_2 + r)(K_1K_2 + K_1 + 1) - (K_2 + 1)(K_1K_2 + rK_1)}{(K_1K_2 + K_1 + 1)^2} = \frac{(K_2 + r)}{(K_1K_2 + K_1 + 1)^2} > 0$$

$$\frac{\partial f^0_{active}}{\partial K_2} = \frac{K_1(K_1K_2 + K_1 + 1) - K_1(K_1K_2 + rK_1)}{(K_1K_2 + K_1 + 1)^2} = \frac{K_1(1 + K_1(1 - r))}{(K_1K_2 + K_1 + 1)^2} > 0$$

Gradients of $IC_{50}$:

$$\frac{\partial IC_{50}}{\partial K_1} = K_d(K_2 + 1) > 0$$

$$\frac{\partial IC_{50}}{\partial K_2} = K_dK_1 > 0$$

Since both $f^0_{active}$ and $IC_{50}$ are strictly increasing functions of both $K_1$ and $K_2$, there is no domain where altering $K_1$ or $K_2$ independently increases $f^0_{active}$ while decreasing $IC_{50}$. While an isolated variant could theoretically exhibit a higher $f^0_{active}$ and lower $IC_{50}$ via precisely balanced, opposing shifts in $K_1$ and $K_2$ (e.g., a decrease in $K_2$ offset by a large increase in $K_1$), *Model 2-1* cannot produce a dominant inverse correlation between $f^0_{active}$ and $IC_{50}$ across a general sampling of the parameter space.

We simulated *Model 2-1* under conditions in which the intermediate state was assigned activities ranging from completely inactive to equivalent to the open state. The concentration of each species was simulated by solving for $[E_o]$, $[E_i]$, $[E_c]$ and $[E_cI]$ from the following system of equations:

$K_1 = [E_i]/[E_c]$

$K_2 = [E_o]/[E_i]$

$[I]/K_d = [E_cI]/[E_c]$

$[E_o] + [E_i] + [E_c] + [E_cI] = 1$

Where $[E_o]$ is the fraction of enzyme in the open state, $[E_i]$ is the fraction of enzyme in the intermediate state, $[E_c]$ is the fraction of enzyme in the closed state, $[E_cI]$ is the fraction of enzyme in the inhibitor bound state, and $[I]$ is the inhibitor concentration.

The equations were solved with the following parameters:

$K_1 = 0.01\ to\ 100$

$K_2 = 0.01\ to\ 100$

$[I] = 10^{-2+\frac{i}{5}} K_d,\ i = 0, 1, 2, \ldots, 30$. (31 logarithmically spaced values from 0.01 to 10000)

$r = 0, 0.1, 1$

As a reference, we defined the WT condition as $K_1 = 0.11$ and $K_2 = 0.85$. These values were chosen based on estimates from published experimental data (*5*).

As expected, *Model 2-1* was unable to recapitulate the incomplete inhibition or the experimentally observed shift in $IC_{50}$ for more open variants across a variety of potential *r* values (**Figs. S75B**&**C**).

*Model 2-2*

In this variant of the three-state model, the inhibitor binds exclusively to the intermediate state (**Fig. S75D**). The species therefore include $[E_o]$, $[E_i]$, $[E_c]$ and the inhibitor bound intermediate complex $[E_iI]$, and $r$ represents the fractional activity of the intermediate state relative to the open state. The concentration of each species is determined by the following system of equations:

$K_1 = [E_i]/[E_c]$

$K_2 = [E_o]/[E_i]$

$[I]/K_d = [E_iI]/[E_i]$

$[E_o] + [E_i] + [E_c] + [E_iI] = 1$

Expressing everything in $[E_c]$:

$[E_i] = K_1[E_c]$

$[E_o] = K_2[E_i] = K_1K_2[E_c]$

$[E_iI] = \frac{[I]}{K_d}[E_i] = \frac{[I]}{K_d}K_1[E_c]$

Substitute these into the mass balance equation to solve for $[E_c]$:

$$[E_o] + [E_i] + [E_c] + [E_iI] = 1$$

$$\Rightarrow [E_c](K_1K_2 + K_1 + 1 + \frac{[I]}{K_d}K_1) = 1$$

$$\Rightarrow [E_c] = \frac{1}{K_1K_2 + K_1 + 1 + \frac{[I]}{K_d}K_1}$$

The activity can then be expressed as:

$$k_{obs}([I]) = k([E_o] + r([E_i] + [E_iI]) = k[E_c]\left(K_1K_2 + rK_1 + r\frac{[I]}{K_d}K_1\right) = k\frac{K_1K_2 + rK_1 + r\frac{[I]}{K_d}K_1}{K_1K_2 + K_1 + 1 + \frac{[I]}{K_d}K_1}$$

To determine if the enzyme retains activity at saturating inhibitor concentrations, we evaluate the limit of $k_{obs}([I])$ as $[I]$ approaches infinity.

$$\lim_{[I]\to\infty} k_{obs}([I]) = \lim_{[I]\to\infty} k\frac{K_1K_2 + rK_1 + r\frac{[I]}{K_d}K_1}{K_1K_2 + K_1 + 1 + \frac{[I]}{K_d}K_1} = \lim_{[I]\to\infty} k\frac{r\frac{[I]}{K_d}K_1}{\frac{[I]}{K_d}K_1} = rk$$

Thus, in _Model 2-2_, there will be residual activity at saturating inhibitor concentrations.

While we do not have a direct measurement of $r$, we can estimate the bounds of $r$ under this model. To physically function as an inhibitor rather than an activator, the compound must suppress the baseline activity of the enzyme. Mathematically, this dictates that the basal activity in the absence of the inhibitor must be strictly greater than the residual activity at saturating inhibitor concentrations:

$$k_{obs}(0) > \lim_{[I]\to\infty} k_{obs}([I])$$

$$\Longrightarrow k\frac{K_1K_2 + rK_1}{K_1K_2 + K_1 + 1} > rk$$

$$\Rightarrow r < \frac{K_1K_2}{K_1K_2+1}$$

This inequality defines the upper bound for the fractional activity of the intermediate state $r$. Applying this boundary to the WT condition, constrained by the experimentally estimated equilibrium constants $K_1 = 0.11$ and $K_2 = 0.85$:

$$r < \frac{0.11 \times 0.85}{0.11 \times 0.85 + 1} \approx 0.086$$

To determine if a variant with higher $f^0_{active}$ can yield a lower $IC_{50}$, we must examine whether $f^0_{active}$ and $IC_{50}$ can move inversely with respect to the underlying parameters $K_1$ and $K_2$. We examine the partial derivatives of both functions in the domain $K_1, K_2 > 0$ and $r \in [0,0.086]$.

The intrinsic fraction active $f^0_{active}$ is:

$$f^0_{active} = \frac{k(r[E_i]+[E_o])}{k([E_o]+[E_i]+[E_c])} = \frac{K_1K_2 + rK_1}{K_1K_2 + K_1 + 1}$$

Define $IC_{50}$ as the inhibitor concentration where the activity is halfway between baseline and saturation:

$$k_{obs}(IC_{50}) = \frac{k_{obs}(0) + \lim_{[I]\to\infty} k_{obs}([I])}{2} = \frac{k(K_1K_2 + rK_1) + rk(K_1K_2 + K_1 + 1)}{2(K_1K_2 + K_1 + 1)}$$

Solving for $IC_{50}$ yields:

$$IC_{50} = K_d(K_2 + 1 + \frac{1}{K_1})$$

We analyzed the gradient of $f^0_{active}$ and $IC_{50}$ with respect to $K_1$, $K_2$:

$$\frac{\partial f^0_{active}}{\partial K_1} = \frac{(K_2+r)(K_1K_2+K_1+1) - (K_2+1)(K_1K_2+rK_1)}{(K_1K_2+K_1+1)^2} = \frac{(K_2+r)}{(K_1K_2+K_1+1)^2} > 0$$

$$\frac{\partial f^0_{active}}{\partial K_2} = \frac{K_1(K_1K_2+K_1+1) - K_1(K_1K_2+rK_1)}{(K_1K_2+K_1+1)^2} = \frac{K_1(1+K_1(1-r))}{(K_1K_2+K_1+1)^2} > 0$$

$$\frac{\partial IC_{50}}{\partial K_1} = -K_d(1/K_1)^2 < 0$$

$$\frac{\partial IC_{50}}{\partial K_2} = K_d > 0$$

Since $\frac{\partial f^0_{active}}{\partial K_1}$ and $\frac{\partial IC_{50}}{\partial K_1}$ have opposite signs, if a variant has a higher $f^0_{active}$ specifically due to an increase in $K_1$ (destabilizing the closed state relative to the intermediate state), the model dictates that the $IC_{50}$ will decrease. Thus, _Model 2-2_ is expected to produce a population-level inverse correlation between $f^0_{active}$ and $IC_{50}$ under random sampling of the conformational landscape.

We simulated _Model 2-2_. The species concentrations were obtained by solving the system of equations:

$K_1 = [E_i]/[E_c]$

$K_2 = [E_o]/[E_i]$

$[I]/K_d = [E_iI]/[E_i]$

$[E_o] + [E_i] + [E_c] + [E_iI] = 1$

The equations were solved with the following parameters:

$K_1 = random\ from\ 0.01\ to\ 100$

$K_2 = random\ from\ 0.01\ to\ 100$

$[I] = 10^{-2+\frac{i}{5}} K_d,\ i = 0, 1, 2, \ldots, 30$. (31 logarithmically spaced values from 0.01 to 10000)

$r = 0, 0.01, 0.05$

After solving $[E_o]$, $[E_i]$ , $[E_c]$, and $[E_i I]$ at a range of inhibitor concentrations $[I]$ for a given value of $K_1$, $K_2$ and r, we calculated the enzyme activity at each condition as:

$$Activity = k \times [E_o] + r \times k \times ([E_i] + [E_i I])$$

and

$$f^0_{active} = \frac{[E_o] + r * [E_i]}{[E_c] + [E_i] + [E_o]} = \frac{K_1 K_2 + r K_1}{1 + K_1 + K_1 K_2}$$

Simulations of *Model 2-2* demonstrated that when $r > 0$, the model reproduced incomplete inhibition (**Fig. S75E**). In addition, conditions with a lower intrinsic fraction closed exhibited lower $IC_{50}$ values (**Fig. S75F**), matching experimental observations. Thus, *Model 2-2* represents the minimal model capable of recapitulating the experimental data.

*Model 2-3*

Although *Model 2-2* is the minimal model that could recapitulate the experimental results, the assumption that the drug only binds to the intermediate state is inconsistent with the crystal structure, which showed that the inhibitor binds to the closed state. To reconcile this discrepancy, we amended the model by allowing the drug to bind to the closed state as well (**Fig. S76A**). The species therefore include $[E_o]$, $[E_i]$, $[E_c]$ and the inhibitor bound complexes $[E_c I]$ and $[E_i I]$. This also introduces an additional parameter α, which determines the relative affinity of $[E_i]$, $[E_c]$ towards the inhibitor. The species concentrations were obtained by solving the system of equations:

$K_1 = [E_i]/[E_c]$

$K_2 = [E_o]/[E_i]$

$[I]/K_d = [E_c I]/[E_c]$

$[I]/\alpha K_d = [E_i I]/[E_i]$

$[E_o] + [E_i] + [E_c] + [E_c I] + [E_i I] = 1$

Expressing everything in $[E_c]$:

$[E_i] = K_1 [E_c]$

$[E_o] = K_2 [E_i] = K_1 K_2 [E_c]$

$[E_c I] = \frac{[I]}{K_d} [E_c]$

$[E_i I] = \frac{[I]}{\alpha K_d} [E_i] = \frac{K_1 [I]}{\alpha K_d} [E_c]$

Substitute these into the mass balance equation to solve for $[E_c]$:

$$[E_o] + [E_i] + [E_c] + [E_c I] + [E_i I] = 1$$

$$\Rightarrow [E_c](K_1K_2 + K_1 + 1 + \frac{[I]}{K_d} + \frac{K_1[I]}{\alpha K_d}) = 1$$

$$\Rightarrow [E_c] = \frac{1}{K_1K_2 + K_1 + 1 + \frac{[I]}{K_d} + \frac{K_1[I]}{\alpha K_d}}$$

The activity can then be expressed as:

$$k_{obs}([I]) = k([E_o] + r([E_i] + [E_iI]) = k\frac{K_1K_2 + rK_1 + r\frac{K_1[I]}{\alpha K_d}}{K_1K_2 + K_1 + 1 + \frac{[I]}{K_d} + \frac{K_1[I]}{\alpha K_d}}$$

To determine if the enzyme retains activity at saturating inhibitor concentrations, we evaluate the limit of $k_{obs}([I])$ as $[I]$ approaches infinity.

$$\lim_{[I]\to\infty} k_{obs}([I]) = \lim_{[I]\to\infty} k\frac{K_1K_2 + rK_1 + r\frac{K_1[I]}{\alpha K_d}}{K_1K_2 + K_1 + 1 + \frac{[I]}{K_d} + \frac{K_1[I]}{\alpha K_d}} = \lim_{[I]\to\infty} k\frac{r\frac{K_1[I]}{\alpha K_d}}{\frac{[I]}{K_d} + \frac{K_1[I]}{\alpha K_d}} = k\frac{rK_1}{\alpha + K_1}$$

Thus, in *Model 2-3*, there will be residual activity at saturating inhibitor concentrations. Note that if $\alpha \to \infty$, the limit goes to 0, reverting to *Model 2-1*; if $\alpha \to 0$, the limit goes to $rk$, reverting to *Model 2-2.*

To determine if a variant with higher $f^0_{active}$ can yield a lower $IC_{50}$, we must examine whether $f^0_{active}$ and $IC_{50}$ can move inversely with respect to the underlying parameters $K_1$ and $K_2$. We examine the partial derivatives of both functions in the domain $K_1$, $K_2 > 0$, $\alpha > 0$, and $r > 0$.

The intrinsic fraction active $f^0_{active}$ is:

$$f^0_{active} = \frac{k(r[E_i] + [E_o])}{k([E_o] + [E_i] + [E_c])} = \frac{K_1K_2 + rK_1}{K_1K_2 + K_1 + 1}$$

Define $IC_{50}$ as the inhibitor concentration where the activity is halfway between baseline and saturation:

$$k_{obs}(IC_{50}) = \frac{k_{obs}(0) + \lim_{[I]\to\infty} k_{obs}([I])}{2} = \frac{k\frac{K_1K_2 + rK_1}{K_1K_2 + K_1 + 1} + k\frac{rK_1}{\alpha + K_1}}{2}$$

Solving for $IC_{50}$ yields:

$$IC_{50} = K_d\frac{K_1K_2 + K_1 + 1}{1 + \frac{K_1}{\alpha}}$$

We analyzed the gradient of $f^0_{active}$ and $IC_{50}$ with respect to $K_1$, $K_2$:

$$\frac{\partial f^0_{active}}{\partial K_1} = \frac{(K_2 + r)(K_1K_2 + K_1 + 1) - (K_2 + 1)(K_1K_2 + rK_1)}{(K_1K_2 + K_1 + 1)^2} = \frac{(K_2 + r)}{(K_1K_2 + K_1 + 1)^2} > 0$$

$$\frac{\partial f^0_{active}}{\partial K_2} = \frac{K_1(K_1K_2 + K_1 + 1) - K_1(K_1K_2 + rK_1)}{(K_1K_2 + K_1 + 1)^2} = \frac{K_1(1 + K_1(1 - r))}{(K_1K_2 + K_1 + 1)^2} > 0$$

$$\frac{\partial IC_{50}}{\partial K_1} = K_d\frac{(1 + K_2)\left(1 + \frac{K_1}{\alpha}\right) - (K_1K_2 + K_1 + 1)\frac{1}{\alpha}}{(1 + \frac{K_1}{\alpha})^2} = K_d\frac{1 + K_2 - \frac{1}{\alpha}}{(1 + \frac{K_1}{\alpha})^2}$$

$$\frac{\partial IC_{50}}{\partial K_2} = K_d \frac{K_1}{1 + \frac{K_1}{\alpha}} > 0$$

Since $\frac{\partial f_{active}^0}{\partial K_1}$, $\frac{\partial f_{active}^0}{\partial K_2}$, and $\frac{\partial IC_{50}}{\partial K_2}$ are strictly positive, generating a population-level inverse correlation between $f_{active}^0$ and $IC_{50}$ requires $\frac{\partial IC_{50}}{\partial K_1}$ to be negative. This occurs when:

$$1 + K_2 - \frac{1}{\alpha} < 0$$

$$\Rightarrow \alpha < \frac{1}{1 + K_2}$$

Because $K_2 > 0$, this requires $\alpha < 1$ for *Model 2-3* to recapitulate the experimental data.

We simulated *Model 2-3*. The species concentrations were obtained by solving the system of equations with the following parameters:

$K_1 = random\ from\ 0.01\ to\ 100$

$K_2 = random\ from\ 0.01\ to\ 100$

$[I] = 10^{-2+\frac{i}{5}} K_d$, $i = 0, 1, 2, \ldots, 30$. (31 logarithmically spaced values from 0.01 to 10000)

$r = 0.1$

$\alpha = 0.1, 1, 10$

After solving $[E_o]$, $[E_i]$ , $[E_c]$, $[E_iI]$ and $[E_cI]$ at a range of inhibitor concentrations $[I]$ for a given value of $K_1$, $K_2$, r, and α, the enzyme activity at each condition was calculated as:

$$Activity = k \times [E_o] + r \times k \times ([E_i] + [E_iI])$$

The intrinsic fraction active was then calculated by:

$$f_{active}^0 = \frac{[E_o] + r * [E_i]}{[E_c] + [E_i] + [E_o]} = \frac{K_1K_2 + rK_1}{1 + K_1 + K_1K_2}$$

Simulations of *Model 2-3* showed that when $\alpha = 0.1$ the model reproduced both incomplete inhibition and the reduced $IC_{50}$ values observed for conditions with a lower intrinsic fraction closed **(Fig. S76B&S76C)**. Thus, *Model 2-3* is capable of recapitulating the experimental results provided that drug binding is stronger to the intermediate state than to the closed state.

**S4. Rationale for parameter selection for simulating *Model 2-3***

As direct biochemical measurements of the fractional activity of the transient intermediate state ($r$) and the microscopic relative binding affinity ($\alpha$) are not currently accessible, we mathematically define a biologically plausible parameter space to simulate *Model 2-3* (**Fig. S77**). Their permissible values are bounded by the following theoretical and empirical constraints:

1. **The inverse $f_{active}^0$ to $IC_{50}$ relationship**

As derived in the *Model 2-3* analysis, for a variant with higher $f_{active}^0$ to yield a lower $IC_{50}$, the system must satisfy the inequality $\alpha < 1$ **(Fig. S77)**.

2. **The inhibition versus activator phase boundary**

To function biologically as an inhibitor, a compound must suppress the baseline activity of the enzyme. This implies that the activity in the absence of the drug, $k_{obs}(0)$, must be strictly greater than the residual activity at saturating drug concentrations, $\lim_{[I]\to\infty} k_{obs}([I])$. We establish the inequality for inhibition:

$$k_{obs}(0) > \lim_{[I]\to\infty} k_{obs}([I])$$

First, we substitute the exact expressions previously derived for *Model 2-3* for both the basal activity and the infinite-concentration limit:

$$k\frac{K_1K_2 + rK_1}{K_1K_2 + K_1 + 1} > k\frac{rK_1}{\alpha + K_1}$$

Which can be subsequently simplified to isolate $\alpha$:

$$\alpha > \frac{r(1 + K_1K_2) - K_1K_2}{K_2 + r}$$

Substituting the WT enzyme parameters ($K_1 = 0.11$ and $K_2 = 0.85$) into this inequality yields:

$$\alpha > \frac{1.1r - 0.094}{0.85 + r}$$

For the compound to act as an inhibitor on the WT enzyme, the selected parameters must reside in this defined phase regime where $\alpha$ must be above a critical boundary $\alpha_c = \frac{1.1r-0.094}{0.85+r}$ **(Fig. S77)**.

#### 3. Empirical WT incomplete inhibition plateau

The TNO155 dose-response data demonstrated an incomplete inhibition plateau for SHP2 variants. Across 24 experimental replicates, the mean normalized residual fractional activity for SHP2 WT at saturating TNO155 concentrations ($P$) was $0.566 \pm 0.077$.

The theoretical residual fractional activity $P$ is defined by the model as the ratio of the saturated limit to the basal activity:

$$P = \frac{\lim_{[I]\to\infty} k_{obs}([I])}{k_{obs}(0)} = \frac{r(K_1K_2 + K_1 + 1)}{(\alpha + K_1)(r + K_2)}$$

Solving for $\alpha$ establishes a direct equality curve linking $\alpha$ and $r$ based on this experimentally observed plateau:

$$\alpha = \frac{r(K_1K_2 + K_1 + 1)}{P(r + K_2)} - K_1$$

Substituting the WT enzyme parameter ($K_1 = 0.11$ and $K_2 = 0.85$) into the equation gives:

$$\alpha = \frac{r(1.2)}{P(r + 0.85)} - 0.11$$

The experimental uncertainty in the normalized inhibition dictates that the true physiological plateau lies within a statistical band, where $P \in [0.489, 0.643]$.

Substituting these bounds yields the boundary curves for the permissible parameter **(Fig. S77)**.

#### 4. Empirical maximum IC$_{50}$ sensitization limit

Analytical derivation of the $IC_{50}$ under *Model 2-3* reveals that the shift in $IC_{50}$ for any variant is governed by the relative binding affinity ($\alpha$), the equilibrium constants ($K_1$, $K_2$), and $K_d$.

The maximum theoretical fold-decrease in $IC_{50}$ relative to WT ($FC_{max}$) purely caused by conformational equilibrium change (achieved as a mutant approaches complete occupancy of the intermediate state, $K_1 \rightarrow \infty$, $K_2 \rightarrow 0$) is defined by:

$$FC_{max} = \frac{IC_{50}^{WT}}{IC_{50}^{min}}$$

$$\Rightarrow FC_{max} = \frac{K_d \dfrac{K_1^{WT} K_2^{WT} + K_1^{WT} + 1}{1 + \dfrac{K_1^{WT}}{\alpha}}}{\lim\limits_{K_1 \rightarrow \infty, K_2 \rightarrow 0} K_d \dfrac{K_1 K_2 + K_1 + 1}{1 + \dfrac{K_1}{\alpha}}}$$

$$\Rightarrow FC_{max} = \frac{K_d \dfrac{K_1^{WT} K_2^{WT} + K_1^{WT} + 1}{1 + \dfrac{K_1^{WT}}{\alpha}}}{\alpha K_d}$$

$$\Rightarrow FC_{max} = \frac{1 + K_1^{WT} + K_1^{WT} K_2^{WT}}{\alpha + K_1^{WT}}$$

Inspection of the empirical data reveals that the predominant cluster of highly active variants converges on an approximate 4-fold sensitization relative to WT. To physically accommodate this representative 4-fold shift, the mathematical ceiling of the model must be greater than or equal to this observed value. Substituting the WT equilibrium constants ($K_1 = 0.11$ and $K_2 = 0.85$) into the inequality establishes an upper limit for $\alpha$:

$$4 = \frac{1 + 0.11 + 0.11 \times 0.85}{\alpha + 0.11}$$

$$\Rightarrow \alpha < 0.19$$

Thus, to recapitulate the empirically observed $IC_{50}$ shifts, the inhibitor must bind the intermediate state tightly enough that $\alpha < 0.19$ **(Fig. S77)**.

#### 5. Empirical basal fraction active limit

The $f_{active}^0$ of the WT enzyme was experimentally estimated at approximately 7.3%. The theoretical basal fraction active is defined by the model as:

$$f_{active}^0 = \frac{K_1 K_2 + r K_1}{K_1 K_2 + K_1 + 1}$$

Applying the established WT equilibrium constants derived from previous single molecule studies ($K_1 = 0.11$ and $K_2 = 0.85$) dictates a theoretical minimum for baseline activity (*5*). If the intermediate state is assumed to be completely inactive ($r = 0$), the equilibrium population of the fully open state alone provides a $f_{active}^0$ of roughly 7.8%.

Attempting to analytically solve for $r$ using our 7.3% empirical $f_{active}^0$ measurement alongside these single-molecule-derived constants yields a small, mathematically impossible negative value. This slight discrepancy likely stems from minor experimental variance in either the single-molecule-derived equilibrium constants ($K_1$, $K_2$) or the bulk fraction active assay ($f_{active}^0$). However, because the measured 7.3% activity is close to the 7.8% theoretical minimum, the empirical data rests right at the boundary of the physically possible regime. To approach the experimental measurement as closely as physically allowed, the intermediate state must possess low activity. Therefore, we can conclude that $r$ is small.

**Selection of parameters**

Because the parameters $r$ and $\alpha$ cannot be directly measured experimentally, we sought to define a best-estimate parameter set that faithfully represents the system. Guided by the conclusion that $r$ must be small to satisfy the basal activity limits (Constraint 5), we selected $r = 0.1$ as a pragmatic estimate for our simulations. Note that selecting $r = 0.1$ leads to a theoretical $f_{active}^{0}$ of 8.7%, representing an 18.9% relative difference from empirically measured $f_{active}^{0}$ at 7.3%, within reasonable experimental error margins. We then paired this with $\alpha = 0.1$, which satisfies all established boundaries (**Fig. S77**). Therefore, while $r$ and $\alpha$ cannot be directly measured, the numbers selected here ($r = 0.1$, $\alpha = 0.1$) represents our most rigorous attempt to establish a plausible starting point, strictly confined by both the model's theoretical limits and the empirical data. Using these fixed parameters alongside the established WT equilibrium constants ($K_1 = 0.11$ and $K_2 = 0.85$), we calculated the dissociation constant to the closed state as $K_d = 24\,nM$ from the experimentally measured TNO155 $IC_{50}^{WT}$. This $K_d$ was subsequently used for simulations for all other variants.

The underlying distribution for $K_1$ and $K_2$ for SHP2 variants is unknown. However, experimental data indicates that the WT exhibits one of the lower $f_{active}^{0}$ values among the variants. Since increases in $K_1$ and $K_2$ directly lead to higher $f_{active}^{0}$, we chose to sample $K_1$ and $K_2$ values 10-fold lower to 100-fold higher than WT $K_1$ and $K_2$. This range accommodates $f_{active}^{0}$ values up to ~99%, while positioning the WT near the lower end of $f_{active}^{0}$ spectrum.

**S5. Rationale for the possibility of having multiple intermediate states.**

Prior single molecule data show that binding of phosphotyrosine peptide binding to one SH2 domain, either only the nSH2 or only the cSH2, favors the intermediate state (*5*). According to this model, the baseline prediction assumes additive effects such that binding to both should favor the intermediate state as well. That is, if each single binding stabilizes the intermediate by $\Delta G_I$ and the open state by $\Delta G_o$ ($\Delta G_I < \Delta G_o$, favoring *I*), dual binding would stabilize $2 \times \Delta G_I \ll 2 \times \Delta G_o$, making the intermediate dominant. However, binding to both SH2 domains favors the open state. This non-additivity indicates allosteric coupling between the SH2 domains.

The simplest model that explains this observation is negative coupling within the intermediate state: simultaneous binding of nSH2 and cSH2 disfavors the intermediate and instead stabilizes the open state. For any allosteric coupling to occur, the protein must access more than two conformational states. Thus, the intermediate state is not a single conformation but an ensemble of sub-states. In its simplest form, this ensemble comprises two distinct conformations: one compatible only with nSH2 binding, one compatible only with cSH2 binding, and neither accommodating both bindings simultaneously. Under this formalism, binding of both SH2 domains will not be compatible with the intermediate state, and higher concentration of the activator will push the equilibrium towards the open state.

### Supplementary Figures

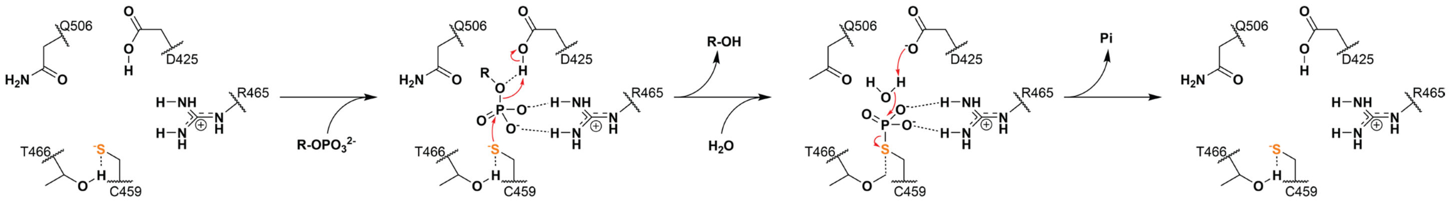


**Figure S1. SHP2 catalytic mechanism and key residues.**

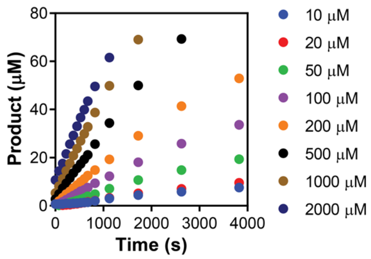


**Figure S2. Example of HT-MEK measured DiFMUP hydrolysis progress curves for SHP2 WT**. Imaging settings were optimized to accurately resolve product at <100 µM formation; above ~100 µM the camera pixels are saturated.

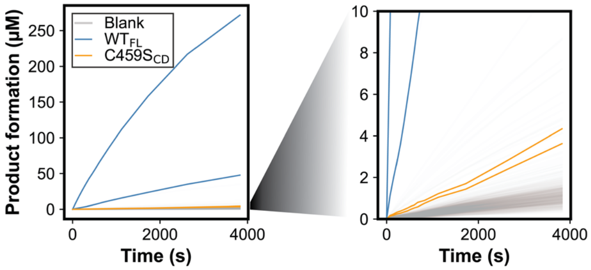


**Figure S3. Representative time-course data of product formation for WT, C459S$_{CD}$, and empty chambers (Blank) measured simultaneously on the same device at 1000 μM substrate concentration.** Two WT traces have enzyme concentrations of 14.0 and 1.3 nM, respectively, and two C459S$_{CD}$ traces have enzyme concentrations of 13.9 and 12.2 nM, respectively.

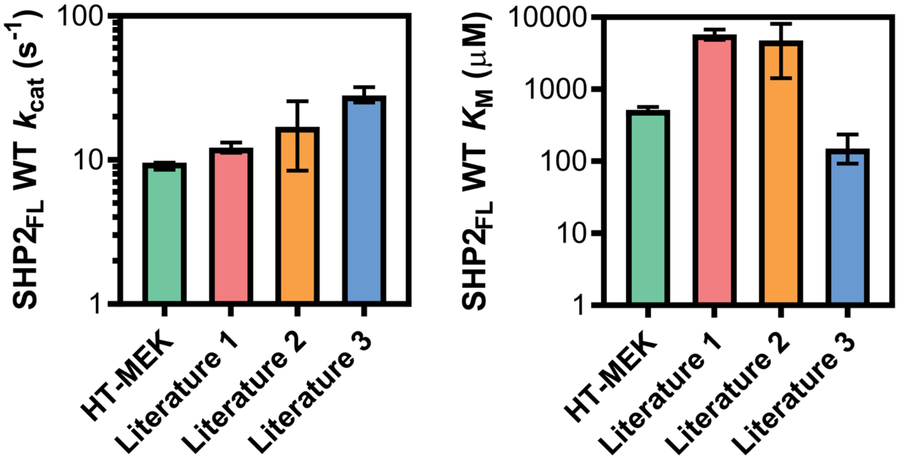


**Figure S4. Comparison of HT-MEK measured Michaelis-Menten parameters for DiFMUP hydrolysis with prior literature reports** (*7*–*9*)**.** HT-MEK uses SHP2(1–525)-EGFP; Literature 1 and 2 use untagged SHP2(1–525); and Literature 3 uses SHP2(1–525) fused to a C-terminal V5 epitope tag. Error bars represent 95% confidence intervals. The wide range likely originates from differences in assay conditions and the specific enzyme constructs utilized.

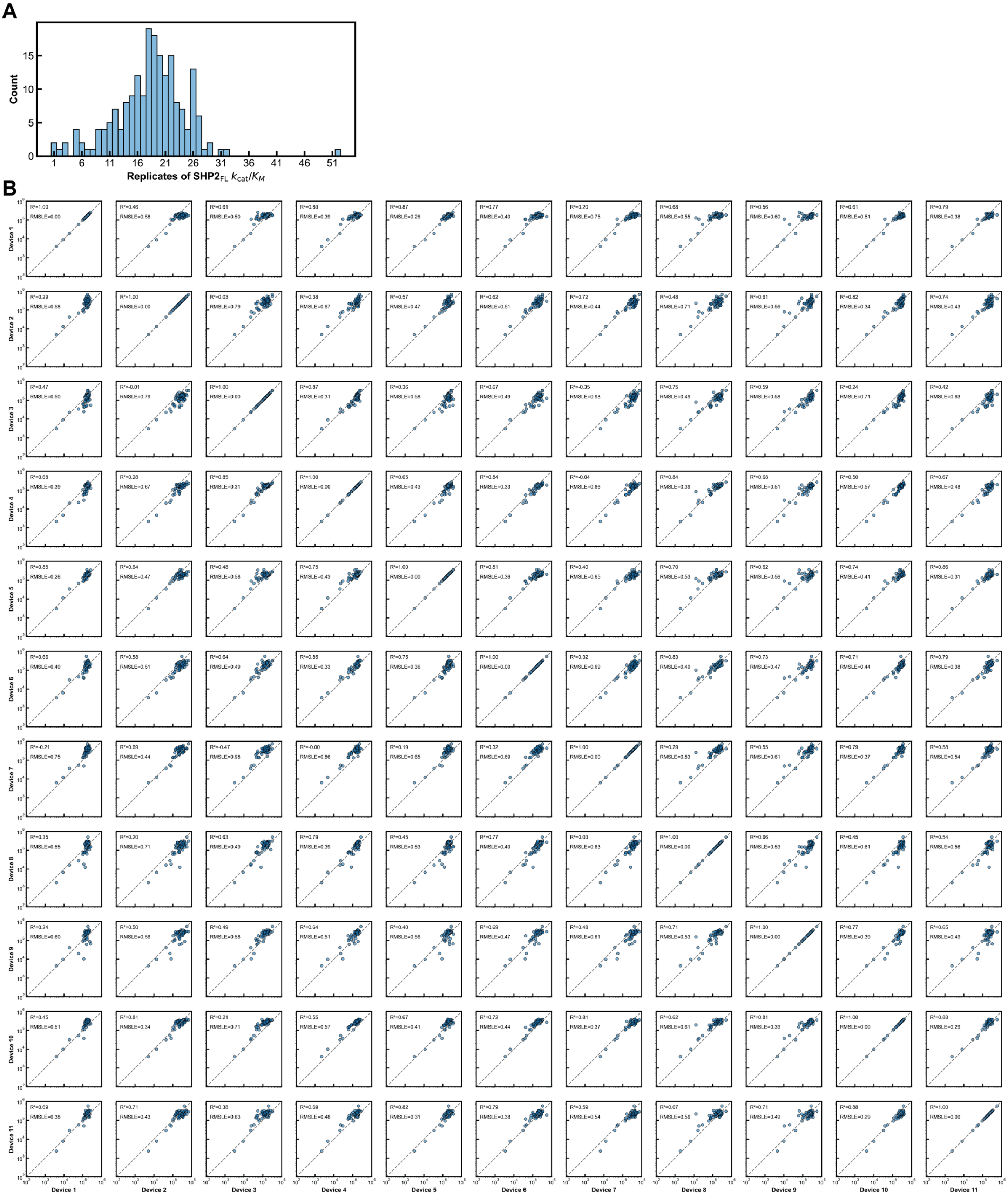

**Figure S5. Reproducibility of $SHP2_{FL}$ DiFMUP hydrolysis activity measurements. (A)** Histogram of number of Michaelis-Menten parameter measurement replicates for 190 $SHP2_{FL}$ variants. **(B)** Reproducibility of $k_{cat}/K_M$ for DiFMUP hydrolysis across device replicates. Each device reports a consensus value derived from 1–4 technical replicates measured in different chambers for the same variant on the same chip. For variants with a single technical replicate, the single measured value is reported; for those with two replicates, the mean is reported; and for variants with three or more replicates, the median is reported. All values are in $M^{-1}s^{-1}$.

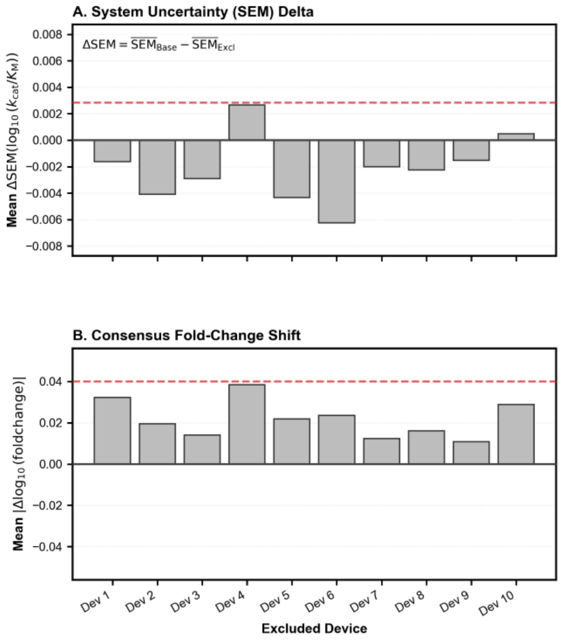


**Figure S6. Leave-one-device-out quality-control analysis of SHP2$_{FL}$ $k_{cat}/K_M$ measurements. (A)** Change in system uncertainty, quantified as the difference in the mean standard error of the mean (SEM) of $\log_{10}(k_{cat}/K_M)$ across variants when each device is excluded in turn relative to the global baseline including all devices. Positive values indicate reduced uncertainty upon exclusion of the device. **(B)** Stability of consensus measurements assessed by the mean absolute shift across all variants in $\log_{10}$(fold change) (difference in $\log_{10}(k_{cat}/K_M)$ relative to an internal anchor variant) compared to the global consensus upon exclusion of each device. The anchor was defined as the variant present on all devices with the highest total number of replicate measurements. Larger values indicate greater deviation from the global consensus upon exclusion of the device. Dashed lines indicate mean + 2 SD thresholds. Because no single device exceeded the threshold for both metrics, all devices were used in the final analysis.

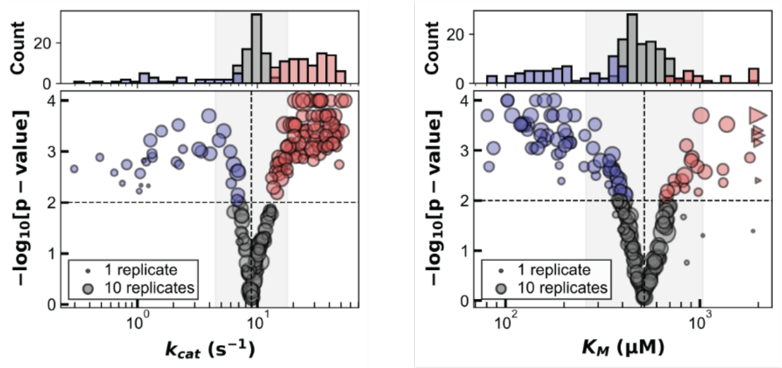


**Figure S7. Median $k_{cat}$ and $K_M$ for DiFMUP hydrolysis across 190 SHP2$_{FL}$ variants**. Triangles indicate mutants with fitted $K_M$ above the highest substrate concentration used; these values are set to the highest substrate concentration used (2000 μM) and represent a lower limit. Blue and red markers indicate variants with a statistically significant decrease or increase compared to WT, respectively ($p < 0.01$). The gray-shaded background denotes a two-fold change range relative to WT.

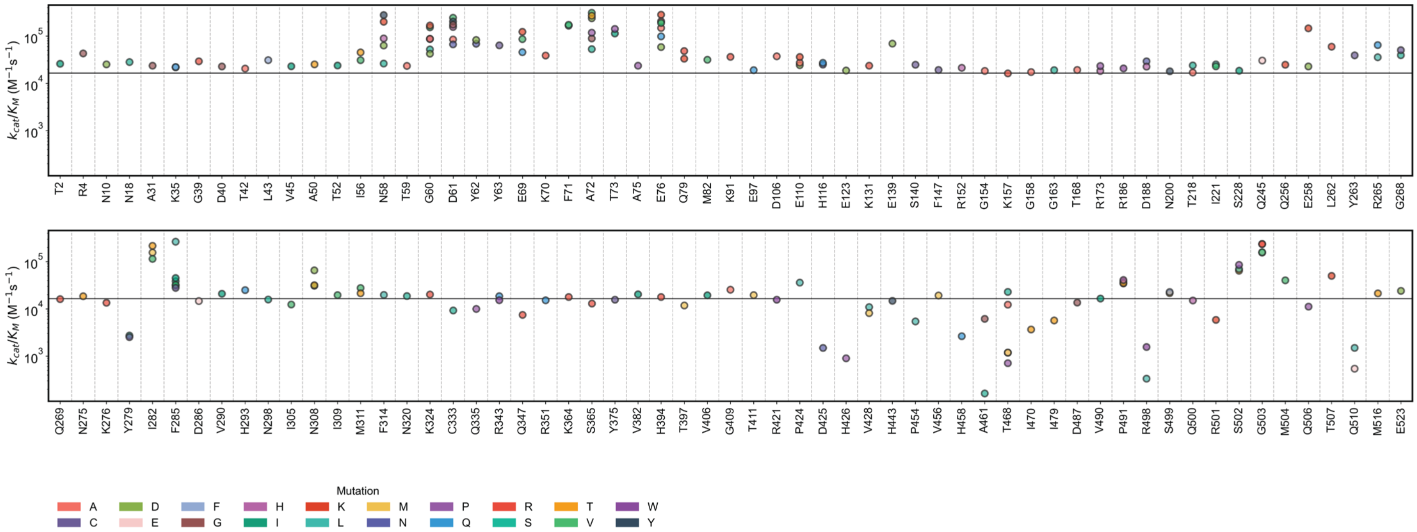


**Figure S8. $k_{cat}/K_M$ of $SHP2_{FL}$ allelic variants by residue position.** Markers represent the median $k_{cat}/K_M$ value for each variant. The horizontal black line indicates the WT median activity. Data are colored by the substituted amino acid at each position.

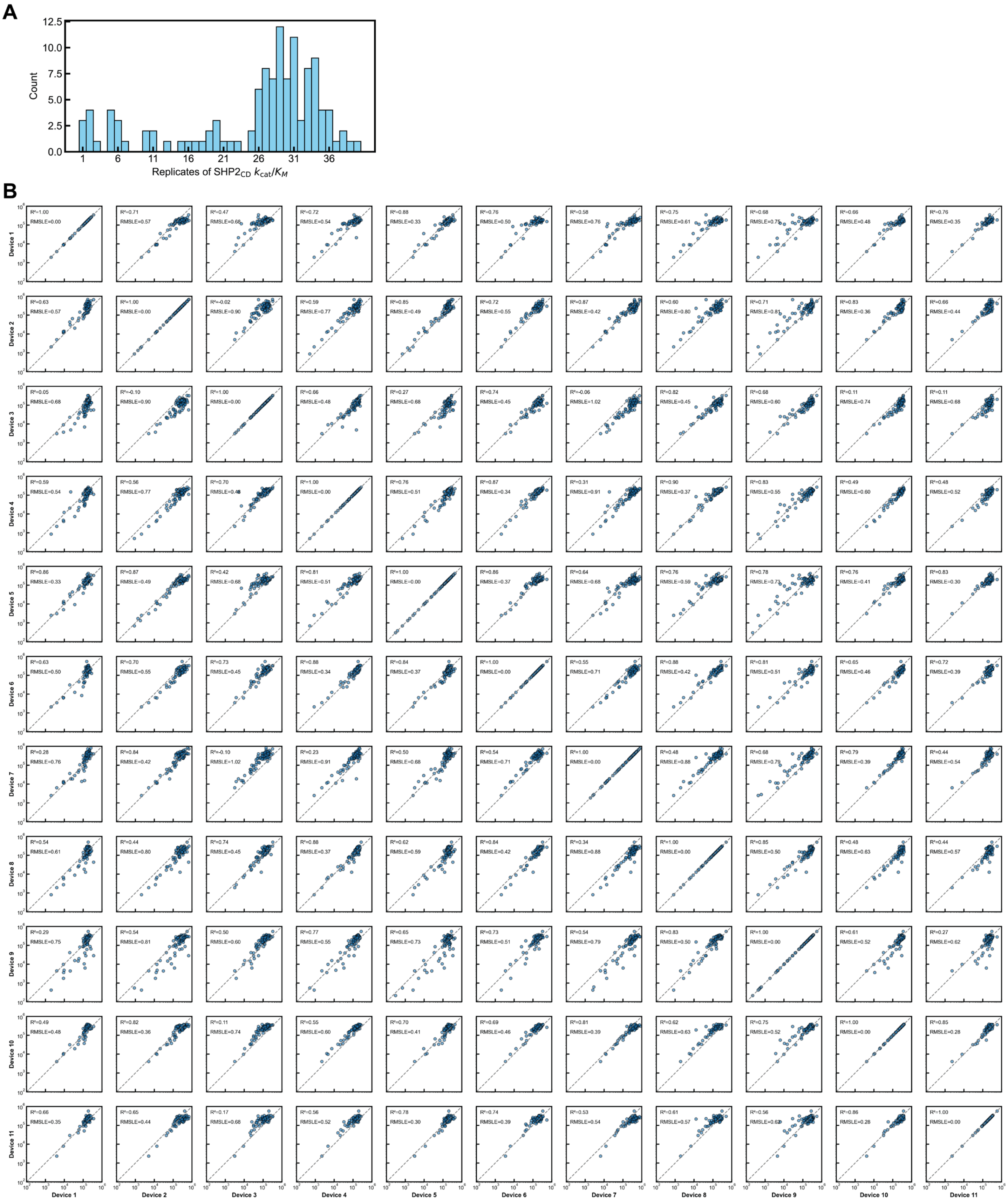


**Figure S9. Reproducibility of SHP2$_{CD}$ DiFMUP hydrolysis activity measurements. (A)** Histogram of number of Michaelis-Menten parameter measurement replicates across 95 SHP2$_{CD}$ variants. **(B)** Reproducibility of $k_{cat}/K_M$ for DiFMUP hydrolysis across device replicates. Each device reports a consensus value derived from 1–4 technical replicates measured in different chambers for the same variant on the same chip. For variants with a single technical replicate, the single measured value is reported; for those with two replicates, the mean is reported; and for variants with three or more replicates, the median is reported. All values are in $M^{-1}s^{-1}$.

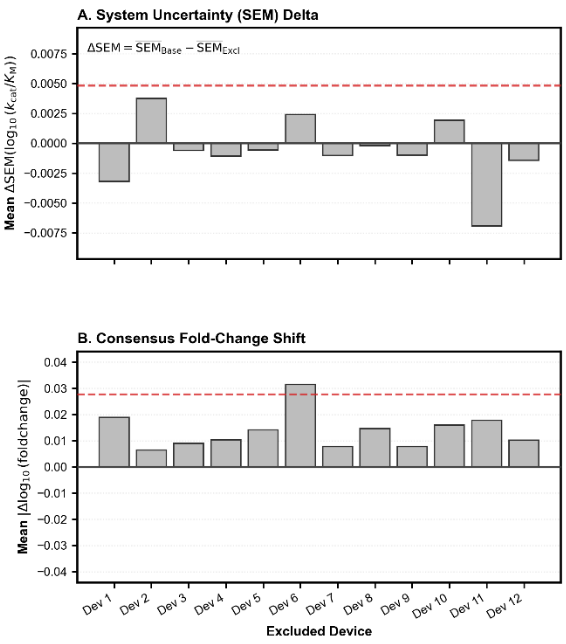


**Figure S10. Leave-one-device-out quality-control analysis of SHP2$_{CD}$ $k_{cat}/K_M$ measurements. (A)** Change in system uncertainty, quantified as the difference in the mean standard error of the mean (SEM) of $\log_{10}(k_{cat}/K_M)$ across variants when each device is excluded in turn relative to the global baseline including all devices. Positive values indicate reduced uncertainty upon exclusion of the device. **(B)** Stability of consensus measurements assessed by the mean absolute shift across all variants in $\log_{10}$(fold change) (difference in $\log_{10}(k_{cat}/K_M)$ relative to an internal anchor variant) compared to the global consensus upon exclusion of each device The anchor was chosen as the most consistently observed variant across devices. Larger values indicate greater deviation from the global consensus upon exclusion of the device. Dashed lines indicate mean + 2 SD thresholds. Because no single device exceeded the 2 SD threshold for both metrics, all devices were retained and none were excluded from the final analysis.

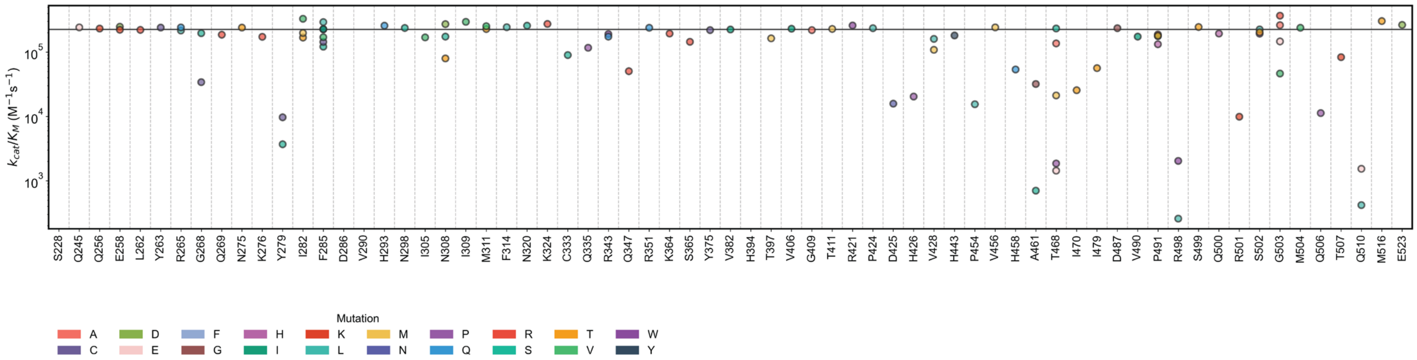


**Figure S11. $k_{cat}/K_M$ of SHP2$_{CD}$ allelic variants by mutation position.** Markers represent the median $k_{cat}/K_M$ value for each variant. The horizontal black line indicates the WT median activity. Data are colored by the substituted amino acid at each position.

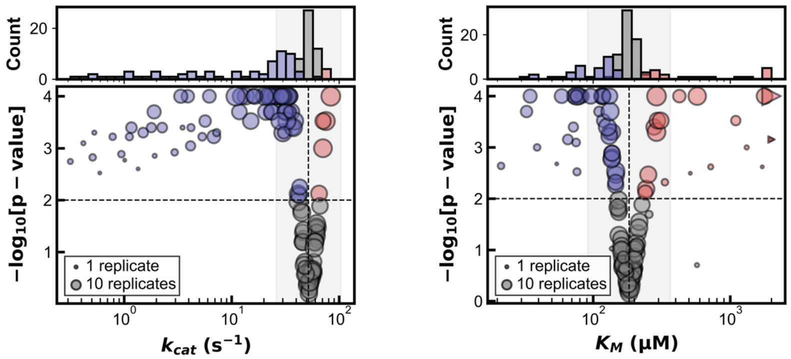


**Figure S12. Median $k_{cat}$ and $K_M$ for DiFMUP hydrolysis across 95 SHP2$_{CD}$ variants.** Triangles indicate mutants with fitted $K_M$ above the highest substrate concentration used; these values are set to the highest substrate concentration used (2000 μM) and represent a lower limit. Blue and red markers indicate variants with a statistically significant decrease or increase compared to WT, respectively ($p < 0.01$). The gray-shaded background denotes a two-fold change range relative to WT.

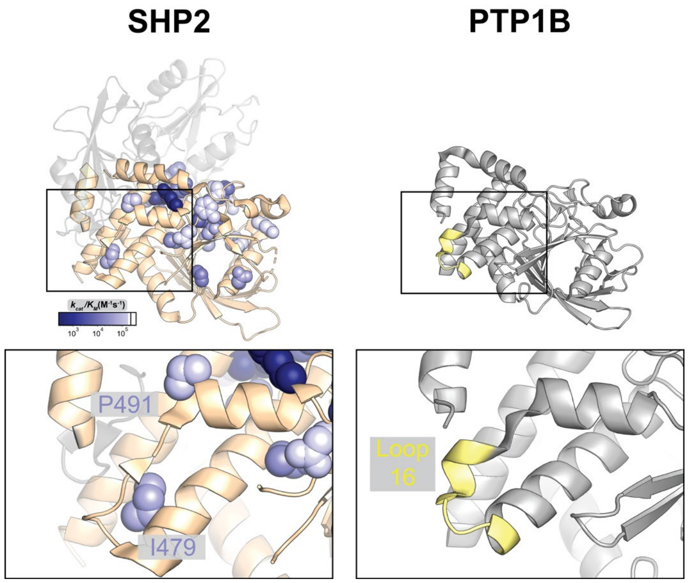


**Figure S13. Locations of residues with potential allosteric impacts on intrinsic catalysis in SHP2 and PTP1B.** View of SHP2 and PTP1B at Loop 16 position. P419 and I479 flank this loop in SHP2.

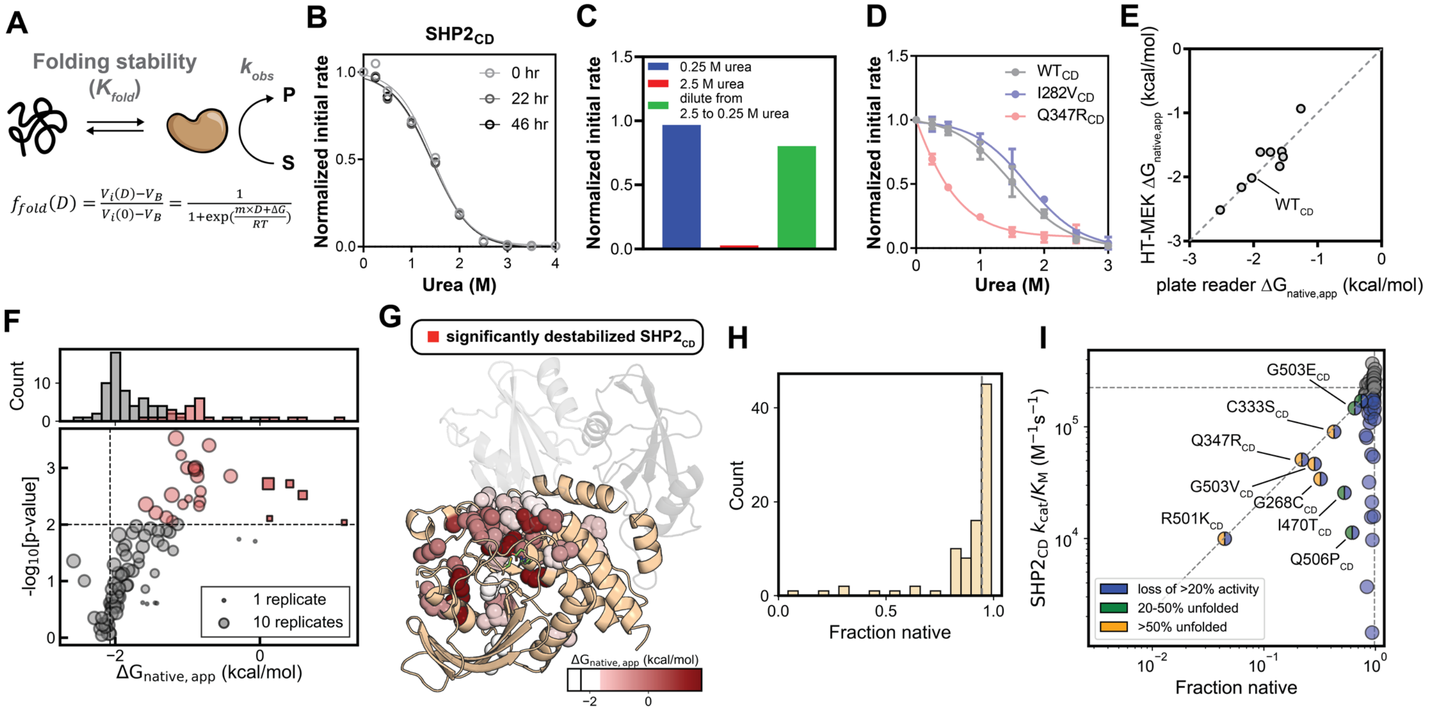


**Figure S14. Measured impacts of SHP2 allelic variants on $SHP2_{CD}$ folding stability. (A)** Schematic illustrating how folding stability can be quantified from urea dose–response curves. **(B)** Representative urea dose-response curves for $SHP2_{CD}$ measured after varying preincubation times with urea. Solid curves represent data fitted to a standard two-state linear extrapolation model. **(C)** Retained activity after treating with 0.25 M urea for 46 hr, 2.5 M urea for 46 hr, and 2.5 M urea for 46 hr followed by a dilution to 0.25 M urea. **(D)** Urea-induced unfolding curves for $SHP2_{CD}$ variants measured by HT-MEK. DiFMUP hydrolysis was measured across urea concentrations with a series of zero urea measurements between each concentration and normalized to the activity in the absence of urea for the immediately preceding measurement (see **Supplementary Text S1**). Solid curves represent data fitted to a standard two-state linear extrapolation model (see **Materials and Methods**). **(E)** Comparison of HT-MEK- and plate reader assay-measured $\Delta G_{native,app}$. **(F)** Median $\Delta G_{native,app}$ of 88 $SHP2_{CD}$ variants. Red: 24 significantly destabilizing variants ($p < 0.01$). Gray: not significantly different from WT ($p \geq 0.01$). Square markers indicate variants that fall below the lower quantitative limit of the assay ($\Delta G_{native,app} < 0$); these variants are >50% non-native at baseline, placing their theoretical transition midpoints outside the physically measurable range of the urea titration. **(G)** Structural mapping of positions with significantly destabilizing mutations in $SHP2_{CD}$ construct. Side chains are shown as red spheres, with intensity reflecting the average stability of all mutations at each position. **(H)** Histogram of natively-folded fraction across $SHP2_{CD}$ variants. **(I)** $SHP2_{CD}$ activity versus natively-folded fraction. Diagonal line indicates 1:1 proportional fold change from WT values, consistent with activity loss originating entirely from loss of native state.

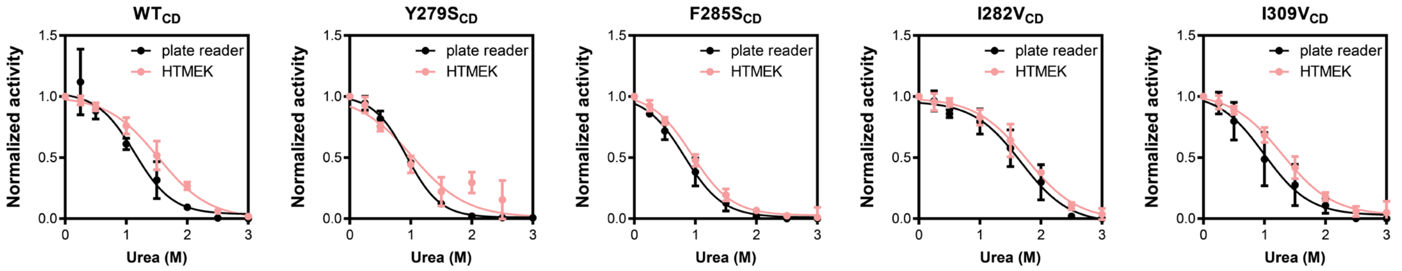


**WT$_{CD}$**

| | $\Delta G_{native, app}$ | *m*-value |
|---|---|---|
| plate reader | -2.2±0.4 | 1.9±0.7 |
| HT-MEK | -2.2±0.1 | 1.4±0.1 |

**Y279S$_{CD}$**

| | $\Delta G_{native, app}$ | *m*-value |
|---|---|---|
| plate reader | -2.0±0.2 | 2.2±0.2 |
| HT-MEK | -1.4±0.8 | 1.3±0.6 |

**F285S$_{CD}$**

| | $\Delta G_{native, app}$ | *m*-value |
|---|---|---|
| plate reader | -1.6±0.3 | 1.9±0.3 |
| HT-MEK | -1.7±0.1 | 1.8±0.1 |

**I282V$_{CD}$**

| | $\Delta G_{native, app}$ | *m*-value |
|---|---|---|
| plate reader | -2.5±0.7 | 1.5±0.4 |
| HT-MEK | -2.5±0.2 | 1.4±0.1 |

**I309V$_{CD}$**

| | $\Delta G_{native, app}$ | *m*-value |
|---|---|---|
| plate reader | -1.6±0.5 | 1.6±0.4 |
| HT-MEK | -1.8±0.2 | 1.4±0.1 |

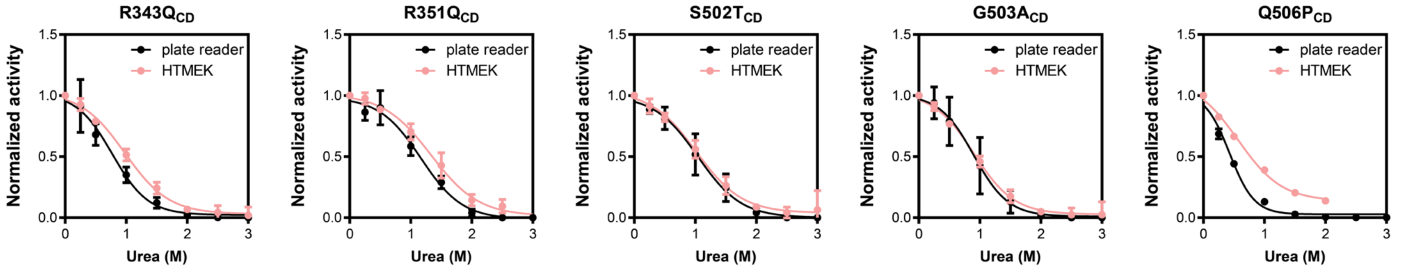


**R343Q$_{CD}$**

| | $\Delta G_{native, app}$ | *m*-value |
|---|---|---|
| plate reader | -1.6±0.4 | 2.0±0.5 |
| HT-MEK | -1.6±0.1 | 1.6±0.1 |

**R351Q$_{CD}$**

| | $\Delta G_{native, app}$ | *m*-value |
|---|---|---|
| plate reader | -2.0±0.5 | 1.7±0.3 |
| HT-MEK | -2.0±0.2 | 1.5±0.1 |

**S502T$_{CD}$**

| | $\Delta G_{native, app}$ | *m*-value |
|---|---|---|
| plate reader | -1.7±0.4 | 1.7±0.3 |
| HT-MEK | -1.6±0.1 | 1.6±0.1 |

**G503A$_{CD}$**

| | $\Delta G_{native, app}$ | *m*-value |
|---|---|---|
| plate reader | -1.9±0.7 | 2.1±0.7 |
| HT-MEK | -1.6±0.1 | 1.8±0.1 |

**Q506P$_{CD}$**

| | $\Delta G_{native, app}$ | *m*-value |
|---|---|---|
| plate reader | -1.3±0.2 | 2.9±0.5 |
| HT-MEK | -0.9±0.3 | 1.6±0.4 |

**Figure S15. Comparison of urea induced unfolding assay performed with plate reader and HT-MEK.** Equilibrium chemical denaturation profiles were measured to test the HT-MEK platform against plate reader-based measurements. Each marker represents the mean of at least 3 replicates, and error bars indicate the standard deviation. Solid lines represent a fit to a two-state linear extrapolation model (see **Materials and Methods** for details). Error represents the 95% CI of the fit. Values for $\Delta G_{native,app}$ are expressed in units of kcal/mol.

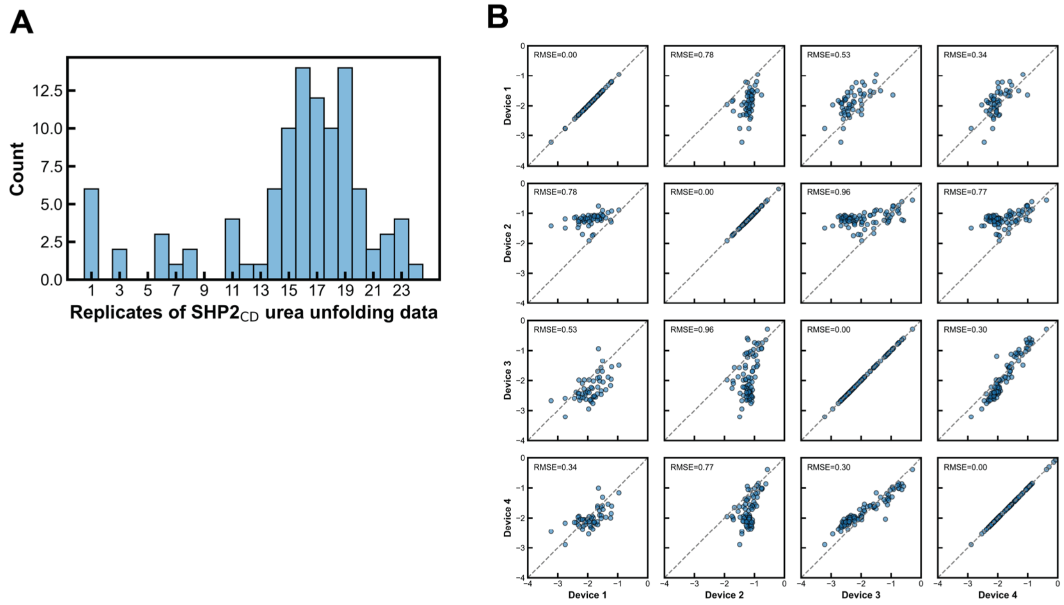


**Figure S16. Reproducibility of $SHP2_{CD}$ folding stability measurements. (A)** Histogram of urea dose-response measurement replicates across $SHP2_{CD}$ variants. **(B)** Reproducibility of $SHP2_{CD}$ $\Delta G_{native,app}$ across device replicates. Each device reports the median of 1–4 replicates from different chambers for the same variant on the same chip. All values are in kcal/mol.

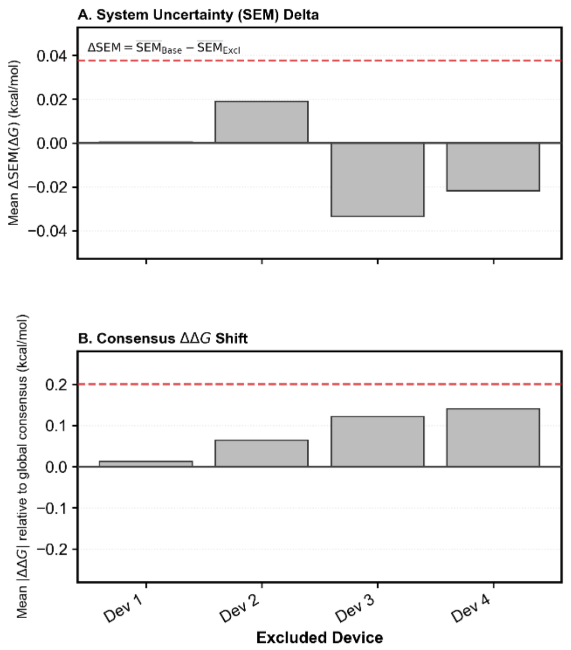


**Figure S17. Leave-one-device-out quality-control analysis of SHP2$_{CD}$ ΔΔG measurements. (A)** Change in system uncertainty, quantified as the difference in the mean standard error of the mean (SEM) of ΔΔG (relative free energy) across variants when each device is excluded in turn relative to the global baseline including all devices. Positive values indicate reduced uncertainty upon exclusion of the device. **(B)** Stability of consensus measurements assessed by the mean absolute shift across all variants in ΔΔG, defined as the difference in ΔG relative to an internal anchor variant, compared to the global consensus upon exclusion of each device. The anchor was chosen as the most consistently observed variant across devices. Larger values indicate greater deviation from the global consensus upon exclusion of the device. Dashed lines indicate mean + 2 SD thresholds. Because no single device exceeded the 2 SD threshold for both metrics, all devices were retained and none were excluded from the final analysis.

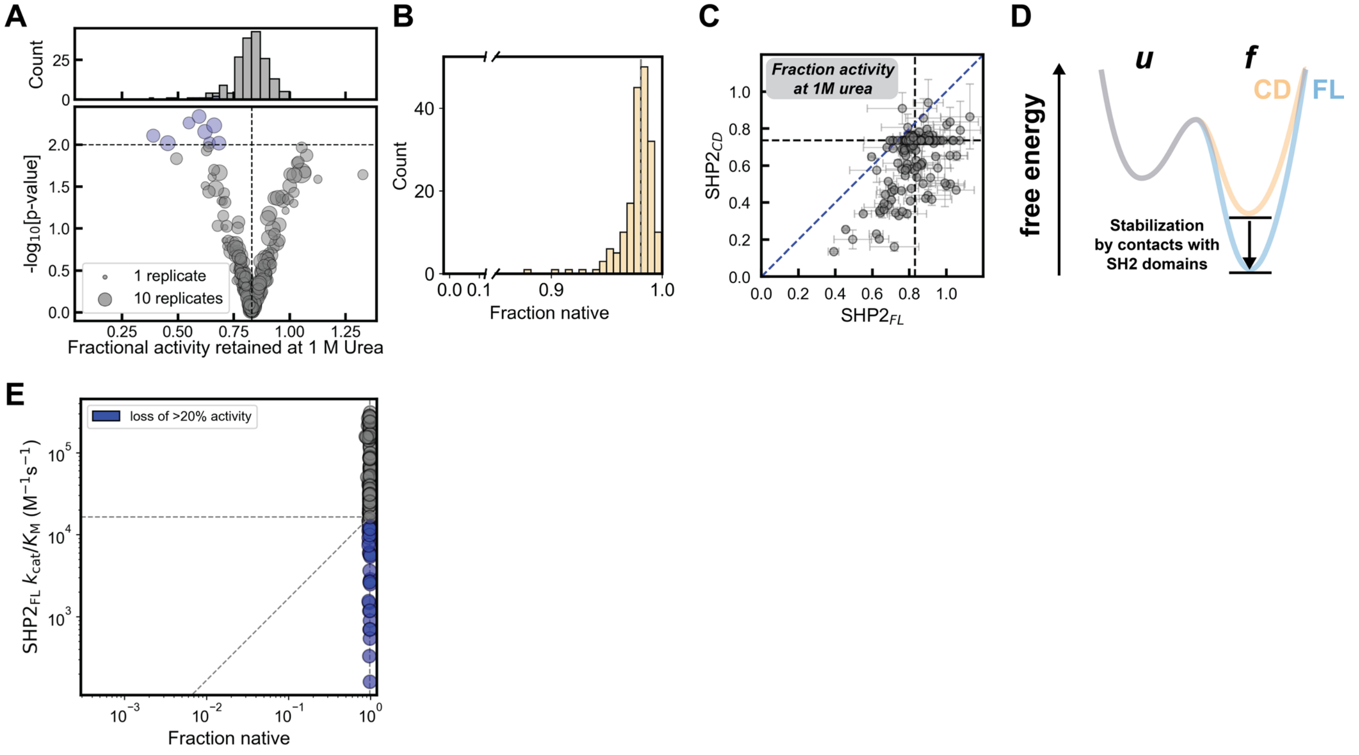


**Figure S18. Estimating the impact of mutations on thermodynamic stability for $SHP2_{FL}$. (A)** Median fractional activity remaining at 1 M urea for $SHP2_{FL}$ variants. **(B)** Histogram of estimated natively-folded fraction at 0 M urea for $SHP2_{FL}$ variants. **(C)** Comparison of fractional activity at 1 M versus 0 M urea for $SHP2_{FL}$ (x-axis) and $SHP2_{CD}$ (y-axis) variants. Dashed lines indicate WT values; diagonal blue line marks 1:1 correspondence. **(D)** Schematic of stabilization effect of the PTP domain from the SH2 domains. **(E)** $SHP2_{FL}$ activity versus natively-folded fraction. Diagonal line indicates proportional fold change from WT values.

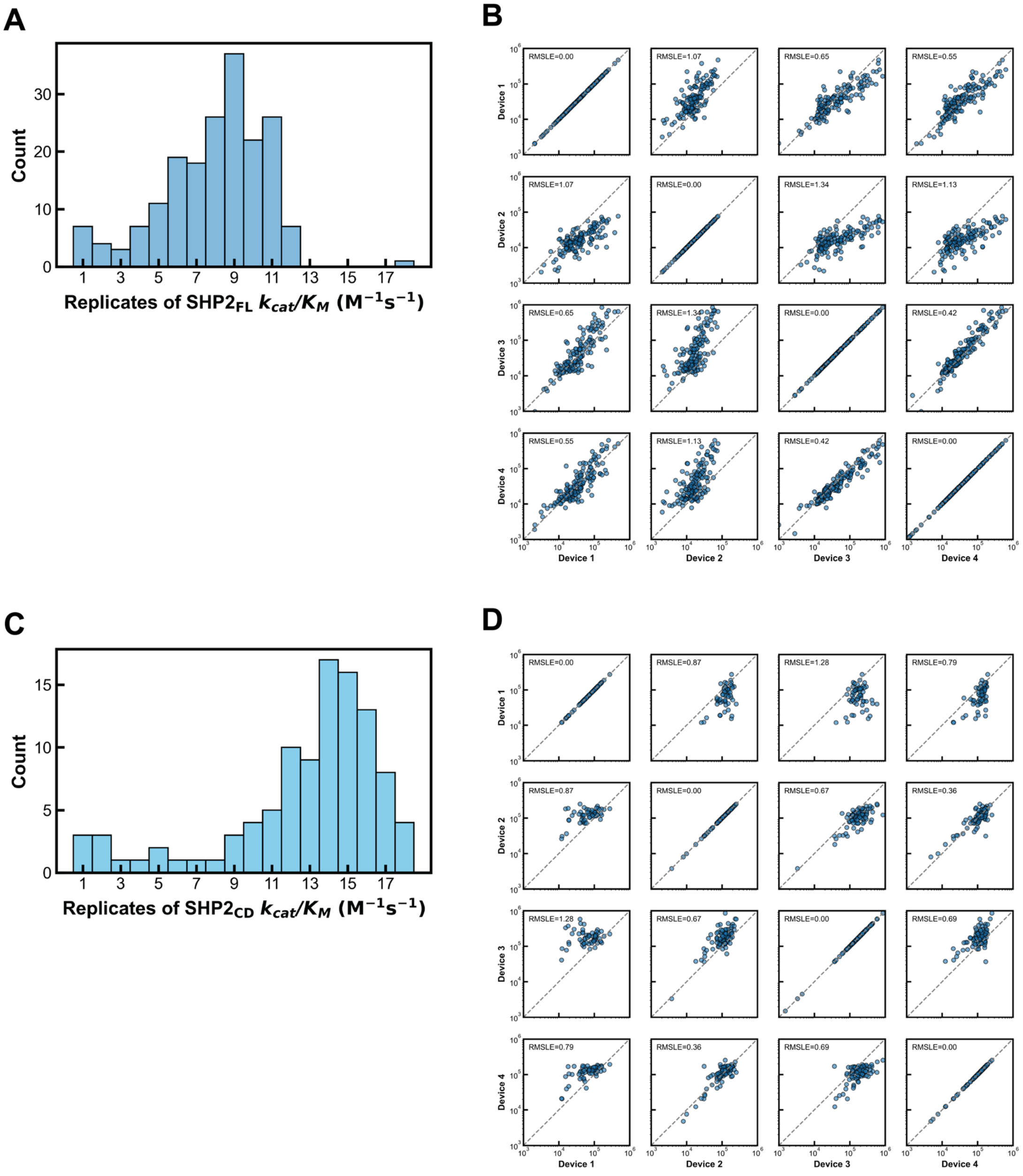


**Figure S19. Reproducibility of SHP2 EGFRpY992 hydrolysis activity measurements.** **(A)** Histogram of number of EGFRpY992 hydrolysis Michaelis-Menten parameter measurement replicates across $SHP2_{FL}$ variants. **(B)** Reproducibility of $SHP2_{FL}$ $k_{cat}/K_M$ for EGFRpY992 hydrolysis across device replicates. **(C)** Histogram of number of EGFRpY992 hydrolysis Michaelis-Menten parameter measurement replicates across $SHP2_{CD}$ variants. **(D)** Reproducibility of $SHP2_{CD}$ $k_{cat}/K_M$ for EGFRpY992 hydrolysis across device replicates. Each device reports the median of 1–4 replicates from different chambers for the same variant on the same chip. All values are in $M^{-1}s^{-1}$.

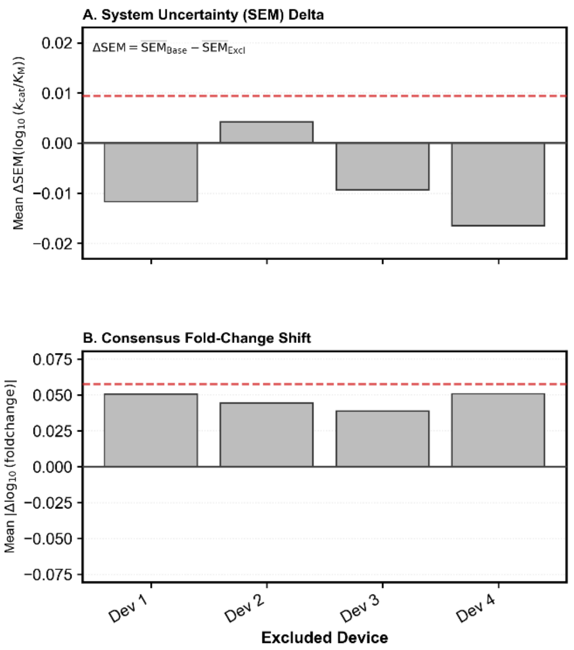


**Figure S20. Leave-one-device-out quality-control analysis of SHP2$_{FL}$ EGFRpY992 hydrolysis $k_{cat}/K_M$ measurements. (A)** Change in system uncertainty, quantified as the difference in the mean standard error of the mean (SEM) of $\log_{10}(k_{cat}/K_M)$ across variants when each device is excluded in turn relative to the global baseline including all devices. Positive values indicate reduced uncertainty upon exclusion of the device. **(B)** Stability of consensus measurements assessed by the mean absolute shift across all variants in $\log_{10}$(fold change) (difference in $\log_{10}(k_{cat}/K_M)$ relative to an internal anchor variant) compared to the global consensus upon exclusion of each device The anchor was chosen as the most consistently observed variant across devices. Larger values indicate greater deviation from the global consensus upon exclusion of the device. Dashed lines indicate mean + 2 SD thresholds. Because no single device exceeded the 2 SD threshold for both metrics, all devices were retained and none were excluded from the final analysis.

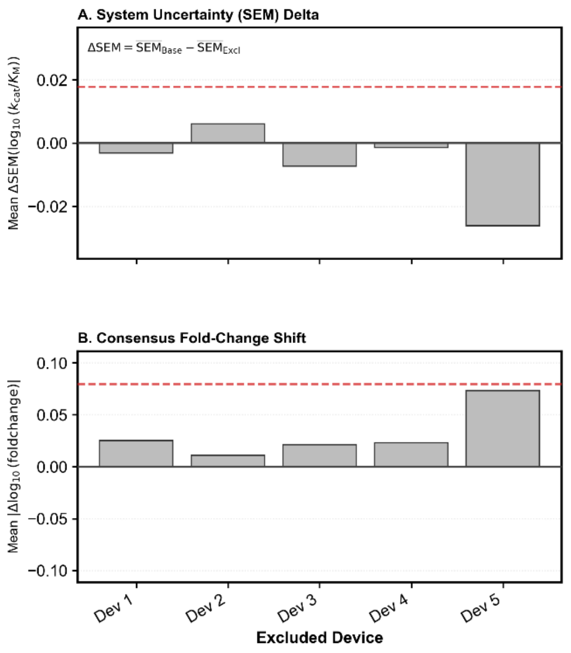


**Figure S21. Leave-one-device-out quality-control analysis of SHP2$_{CD}$ EGFRpY992 hydrolysis $k_{cat}/K_M$ measurements. (A)** Change in system uncertainty, quantified as the difference in the mean standard error of the mean (SEM) of $\log_{10}(k_{cat}/K_M)$ across variants when each device is excluded in turn relative to the global baseline including all devices. Positive values indicate reduced uncertainty upon exclusion of the device. **(B)** Stability of consensus measurements assessed by the mean absolute shift across all variants in $\log_{10}$(fold change) (difference in $\log_{10}(k_{cat}/K_M)$ relative to an internal anchor variant) compared to the global consensus upon exclusion of each device The anchor was chosen as the most consistently observed variant across devices. Larger values indicate greater deviation from the global consensus upon exclusion of the device. Dashed lines indicate mean + 2 SD thresholds. Because no single device exceeded the 2 SD threshold for both metrics, all devices were retained and none were excluded from the final analysis.

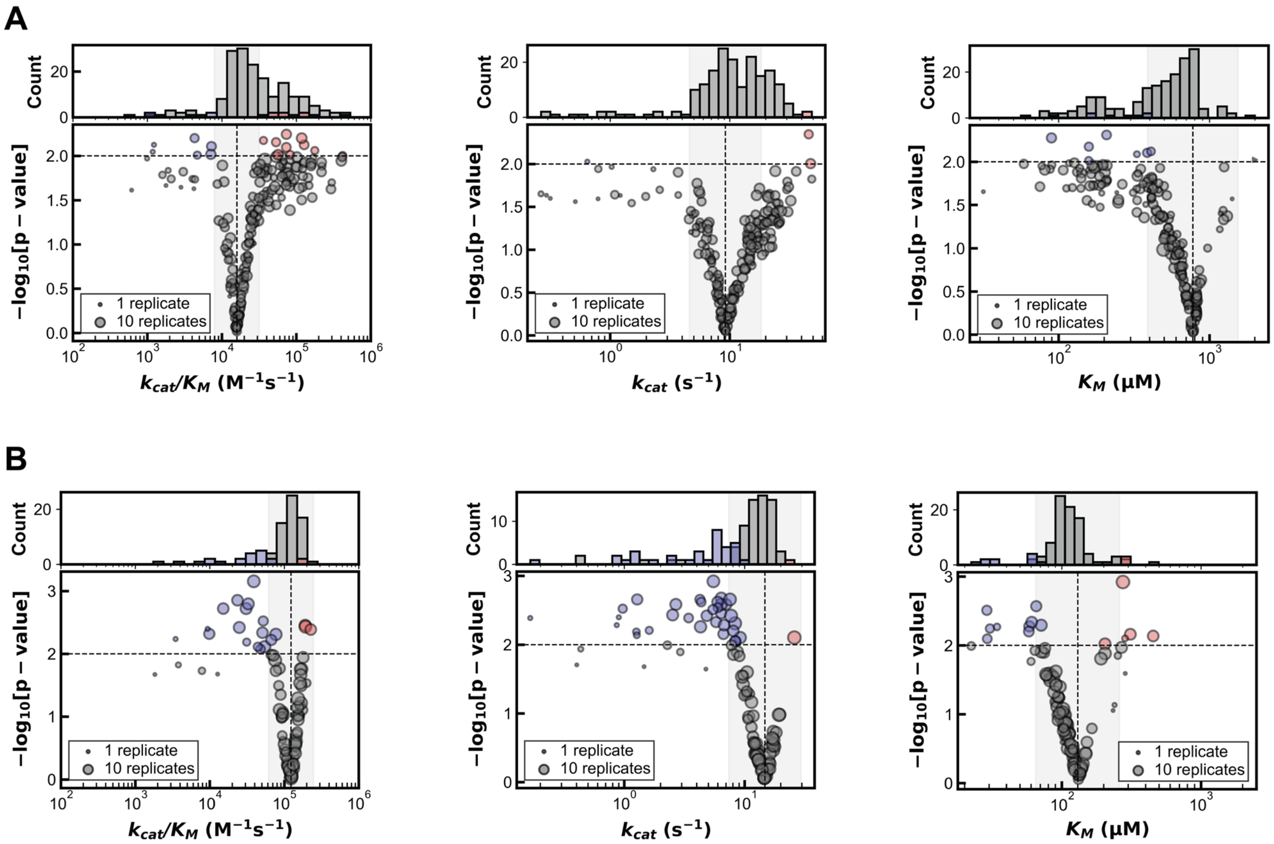


**Figure S22. Volcano plots for measured EGFRpY992 hydrolysis parameters.** Median $k_{cat}/K_M$, $k_{cat}$ and $K_M$ for EGFRpY992 hydrolysis across **(A)** $SHP2_{FL}$ variants and **(B)** $SHP2_{CD}$ variants. Blue and red markers indicate variants with a statistically significant decrease or increase compared to WT, respectively ($p < 0.01$). The gray-shaded background denotes a two-fold change range relative to WT.

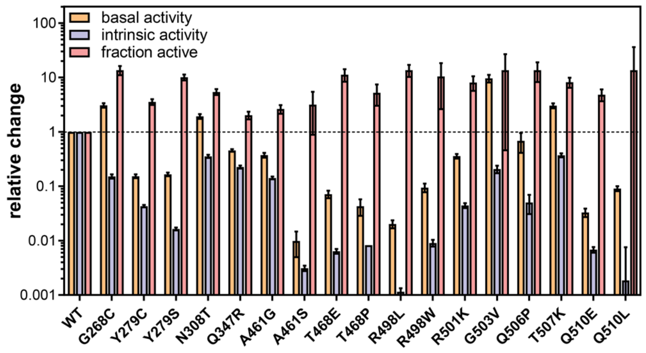


**Figure S23. Variants with both significant reduction in intrinsic activity and increase in fraction active.** Data are presented as mean ± SEM.

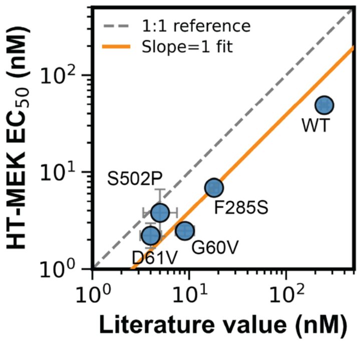


**Figure S24. Comparing HT-MEK measured ppIRS1 $EC_{50}$ to literature values** (*10*). Error bars reflect SEM propagated into $\log_{10}$ space. The dashed line indicates perfect agreement (1:1 reference; RMSE = 0.46 in $\log_{10}$ space). The orange line shows a slope-constrained fit (slope = 1), revealing a systematic 2.6-fold offset where HT-MEK $EC_{50}$ values are consistently lower than literature values ($r^2$ = 0.82), likely reflecting a condition-dependent shift in apparent $EC_{50}$.

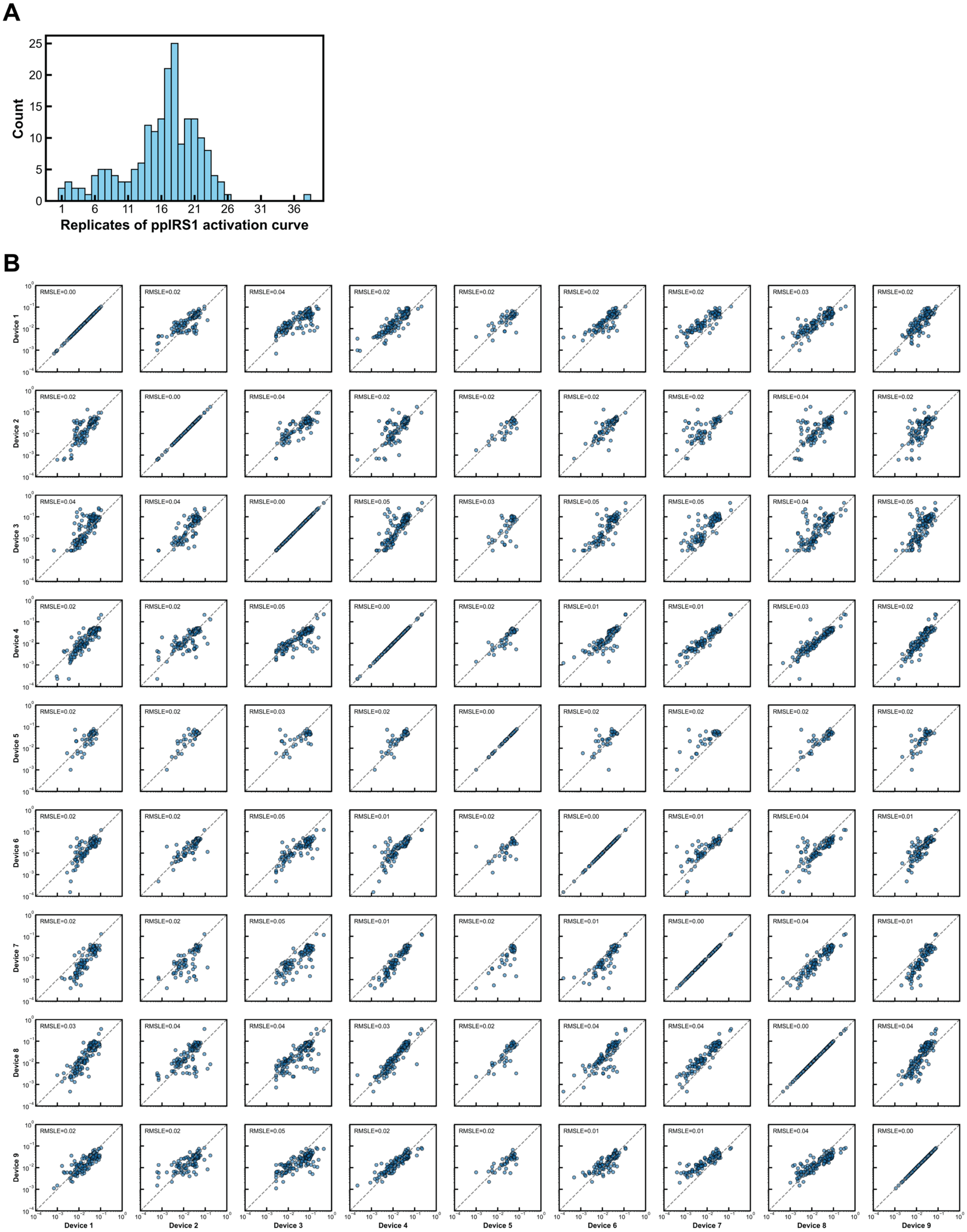


**Figure S25. Reproducibility of ppIRS1 $EC_{50}$ measurements. (A)** Histogram of ppIRS1 dose-response measurement replicates across SHP2 variants. **(B)** Reproducibility of ppIRS1 $EC_{50}$ across device replicates. Each device reports the median of 1–4 replicates from different chambers for the same variant on the same chip. All values are in µM.

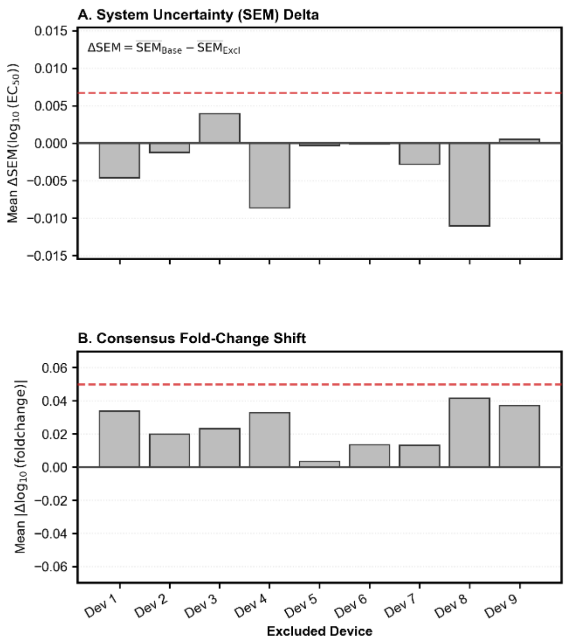


**Figure S26. Leave-one-device-out quality-control analysis of SHP2 $EC_{50}$ measurements derived from ppIRS1 dose-response curve fits. (A)** Change in system uncertainty, quantified as the difference in the mean standard error of the mean (SEM) of $\log_{10}(EC_{50})$ across variants when each device is excluded in turn relative to the global baseline including all devices. Positive values indicate reduced uncertainty upon exclusion of the device. **(B)** Stability of consensus measurements assessed by the mean absolute shift across all variants in $\log_{10}$ fold change (difference in $\log_{10}(EC_{50})$ relative to an internal anchor variant) compared to the global consensus upon exclusion of each device. The anchor was chosen as the most consistently observed variant across devices. Larger values indicate greater deviation from the global consensus upon exclusion of the device. Dashed lines indicate mean + 2 SD thresholds. Because no single device exceeded the 2 SD threshold for both metrics, all devices were retained and none were excluded from the final analysis.

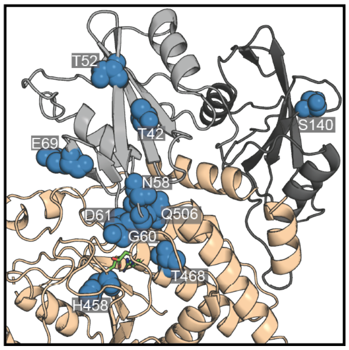


**Figure S27. All positions with mutations with significant deviation from the global $EC_{50}$ versus fraction active fit (Fig. 3K).**

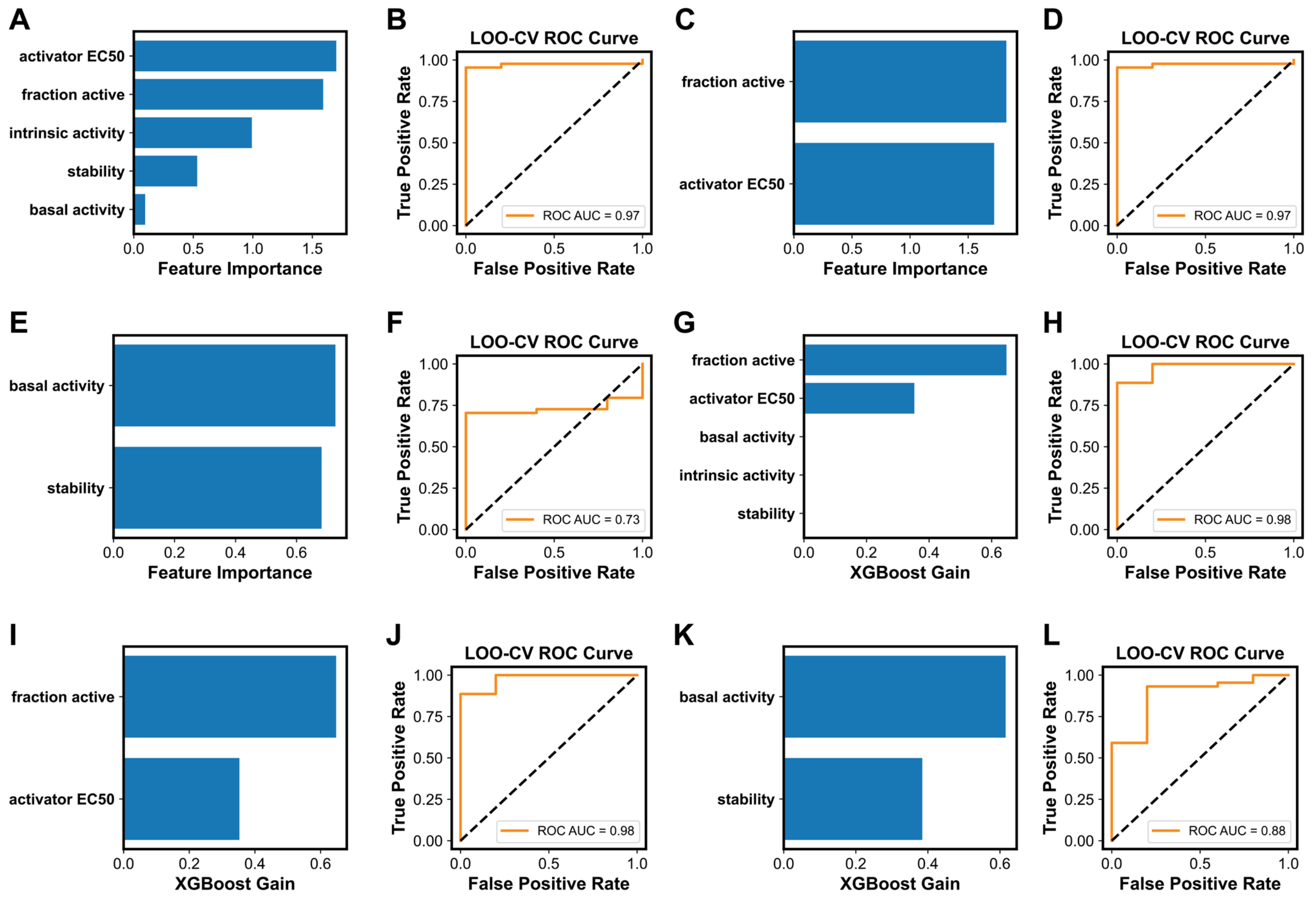


**Figure S28. Identifying biochemical features most predictive of pathogenicity. (A)** Logistic regression feature importance for predicting SHP2 variant pathogenicity. Importance values are absolute model coefficients after *z*-score normalization of biochemical features. **(B)** Receiver operating characteristic (ROC) curve for leave-one-out cross-validation (LOO-CV) classification of SHP2 variants as benign or pathogenic. The area under the curve (AUC) quantifies overall classification performance. **(C)** Logistic regression feature importance and **(D)** LOO-CV ROC curve for predicting SHP2 variant pathogenicity using only fraction active and ppIRS1 $EC_{50}$. **(E)** Logistic regression feature importance and **(F)** LOO-CV ROC curve for predicting SHP2 variant pathogenicity using only basal activity and stability. **(G)** XGBoost relative importance (gain) of each feature in distinguishing pathogenic from benign SHP2 variants. **(H)** LOO-CV ROC curve showing the performance of the XGBoost classifier. **(I)** XGBoost model feature importance and **(J)** LOO-CV ROC curve for predicting SHP2 variant pathogenicity using only fraction active and ppIRS1 $EC_{50}$. **(K)** XGBoost model feature importance and **(L)** LOO-CV ROC curve for predicting SHP2 variant pathogenicity using only basal activity and stability.

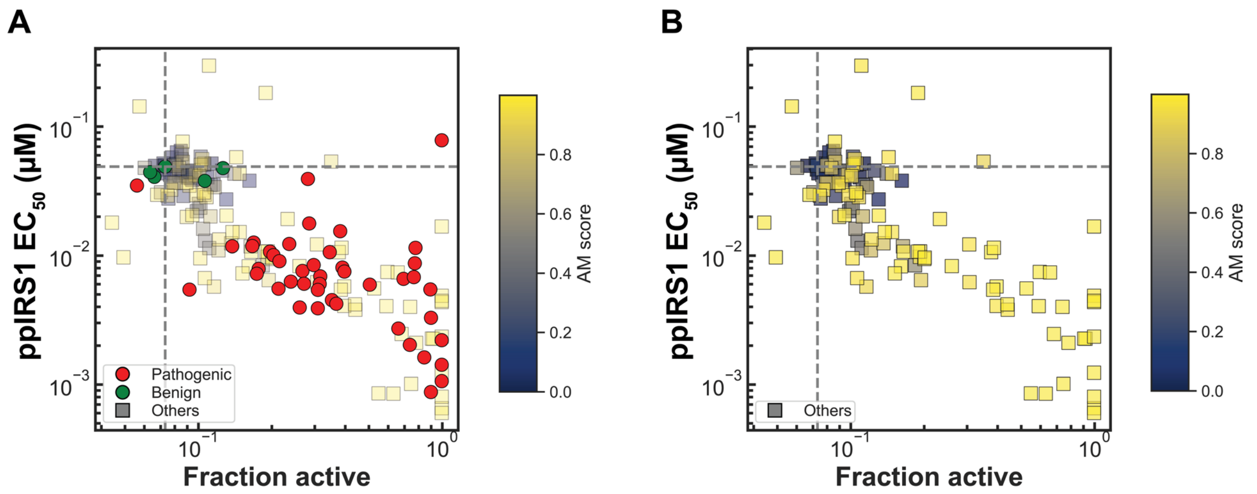


**Figure S29. SHP2 variant ppIRS1 $EC_{50}$ versus fraction active**. **(A)** ClinVar pathogenic and benign variants are shown in red and green, respectively; other variants are plotted as squares and colored by AlphaMissense (AM) pathogenicity score. Dashed lines indicate WT values. **(B)** Variants other than ClinVar pathogenic and benign variants plotted alone to highlight the distribution of AM scores.

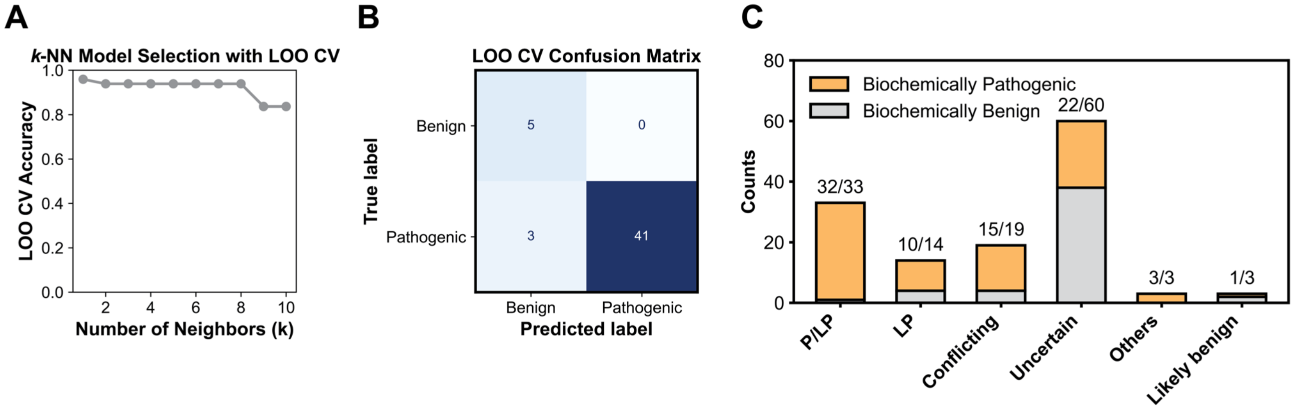


**Figure S30. Predicting likely pathogenicity for variants with uncertain or low-confidence clinical annotations.** Classification of SHP2 variants with uncertain or low-confidence clinical annotations using a *k*-nearest neighbor model trained on fraction active and activator $EC_{50}$. **(A)** Number of neighbors (*k*) selection test for *k* = 1–10. **(B)** LOO-CV confusion matrix for *k* = 1. **(C)** Class prediction result of *k*-nearest neighbor model with *k* = 1. Bars show the number of variants classified as biochemically pathogenic (orange) or biochemically benign (gray) within each clinical significance category. Numbers above bars indicate the number classified as biochemically pathogenic/total variants in each category.

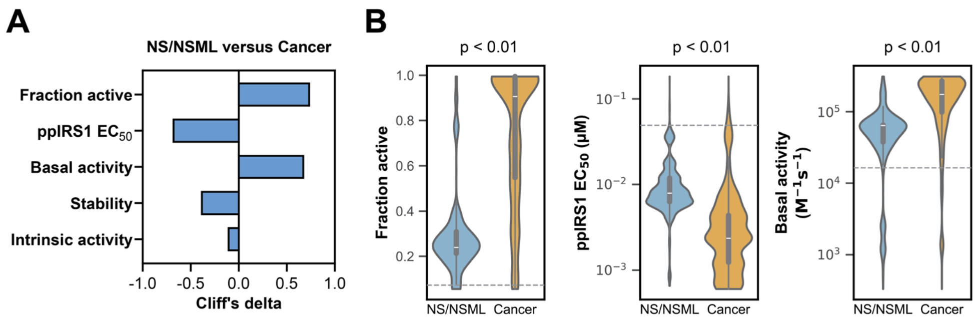


**Figure S31. Analysis of biochemical feature differences between NS/NSML and cancer. (A)** Cliff's delta (a non-parametric measure quantifying the probability that a value from one group exceeds the other, minus the probability of the reverse) for each biochemical feature comparing NS/NSML versus cancer cases, ranked by absolute effect size. Positive values indicate higher feature values in cancer variants, while negative values indicate higher values in NS/NSML variants. Larger absolute values indicate greater differences between groups. **(B)** Distribution of fraction active, activator sensitivity, and basal activity for variants reported in NS/NSML versus cancer cases. *p*-values reflect the probability of observing the difference in medians by chance, calculated against a null distribution generated by pooled bootstrap resampling. Dashed gray lines mark the WT values.

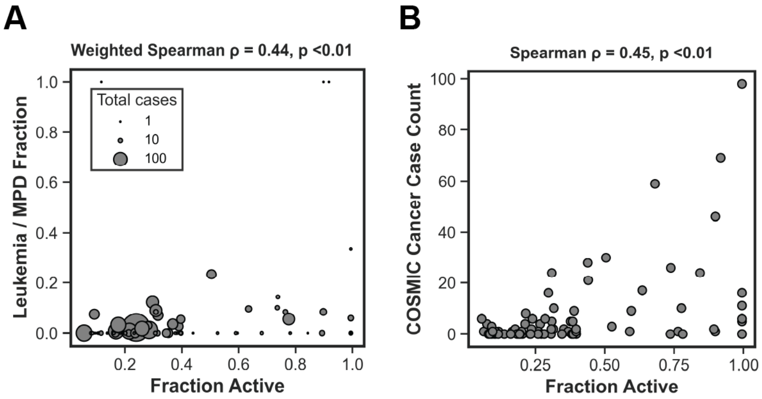


**Figure S32. Fraction active of NS/NSML variants correlates with blood cancer prevalence. (A)** Each point represents an NS/NSML SHP2 variant documented in NSEuroNet, where the y axis represents the fraction of NS/NSML patients who harbor the variant and developed leukemia or myeloproliferative disorder (MPD), and the x axis is the fraction active of the variant. Variants with higher fraction active show a modest but significant increase in leukemia/MPD occurrence. Spearman $\rho$ computed using a weighted Spearman correlation; $p$-value obtained by permutation test. **(B)** Number of COSMIC-reported blood-cancer cases versus fraction active for each NS/NSML variant documented in NSEuroNet. The COSMIC counts reflect all cancer patients in the database and therefore report how often these germline disease variants appear as somatic mutations in cancer. Variants with higher fraction active show a modest but significantly higher occurrence as somatic mutations in cancer; $p$-value obtained by permutation test.

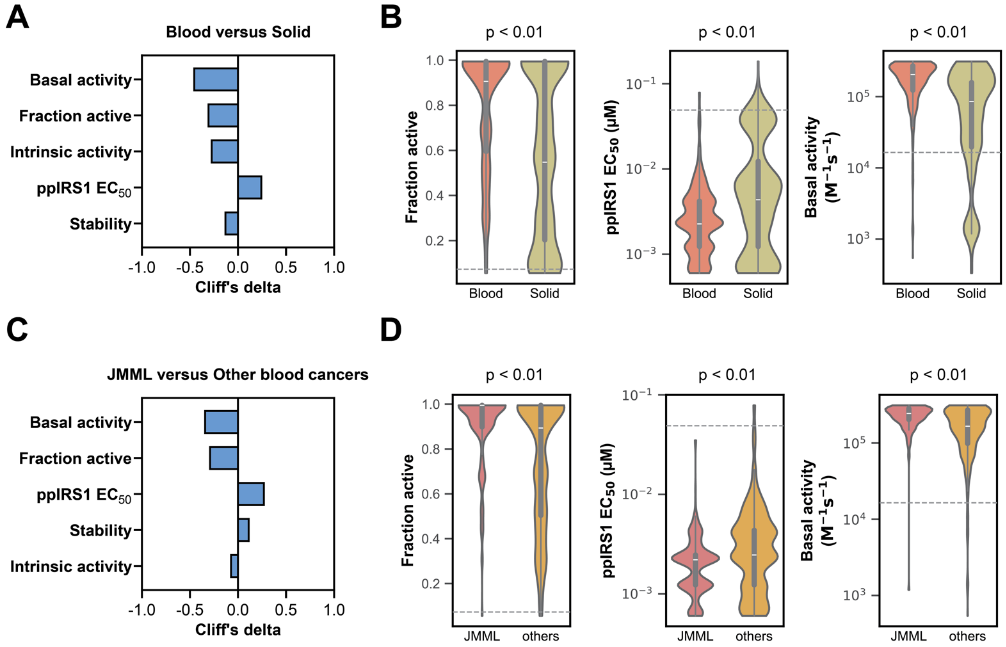


**Figure S33. Analysis of biochemical feature differences between cancer subtypes. (A)** Cliff's delta (a non-parametric measure quantifying the probability that a value from one group exceeds the other, minus the probability of the reverse) for each biochemical feature comparing blood cancer versus solid cancer cases, ranked by absolute effect size. **(B)** Distribution of fraction active, activator sensitivity, and basal activity for variants reported in blood versus solid cancer cases**. (C)** Cliff's delta for each biochemical feature comparing JMML versus other blood cancer cases, ranked by absolute effect size. **(D)** Distribution of fraction active, activator sensitivity, and basal activity for variants reported in JMML versus other blood cancer cases**.** *p*-values reflect the probability of observing the difference in medians by chance, calculated against a null distribution generated by pooled bootstrap resampling. Dashed gray lines mark the WT values.

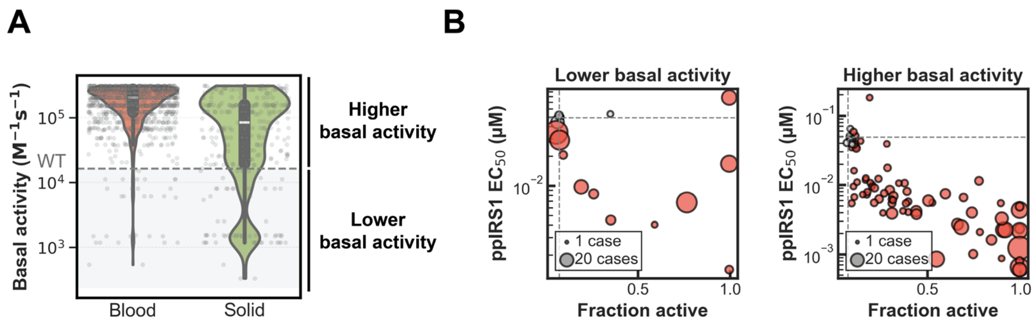


**Figure S34. Cancer variants are primarily associated with an increased fraction active or activator sensitivity, regardless of basal activity. (A)** Distribution of basal activity for variants reported in blood and solid cancer cases. Grey dashed line marks WT value. **(B)** Comparison of ppIRS1 $EC_{50}$ versus fraction active for cancer variants with lower or higher basal activity than WT. Marker size is proportional to the number of case reports per variant. Variants with significant increase in fraction active or reduction in $EC_{50}$ ($p < 0.01$) compared to WT are highlighted in red.

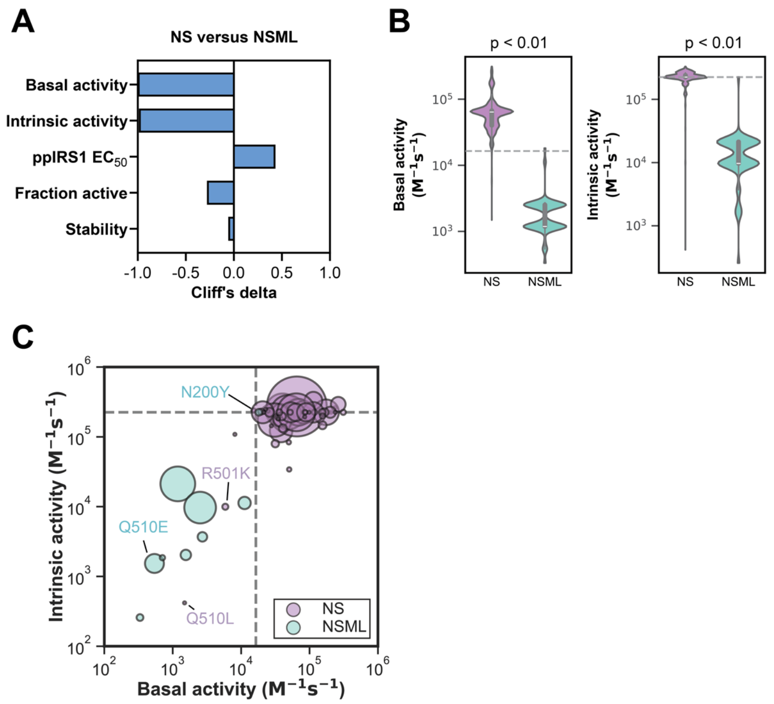


**Figure S35. Analysis of biochemical feature differences between NS and NSML. (A)** Cliff's delta (a non-parametric measure quantifying the probability that a value from one group exceeds the other, minus the probability of the reverse) for each biochemical feature comparing NS versus NSML cases, ranked by absolute effect size. **(B)** Distribution of basal activity and intrinsic activity for variants reported in NS versus NSML cases**.** **(C)** Scatter plot of intrinsic activity versus basal activity for variants reported as NS or NSML. Marker size is proportional to the number of case reports for each variant. Dashed gray lines mark the WT values.

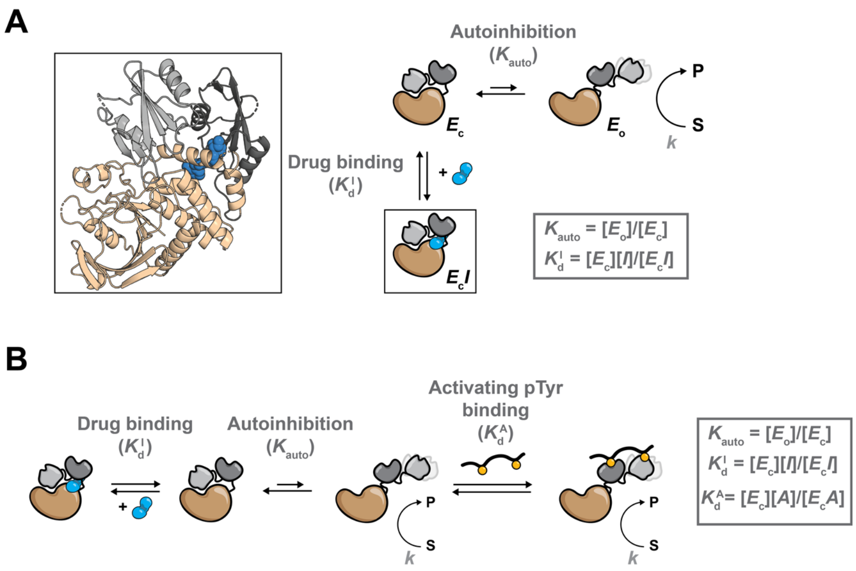


**Figure S36. Two-conformational-state inhibition model for SHP2 allosteric inhibition. (A)** Two-conformational-state inhibition model for SHP2 allosteric inhibition. **(B)** Antagonistic model of simultaneous treatment of SHP2 with allosteric inhibitors and activators.

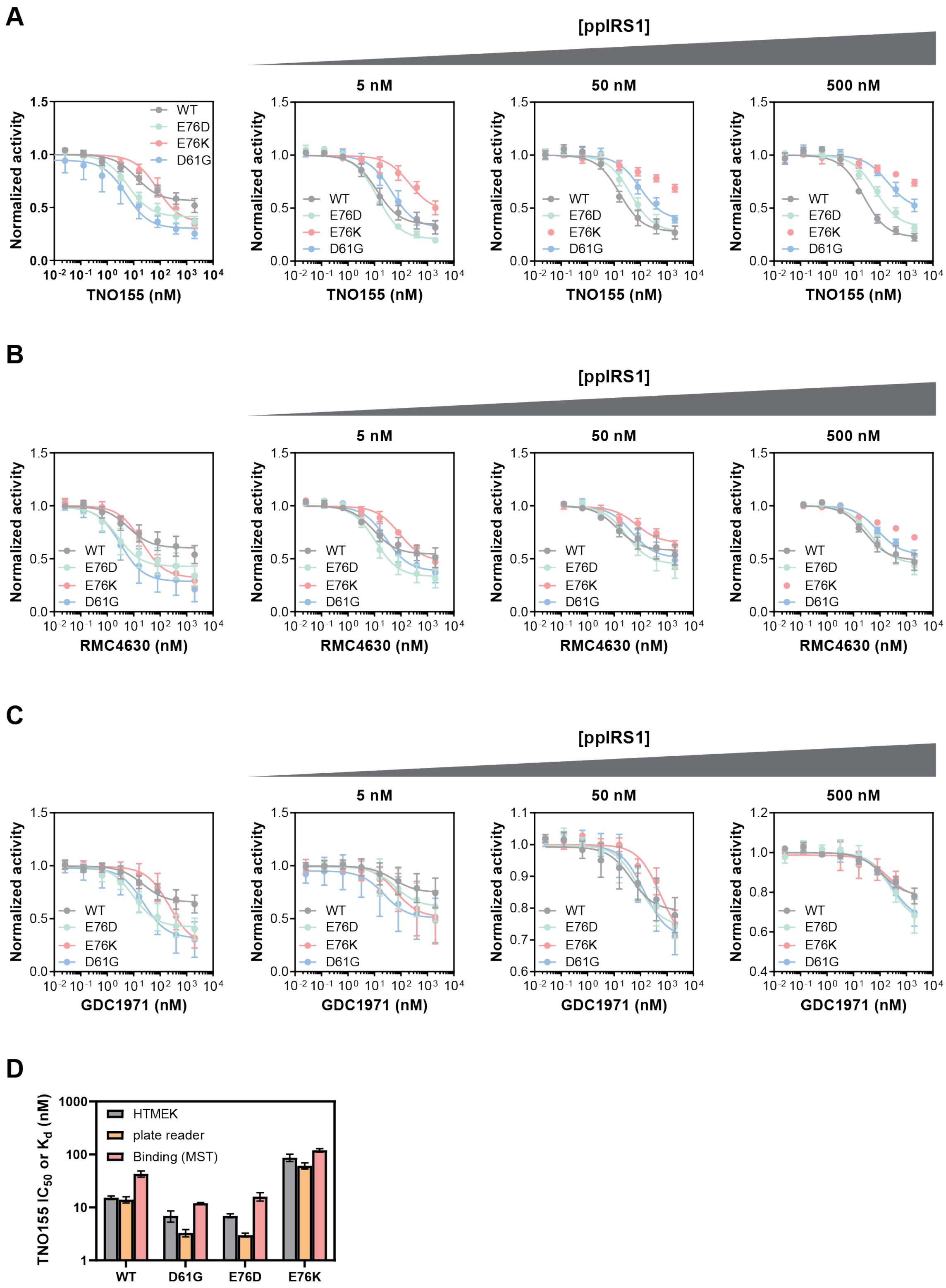


**Figure S37. Dose-response curves for representative SHP2 variants as a function of ppIRS1 concentration. (A, B, C)** HT-MEK-measured dose-response curves for 4 representative SHP2 variants across 4 concentrations of ppIRS1 for TNO155 **(A)**, RMC-4630 **(B)** and GDC-1971 **(C)**. **(D)** HT-MEK-measured $IC_{50}$ compared to plate reader measured $IC_{50}$ and microscale thermophoresis (MST) measured $K_d$.

**A**

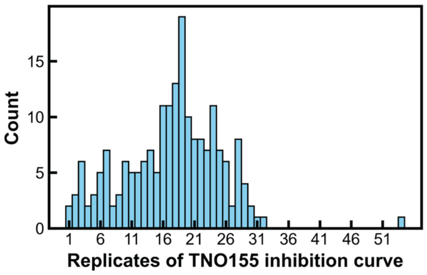


**B**

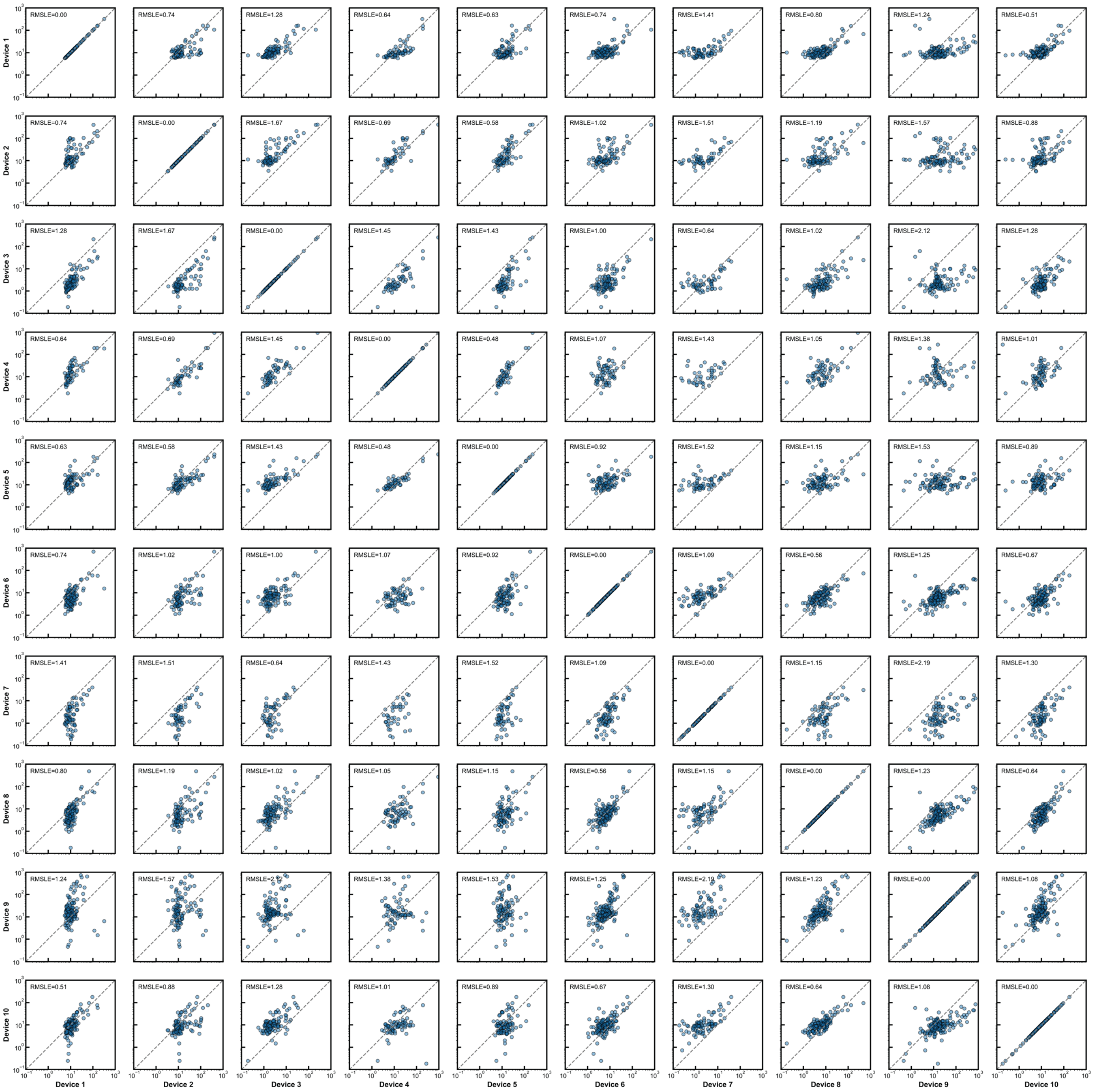


**Figure S38. Reproducibility of TNO155 $IC_{50}$ measurements. (A)** Histogram of TNO155 $IC_{50}$ measurement replicates across 190 $SHP2_{FL}$ variants. **(B)** Reproducibility of TNO155 $IC_{50}$ across device replicates. Each device reports the median of 1–4 replicates from different chambers for the same variant on the same chip. All values are in nM.

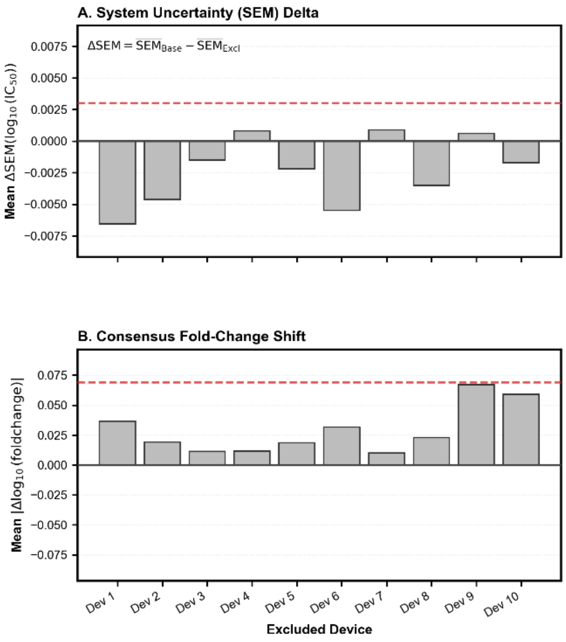


**Figure S39. Leave-one-device-out quality-control analysis of SHP2 TNO155 $IC_{50}$ measurements. (A)** Change in system uncertainty, quantified as the difference in the mean standard error of the mean (SEM) of $\log_{10}(IC_{50})$ across variants when each device is excluded in turn relative to the global baseline including all devices. Positive values indicate reduced uncertainty upon exclusion of the device. **(B)** Stability of consensus measurements assessed by the mean absolute shift across all variants in $\log_{10}$ fold change (difference in $\log_{10}(IC_{50})$ relative to an internal anchor variant) compared to the global consensus upon exclusion of each device. The anchor was chosen as the most consistently observed variant across devices. Larger values indicate greater deviation from the global consensus upon exclusion of the device. Dashed lines indicate mean + 2 SD thresholds. Because no single device exceeded the 2 SD threshold for both metrics, all devices were retained and none were excluded from the final analysis.

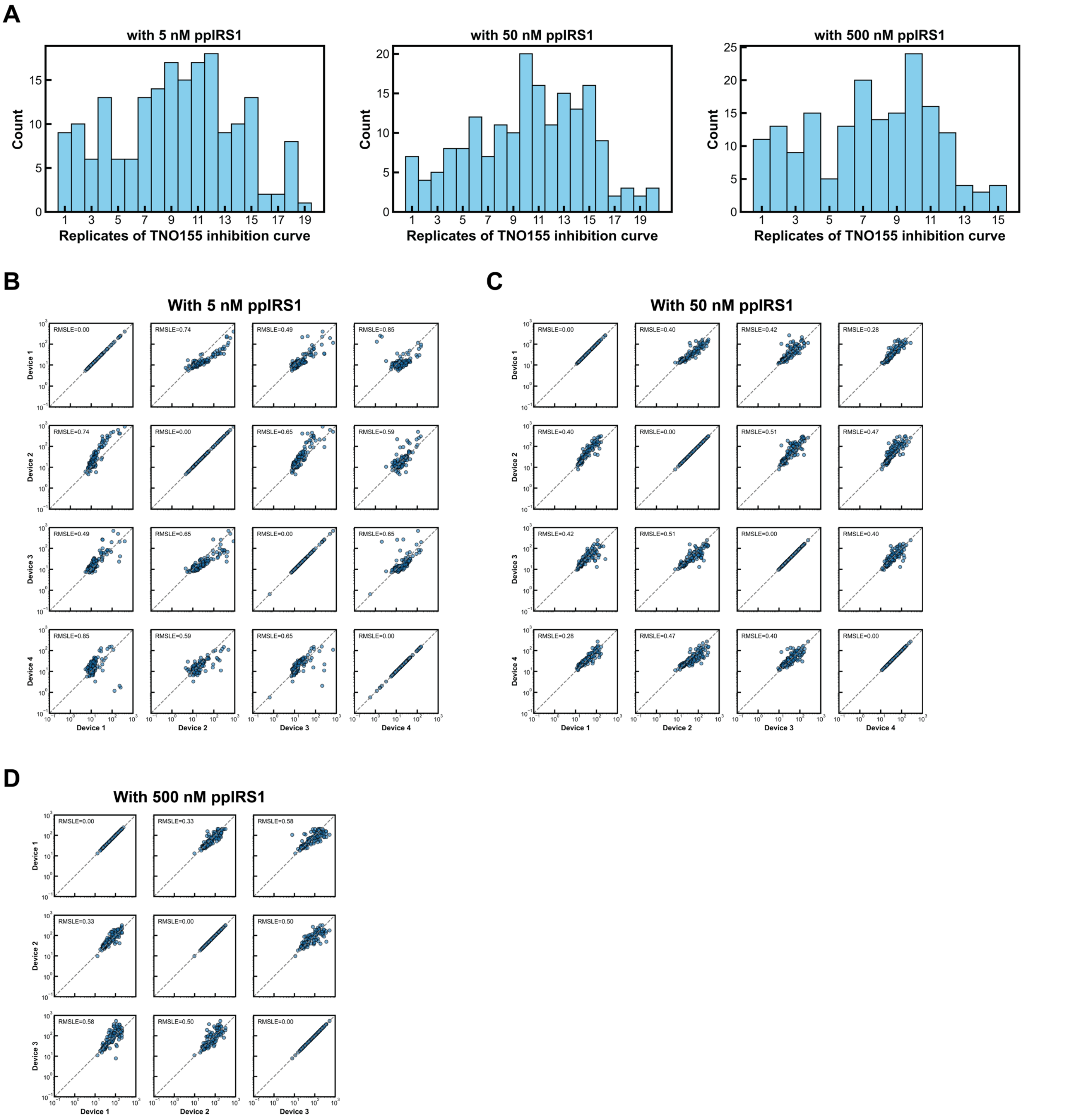


**Figure S40. Reproducibility of TNO155 IC$_{50}$ measurements under different concentrations of ppIRS1.** **(A)** Histogram of replicates for TNO155 IC$_{50}$ measurements under different concentrations of ppIRS1. Reproducibility of TNO155 IC$_{50}$ across device replicates under conditions with **(B)** 5 nM ppIRS1, **(C)** 50 nM ppIRS1, and **(D)** 500 nM ppIRS1. Each device reports the median of 1–4 replicates from different chambers for the same variant on the same chip. All values are in nM.

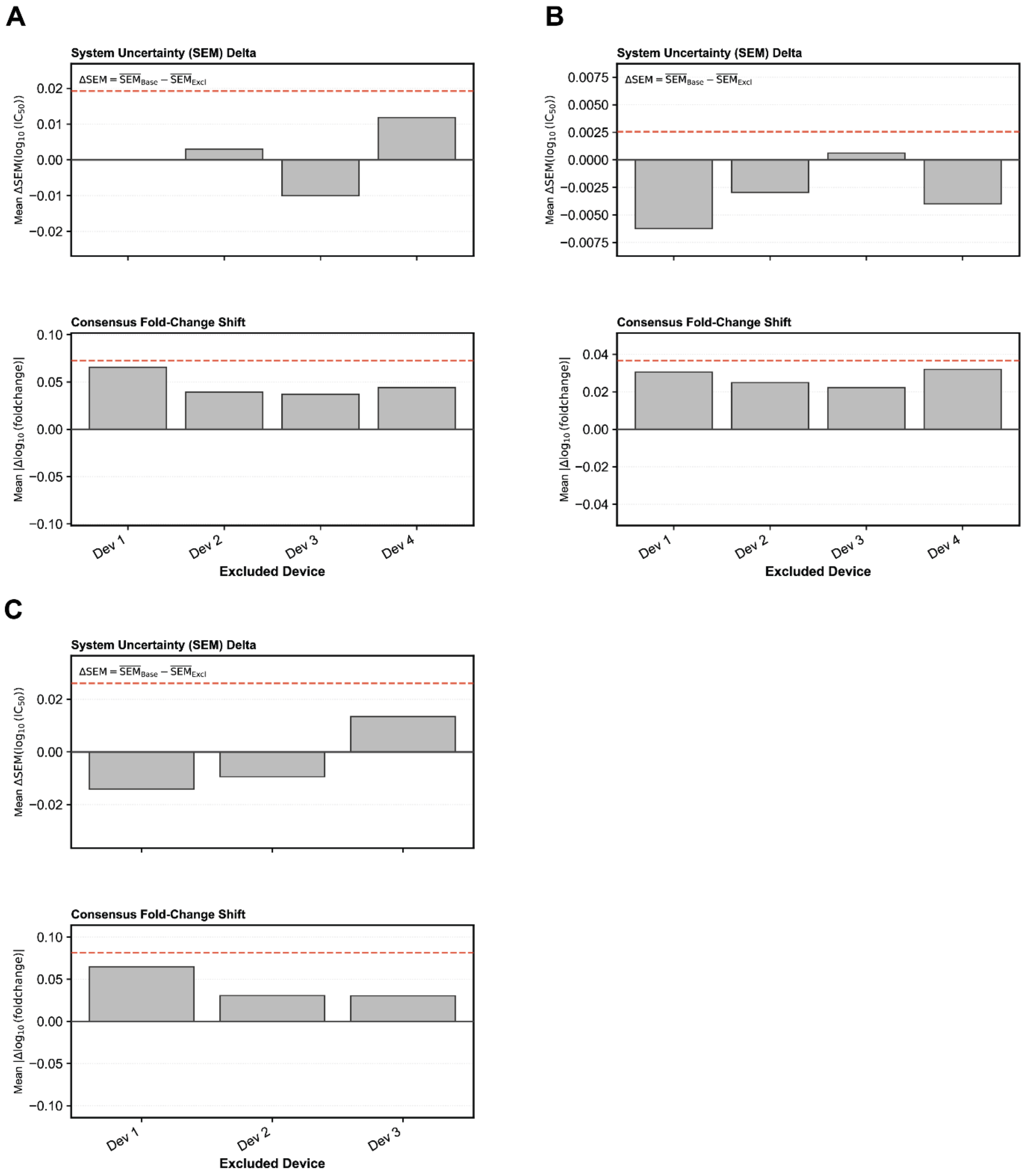


**Figure S41. Leave-one-device-out quality-control analysis of SHP2 TNO155 $IC_{50}$ measurements at 5, 50, and 500 nM ppIRS1.** Analysis for experiments at **(A)** 5 nM, **(B)** 50 nM, and **(C)** 500 nM ppIRS1. Within each panel, the top plot displays the change in system uncertainty, quantified as the difference in the mean standard error of the mean (SEM) of $\log_{10}(IC_{50})$ across variants when each device is excluded in turn relative to the global baseline including all devices. Positive values indicate reduced uncertainty upon exclusion of the device. The bottom plot displays the stability of consensus measurements assessed by the mean absolute shift across all variants in $\log_{10}$ fold change (difference in $\log_{10}(IC_{50})$ relative to an internal anchor variant) compared to the global consensus upon exclusion of each device. The anchor was chosen as the most consistently observed variant across devices. Larger values indicate greater deviation from the global consensus upon exclusion of the device. Dashed lines indicate mean + 2 SD thresholds. Because no single device exceeded the 2 SD threshold for both metrics, all devices were retained and none were excluded from the final analysis.

A

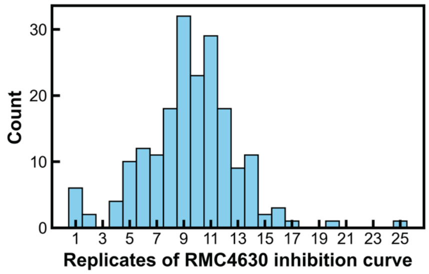


B

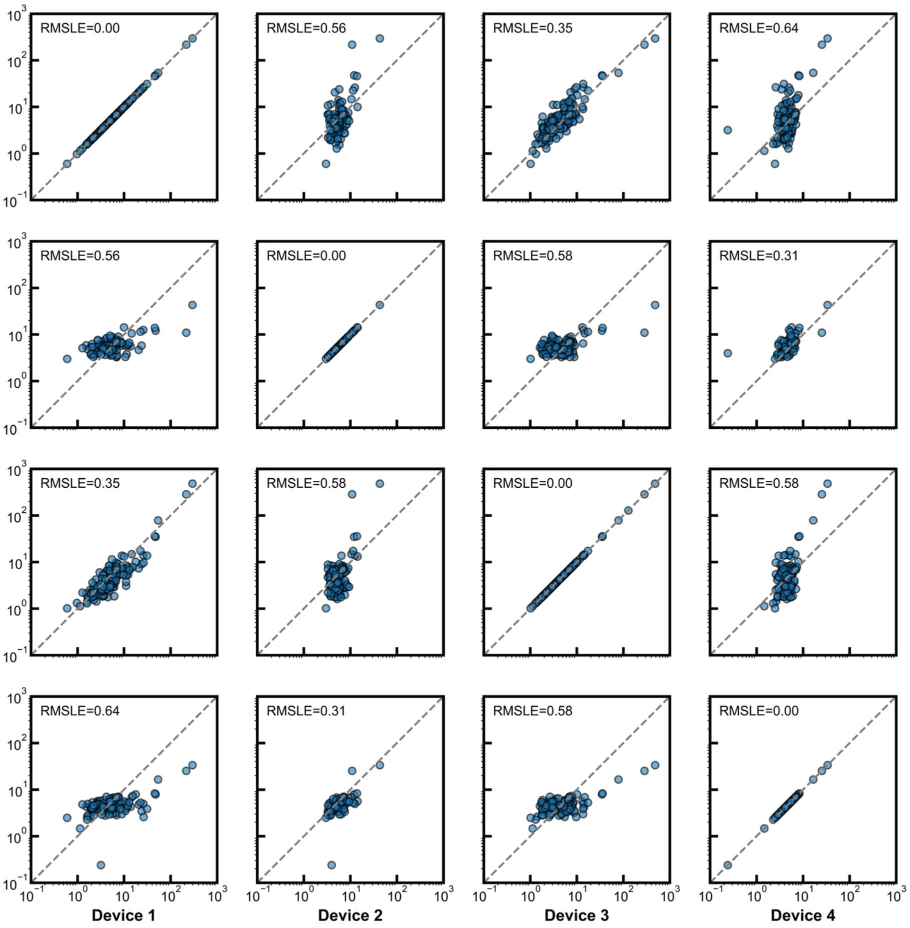


**Figure S42. Reproducibility of RMC-4630 $IC_{50}$ measurements. (A)** Histogram of RMC-4630 $IC_{50}$ measurement replicates across 190 $SHP2_{FL}$ variants. **(B)** Reproducibility of RMC-4630 $IC_{50}$ across device replicates. Each device reports the median of 1–4 replicates from different chambers for the same variant on the same chip. All values are in nM.

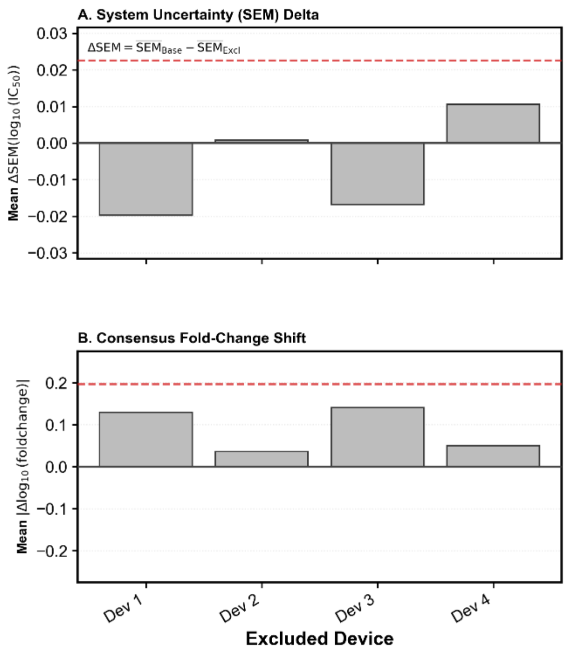


**Figure S43. Leave-one-device-out quality-control analysis of SHP2 RMC-4630 $IC_{50}$ measurements. (A)** Change in system uncertainty, quantified as the difference in the mean standard error of the mean (SEM) of $log_{10}(IC_{50})$ across variants when each device is excluded in turn relative to the global baseline including all devices. Positive values indicate reduced uncertainty upon exclusion of the device. **(B)** Stability of consensus measurements assessed by the mean absolute shift across all variants in $log_{10}$ fold change (difference in $log_{10}(IC_{50})$ relative to an internal anchor variant) compared to the global consensus upon exclusion of each device. The anchor was chosen as the most consistently observed variant across devices. Larger values indicate greater deviation from the global consensus upon exclusion of the device. Dashed lines indicate mean + 2 SD thresholds. Because no single device exceeded the 2 SD threshold for both metrics, all devices were retained and none were excluded from the final analysis.

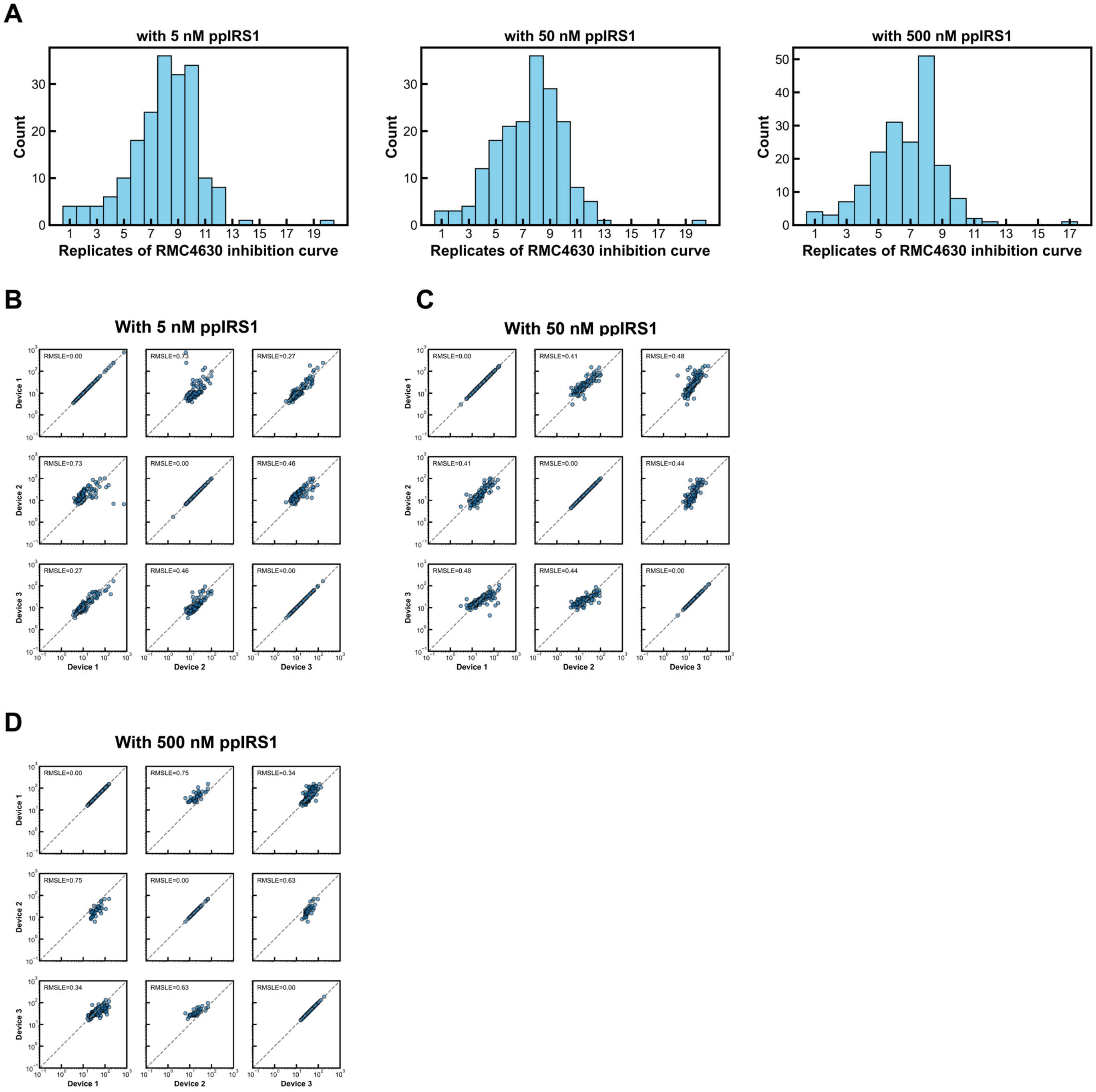


**Figure S44. Reproducibility of RMC-4630 $IC_{50}$ measurements under different concentrations of ppIRS1. (A)** Histogram of replicates for RMC-4630 $IC_{50}$ measurement under different concentrations of ppIRS1. Reproducibility of RMC-4630 $IC_{50}$ across device replicates under conditions with **(B)** 5 nM ppIRS1, **(C)** 50 nM ppIRS1, and **(D)** 500 nM ppIRS1. Each device reports the median of 1–4 replicates from different chambers for the same variant on the same chip. All values are in nM.

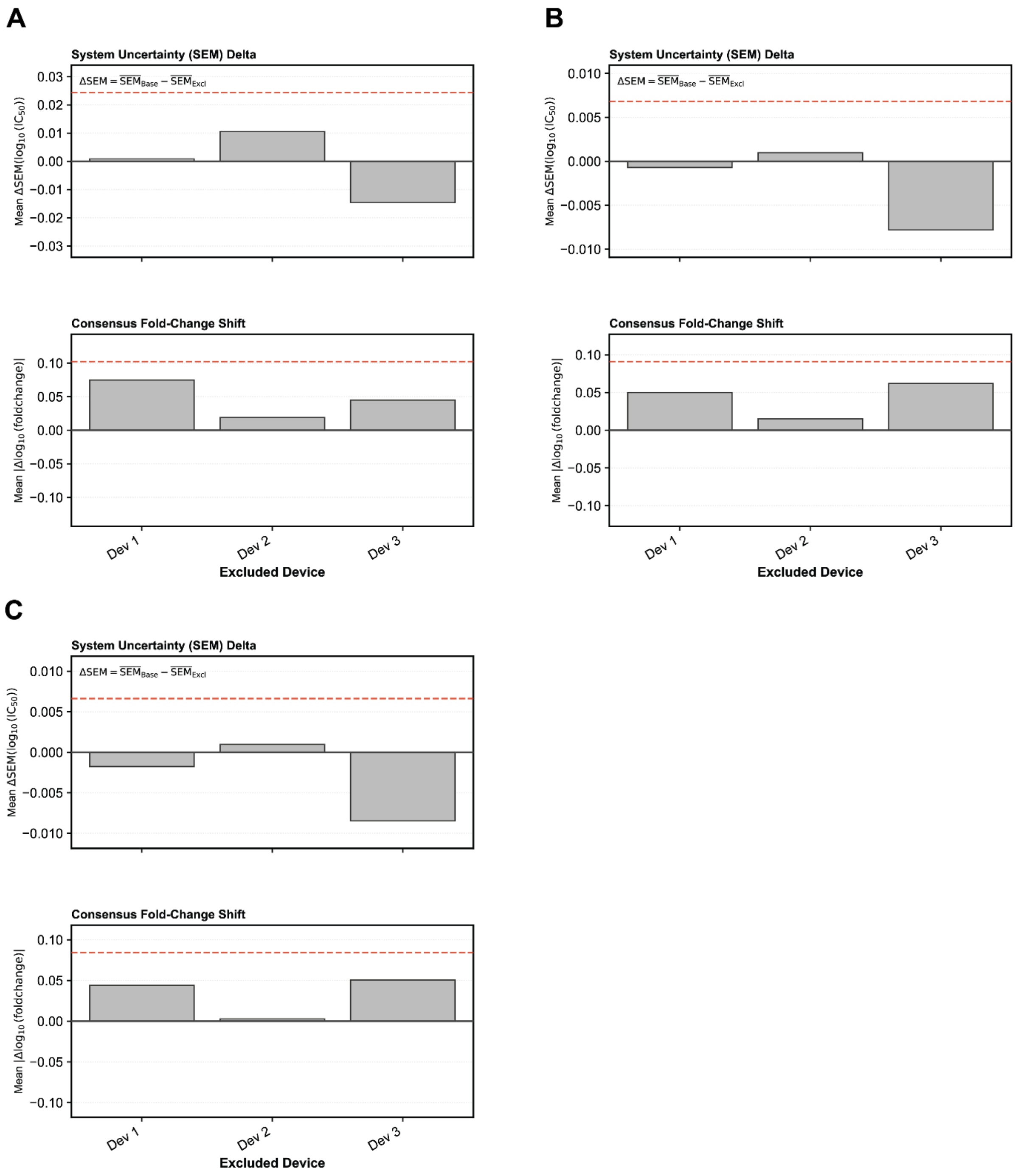


**Figure S45. Leave-one-device-out quality-control analysis of SHP2 RMC-4630 IC$_{50}$ measurements at 5, 50, and 500 nM ppIRS1.** Analysis for experiments at **(A)** 5 nM, **(B)** 50 nM, and **(C)** 500 nM ppIRS1. Within each panel, the top plot displays the change in system uncertainty, quantified as the difference in the mean standard error of the mean (SEM) of $\log_{10}$(IC$_{50}$) across variants when each device is excluded in turn relative to the global baseline including all devices. Positive values indicate reduced uncertainty upon exclusion of the device. The bottom plot displays the stability of consensus measurements assessed by the mean absolute shift across all variants in $\log_{10}$ fold change (difference in $\log_{10}$(IC$_{50}$) relative to an internal anchor variant) compared to the global consensus upon exclusion of each device. The anchor was chosen as the most consistently observed variant across devices. Larger values indicate greater deviation from the global consensus upon exclusion of the device. Dashed lines indicate mean + 2 SD thresholds. Because no single device exceeded the 2 SD threshold for both metrics, all devices were retained and none were excluded from the final analysis.

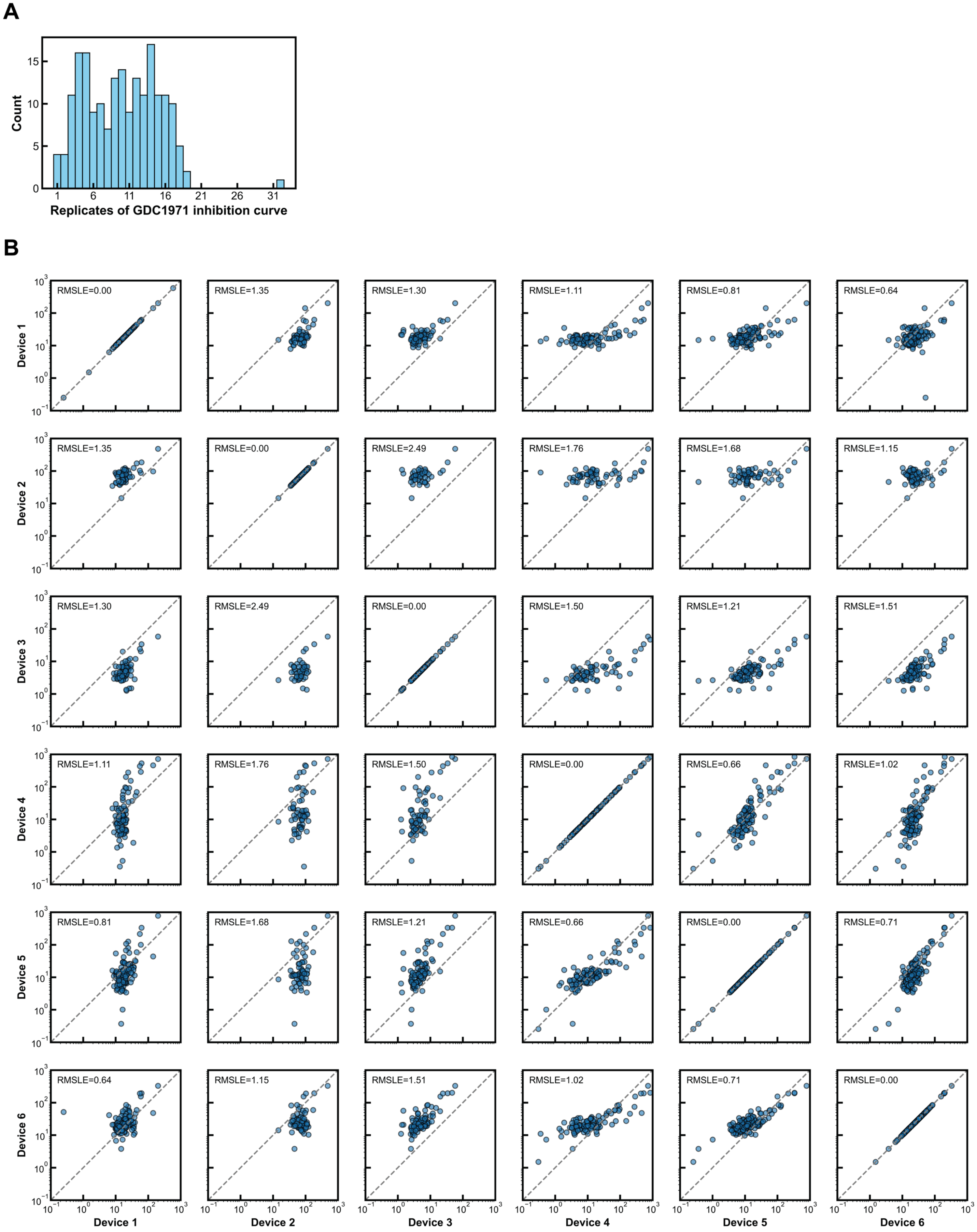


**Figure S46. Reproducibility of GDC-1971 $IC_{50}$ measurements. (A)** Histogram of GDC-1971 $IC_{50}$ measurement replicates across 190 $SHP2_{FL}$ variants. **(B)** Reproducibility of GDC-1971 $IC_{50}$ across device replicates. Each device reports the median of 1–4 replicates from different chambers for the same variant on the same chip. All values are in nM.

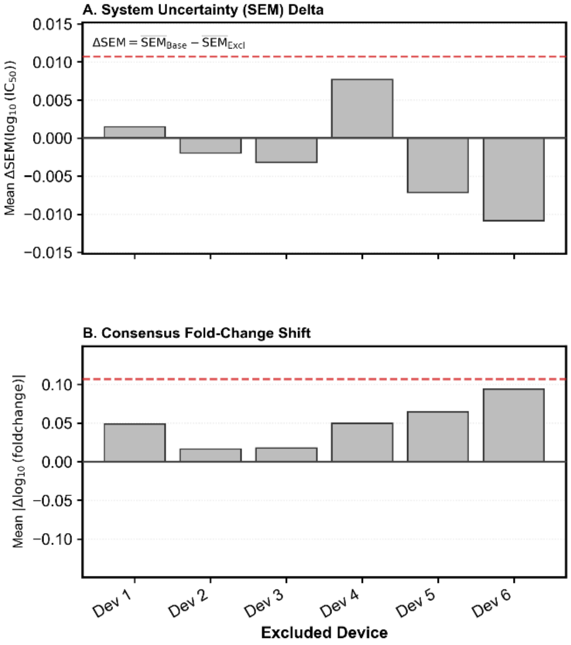


**Figure S47. Leave-one-device-out quality-control analysis of SHP2 GDC-1971 $IC_{50}$ measurements. (A)** Change in system uncertainty, quantified as the difference in the mean standard error of the mean (SEM) of $log_{10}(IC_{50})$ across variants when each device is excluded in turn relative to the global baseline including all devices. Positive values indicate reduced uncertainty upon exclusion of the device. **(B)** Stability of consensus measurements assessed by the mean absolute shift across all variants in $log_{10}$ fold change (difference in $log_{10}(IC_{50})$ relative to an internal anchor variant) compared to the global consensus upon exclusion of each device. The anchor was chosen as the most consistently observed variant across devices. Larger values indicate greater deviation from the global consensus upon exclusion of the device. Dashed lines indicate mean + 2 SD thresholds. Because no single device exceeded the 2 SD threshold for both metrics, all devices were retained and none were excluded from the final analysis.

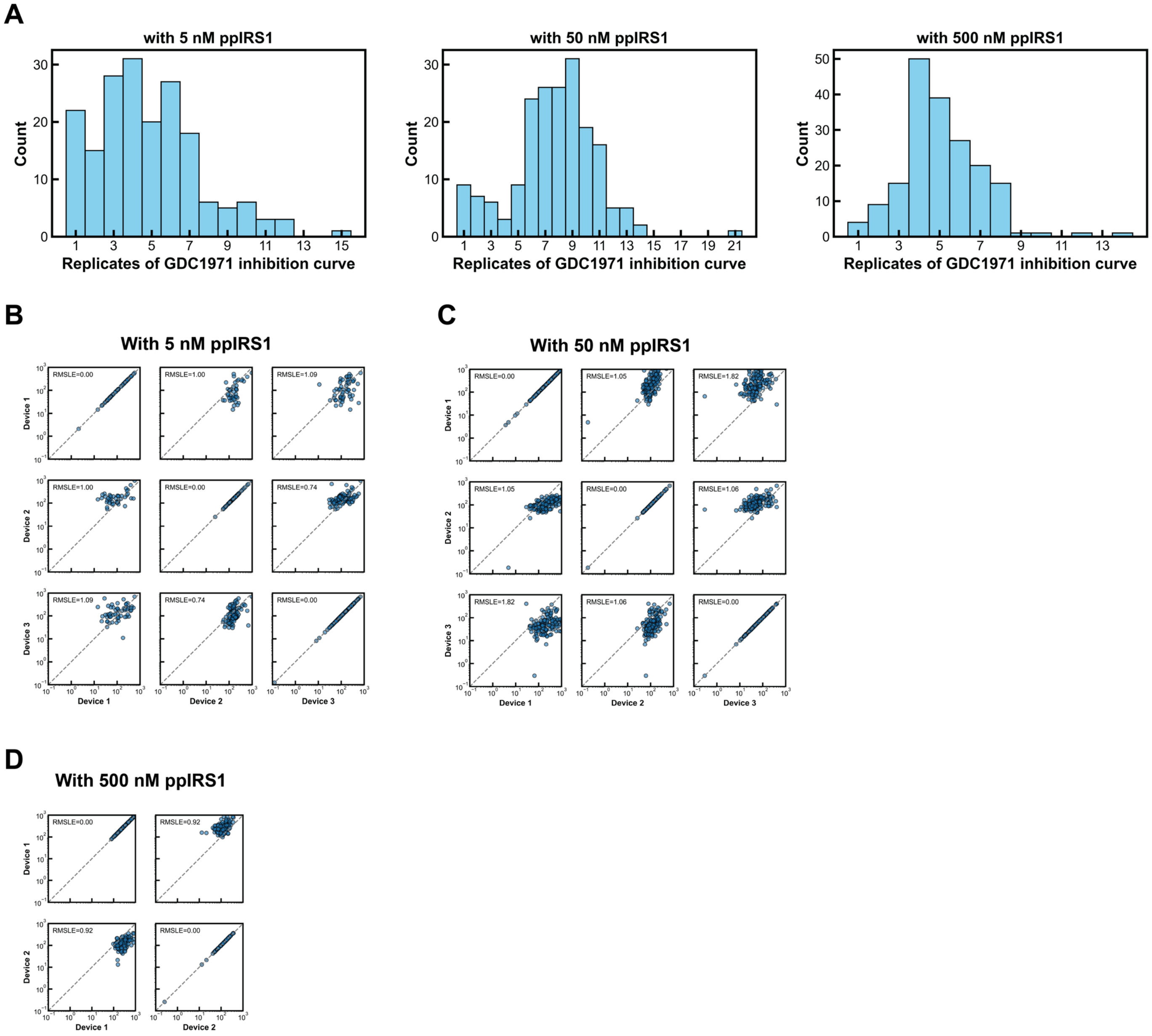


**Figure S48. Reproducibility of GDC-1971 $IC_{50}$ measurements under different concentrations of ppIRS1.** **(A)** Histogram of replicates for GDC-1971 $IC_{50}$ measurement under different concentrations of ppIRS1. Reproducibility of GDC-1971 $IC_{50}$ across device replicates under conditions with **(B)** 5 nM ppIRS1, **(C)** 50 nM ppIRS1, and **(D)** 500 nM ppIRS1. Each device reports the median of 1–4 replicates from different chambers for the same variant on the same chip. All values are in nM.

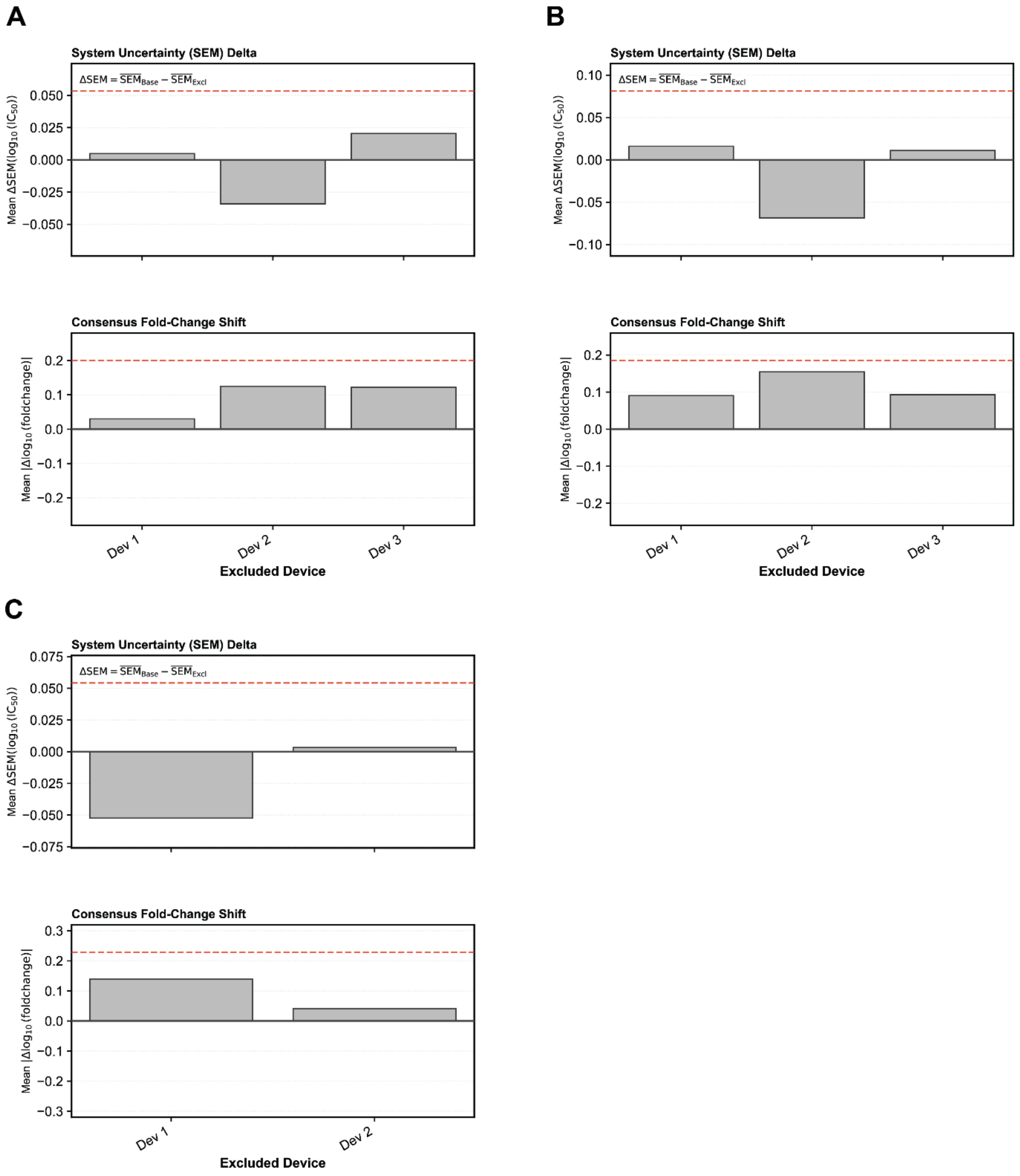


**Figure S49. Leave-one-device-out quality-control analysis of SHP2 GDC-1971 $IC_{50}$ measurements at 5, 50, and 500 nM ppIRS1.** Analysis for experiments at **(A)** 5 nM, **(B)** 50 nM, and **(C)** 500 nM ppIRS1. Within each panel, the top plot displays the change in system uncertainty, quantified as the difference in the mean standard error of the mean (SEM) of $\log_{10}(IC_{50})$ across variants when each device is excluded in turn relative to the global baseline including all devices. Positive values indicate reduced uncertainty upon exclusion of the device. The bottom plot displays the stability of consensus measurements assessed by the mean absolute shift across all variants in $\log_{10}$ fold change (difference in $\log_{10}(IC_{50})$ relative to an internal anchor variant) compared to the global consensus upon exclusion of each device. The anchor was chosen as the most consistently observed variant across devices. Larger values indicate greater deviation from the global consensus upon exclusion of the device. Dashed lines indicate mean + 2 SD thresholds. Because no single device exceeded the 2 SD threshold for both metrics, all devices were retained and none were excluded from the final analysis.

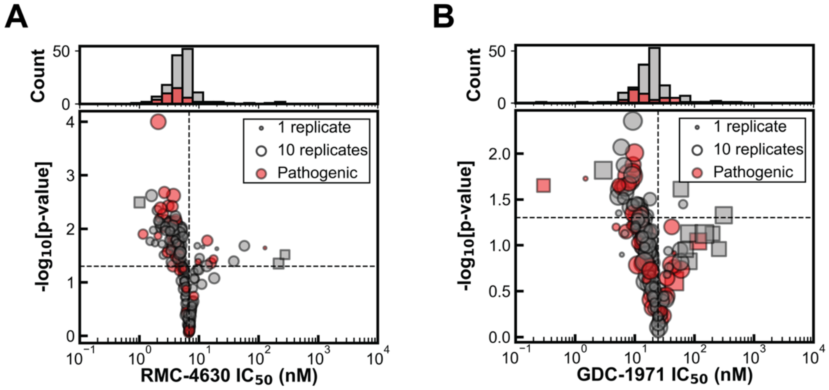


**Figure S50. Per-variant differences in measured $IC_{50}$ values for RMC-4630 and GDC-1971.** Median **(A)** RMC-4630 and **(B)** GDC-1971 $IC_{50}$ of 190 SHP2 variants and their significance of difference from WT (*p*-values). Red: Pathogenic variants in ClinVar.

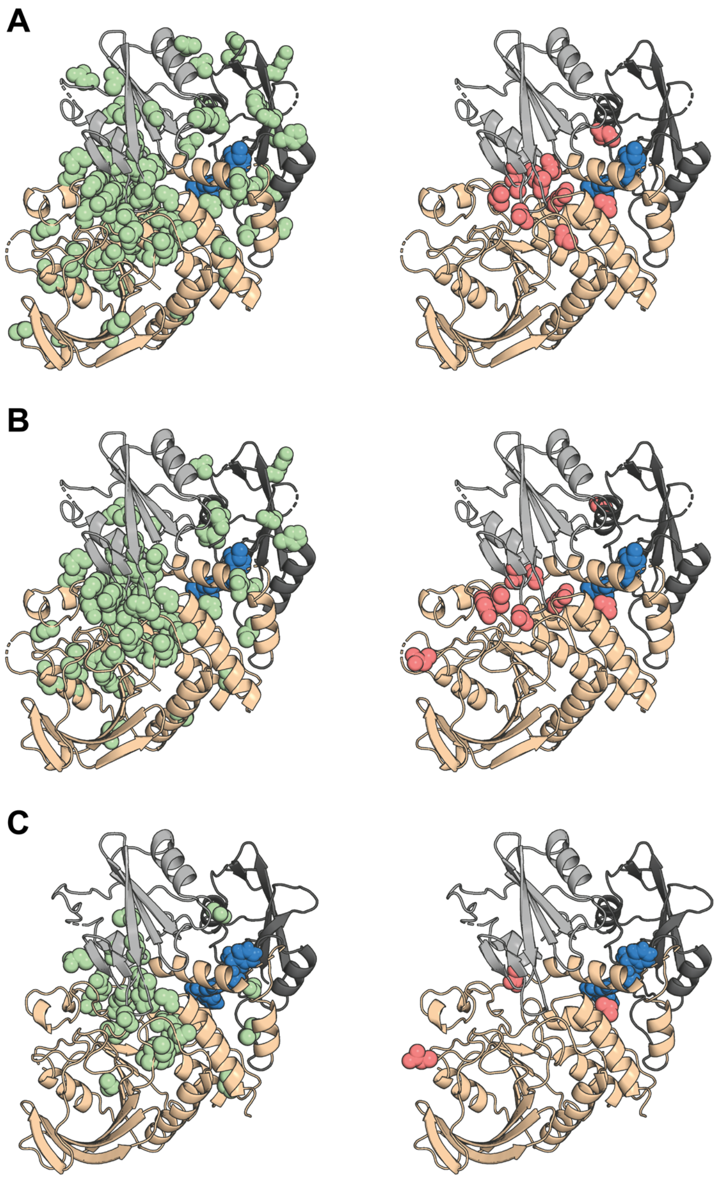


**Figure S51. Locations of variants with statistically significantly different $IC_{50}$ values from WT SHP2.** Positions with mutations that show a statistically significant difference from WT (bootstrap test $p < 0.05$), and more than 2-fold decrease (green) or increase (red) in $IC_{50}$ relative to WT, mapped onto SHP2 structures for **(A)** TNO155 (PDB ID: 7JVM), **(B)** RMC-4630 (illustrated using TNO155-bound structure due to lack of RMC-4630 co-crystal), and **(C)** GDC-1971 (PDB ID: 8T6D).

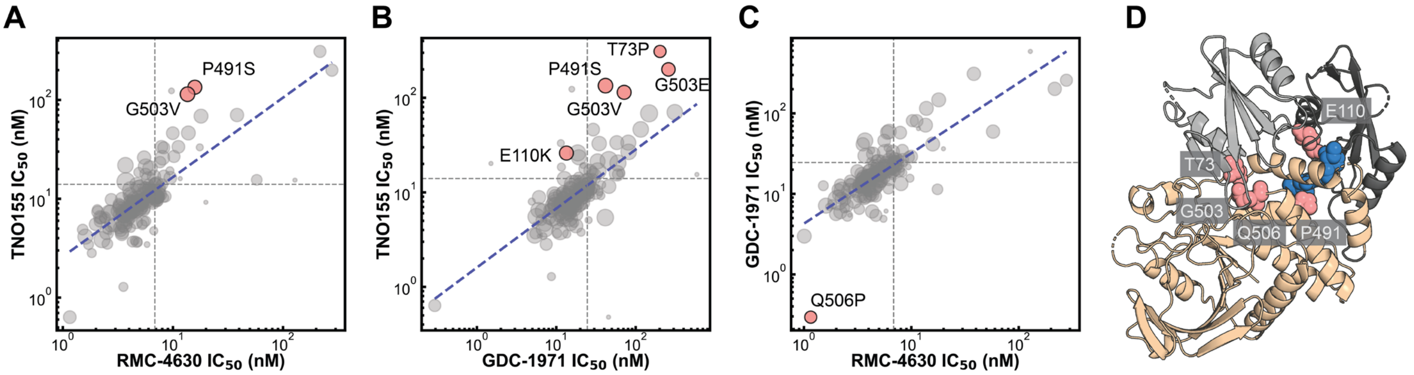


**Figure S52. Comparisons of $IC_{50}$ values measured for different inhibitors. (A)** TNO155 $IC_{50}$ versus RMC-4630 $IC_{50}$. **(B)** TNO155 $IC_{50}$ versus GDC-1971 $IC_{50}$. **(C)** GDC-1971 $IC_{50}$ versus RMC-4630 $IC_{50}$. In each plot, the blue dashed line represents the log–log linear regression. Outliers were defined as variants whose $IC_{50}$ values deviated ≥3-fold from regression-predicted values and passed significance testing (bootstrap test with Benjamini–Hochberg false discovery rate ≤ 0.05. **(D)** Structural location of SHP2 mutations that deviate from the overall correlation in sensitivity across allosteric inhibitors.

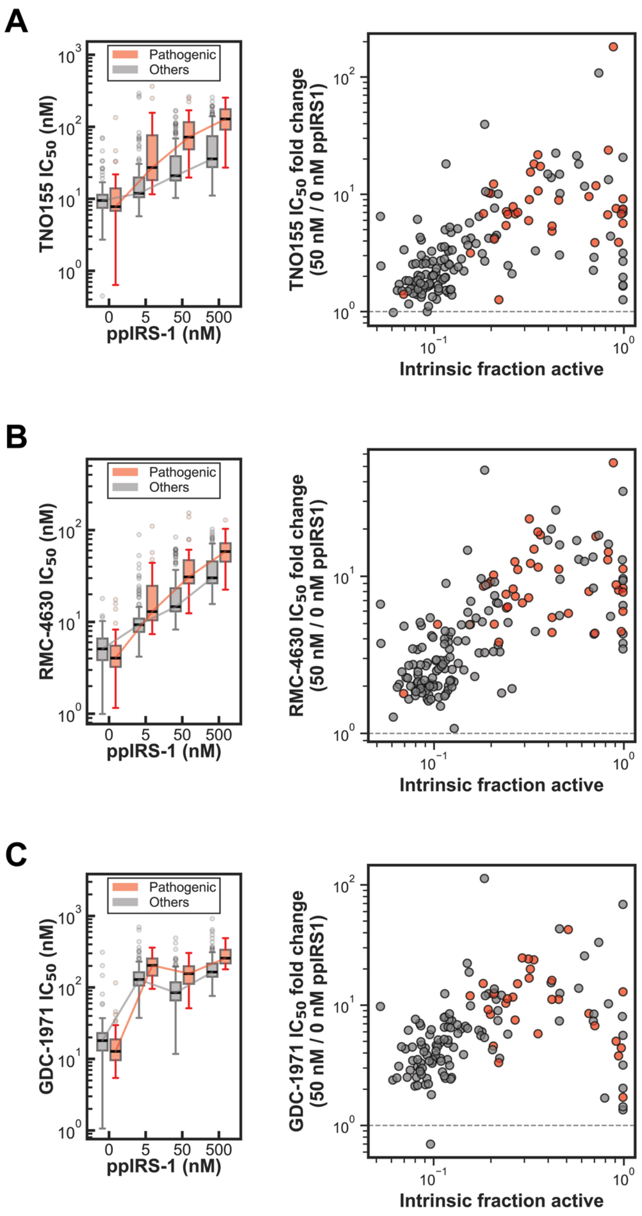


**Figure S53. Differences in $IC_{50}$ values as a function of activating peptide concentration across variants.** **(A)** Left: TNO155 $IC_{50}$ across different ppIRS1 concentrations. Pathogenic variants are marked in red. Right: TNO155 $IC_{50}$ fold change from 0 nM ppIRS1 to 50 nM ppIRS1 across different intrinsic fraction active. **(B)** Left: RMC-4630 $IC_{50}$ across different ppIRS1 concentrations. Pathogenic variants are marked in red. Right: RMC-4630 $IC_{50}$ fold change from 0 nM ppIRS1 to 50 nM ppIRS1 across different intrinsic fraction active. **(C)** Left: GDC-1971 $IC_{50}$ across different ppIRS1 concentrations. Pathogenic variants are marked in red. Right: GDC-1971 $IC_{50}$ fold change from 0 nM ppIRS1 to 50 nM ppIRS1 across different intrinsic fraction active.

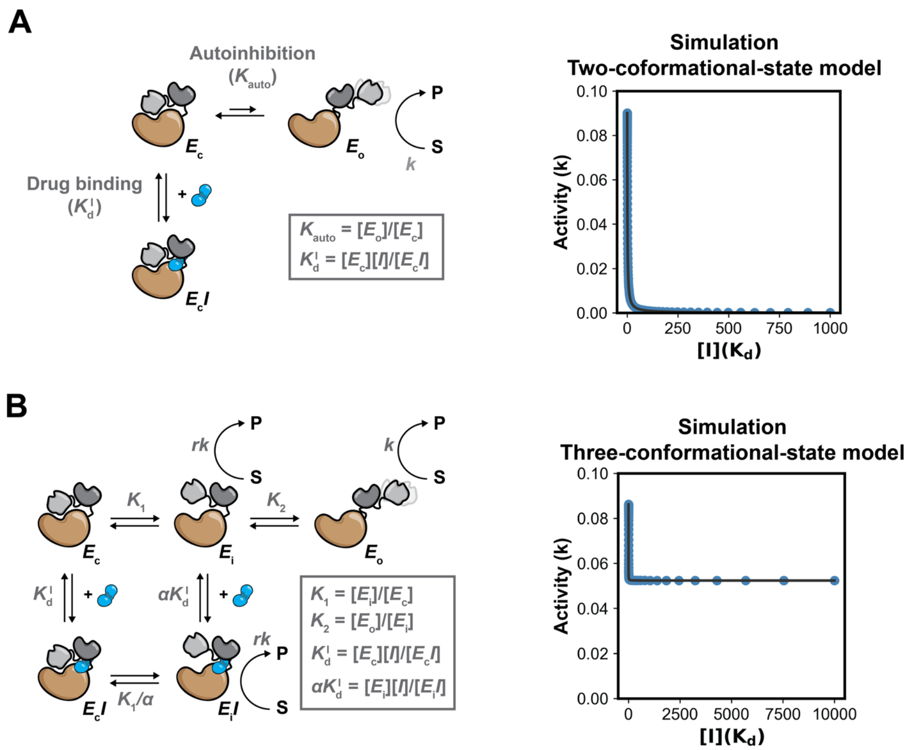


**Figure S54**. **Simulated SHP2 inhibition dose-response curves reveal differences in maximal inhibition depending on inhibition model. (A)** Simulated SHP2 inhibition dose-response curve with the canonical two-conformational-state model. Parameters: $K_{auto}$ = 0.1. **(B)** Simulated SHP2 inhibition dose-response curve with the three-conformational-state model. Parameters: $K_1$ = 0.11; $K_2$ = 0.85; $\alpha$ = 0.1; $r$ = 0.1. See Materials and Methods for details.

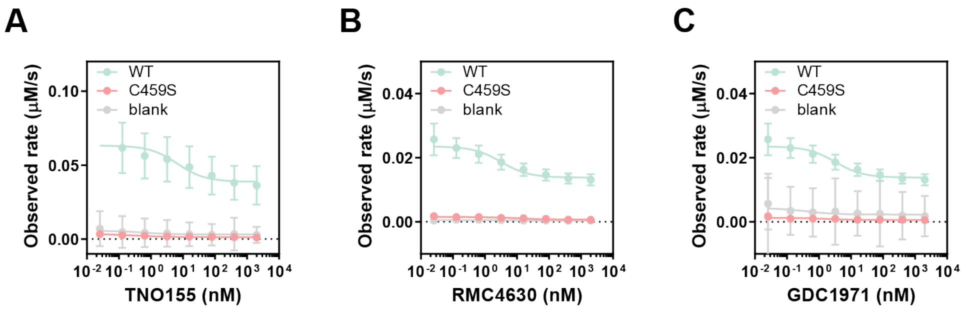


**Figure S55**. **High concentration of drug does not fully inhibit SHP2 hydrolysis of DiFMUP.** Dose-response curves displaying observed DiFMUP hydrolysis rates for WT, catalytically dead C459S, and control chambers with no variant expressed (blank) with varying concentrations of **(A)** TNO155, **(B)** RMC-4630, and **(C)** GDC-1971. Data were obtained via HT-MEK and represent mean ± SEM of the observed rate. Results indicate that even at saturating drug concentrations, WT SHP2 maintains a residual DiFMUP hydrolysis activity.

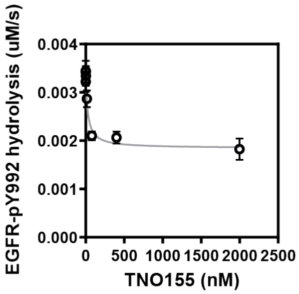


**Figure S56**. **High concentration of drug does not fully inhibit SHP2 hydrolysis of a peptide substrate.** TNO155 inhibition of SHP2 WT dose-response curve for EGFRpY992 hydrolysis. The assay was performed at a sub-saturating substrate concentration of 50 μM EGFRpY992.

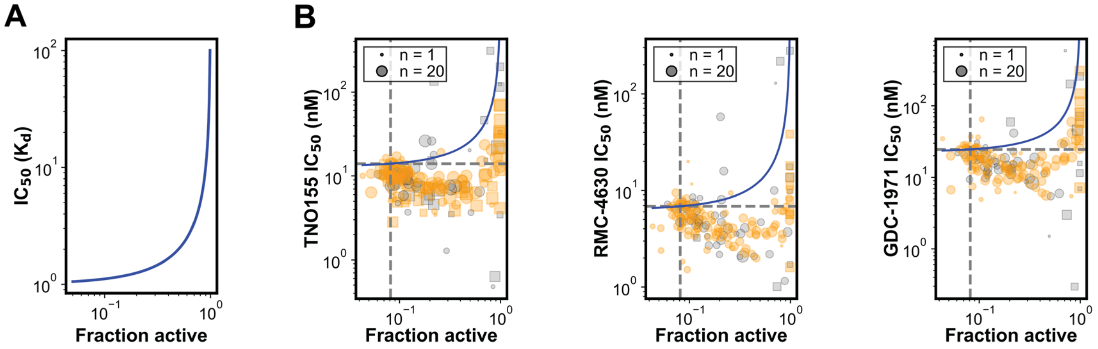


**Figure S57. Comparison between two-conformational-state model predictions and experimental data for the relationship between measured $IC_{50}$ and fraction active. (A)** Two-conformational-state inhibition model (**Fig. S36A**) predicted $IC_{50}$ versus $f^0_{active}$ relationship (**Eqn. 2**). **(B)** TNO155, RMC-4630, GDC-1971 $IC_{50}$ versus $f^0_{active}$. Blue line represents two-conformational-state model predictions (**Eqn. 2**), assuming all variants bind the drug with a constant intrinsic dissociation constant $K^I_d$ equal to that of the WT. Gold: variants with mutation > 15 Å away from the drug. Variants with $IC_{50}$ measurements outside the range of inhibitor concentrations used are plotted in squares. Marker size is proportional to $IC_{50}$ replicates. Grey dash lines show WT values.

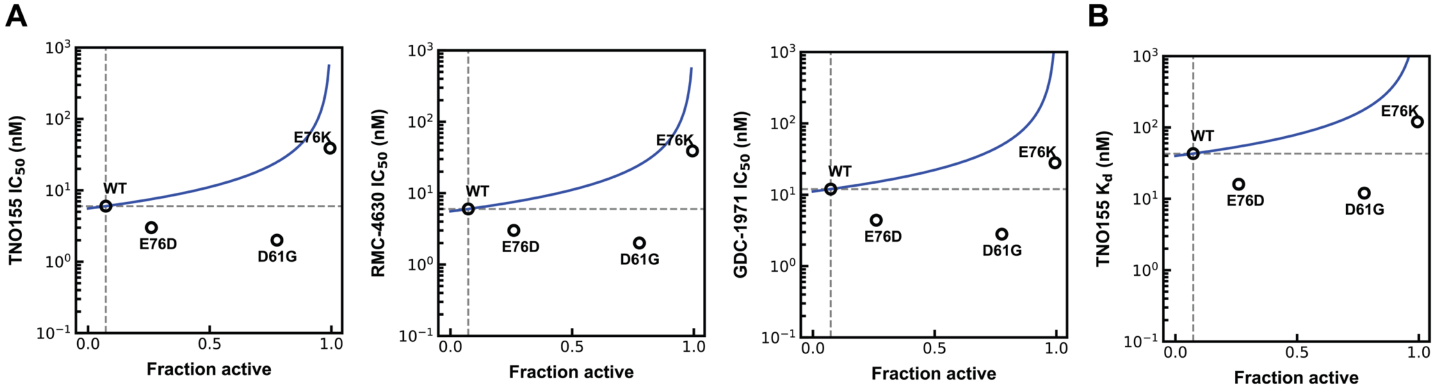


**Figure S58. $IC_{50}$ measured via plate reader assay and microscale thermophoresis versus fraction active for 4 different mutants across 3 drugs. (A)** Plate reader assay measured TNO155 $IC_{50}$, RMC-4630 $IC_{50}$, GDC-1971 $IC_{50}$ versus $f_{active}^{0}$. **(B)** Microscale thermophoresis measured TNO155 $K_d$ versus $f_{active}^{0}$. Blue line represents two-conformational-state model predictions (**Eqn. 2**), assuming all variants bind the drug with a constant dissociation constant $K_d^I$ equal to that of the WT. Grey dash lines show WT values.

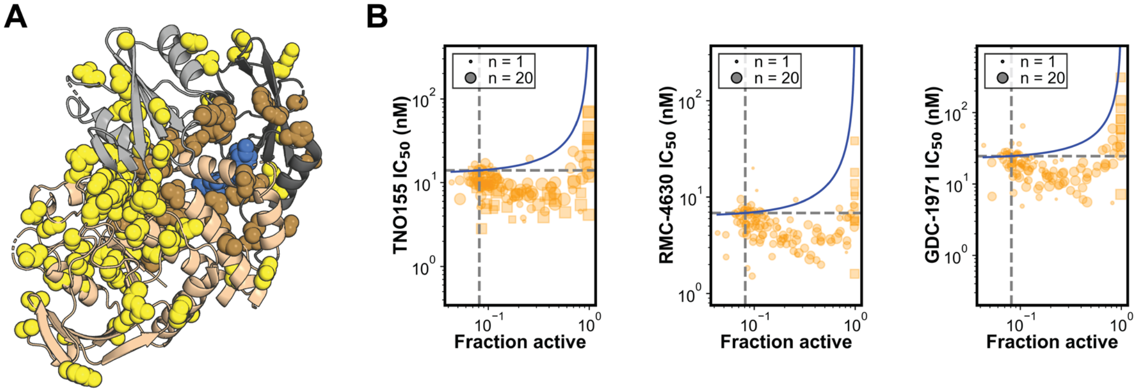


**Figure S59. Variants far from the drug binding site with increased fraction active are more sensitive to drug. (A)** Structural mapping of mutated residues relative to the drug. All mutation sites in the library are shown as spheres, with the drug in spheres as well (blue). Residues within 15 Å of the drug (63/189) are colored brown; those >15 Å away (126/189) are yellow. **(B)** TNO155, RMC-4630, and GDC-1971 $IC_{50}$ versus $f_{active}^{0}$ scatter plot with variants with mutation over 15 Å away from the drug. Variants with $IC_{50}$ measurements outside the range of inhibitor concentrations used are plotted in squares. Marker size is proportional to $IC_{50}$ replicates. Blue line represents two-conformational-state model predictions (**Eqn. 2**), assuming all variants bind the drug with a constant intrinsic dissociation constant $K_d^I$ equal to that of the WT. Grey dash lines show WT values.

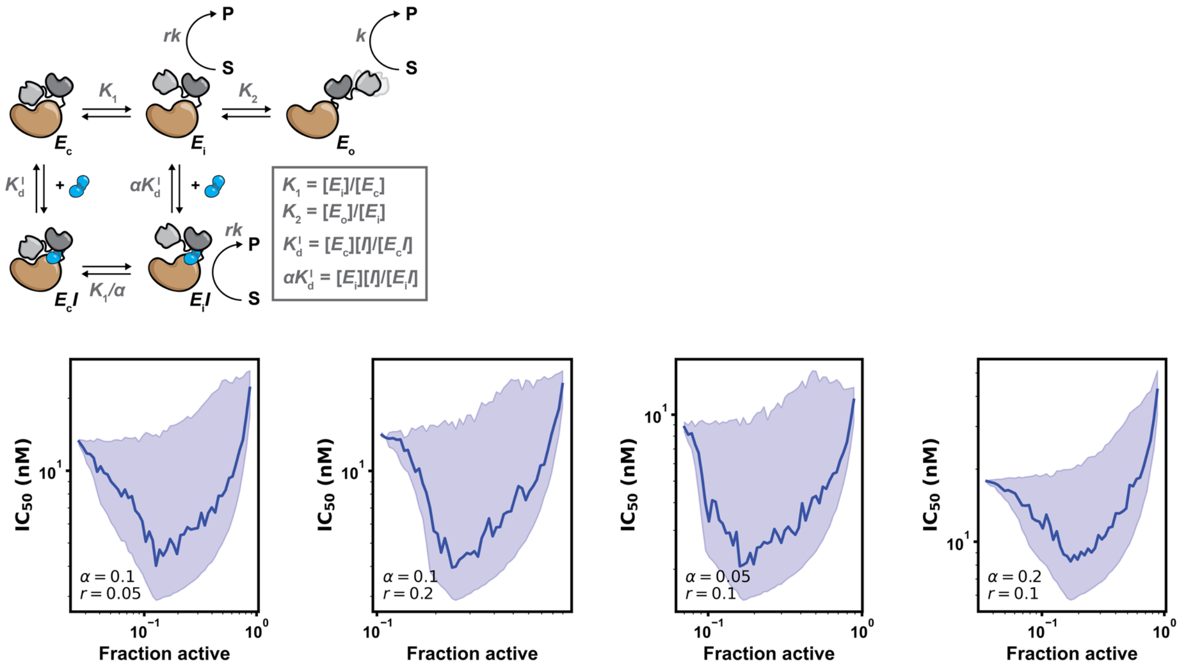


**Figure S60. Parameter sensitivity analysis of the three-conformational-state inhibition model.** Simulated relationships between fraction active and $IC_{50}$ were computed across a four-fold range of $\alpha$ and $r$ values ($\alpha$ = 0.05, 0.1, 0.2; $r$ = 0.05, 0.1, 0.2). $K_1$ and $K_2$ were randomly sampled. Solid lines show the median and shaded regions indicate the 95% interval from 10,000 simulations within logarithmically spaced fraction active bins. See **Materials and Methods** for details.

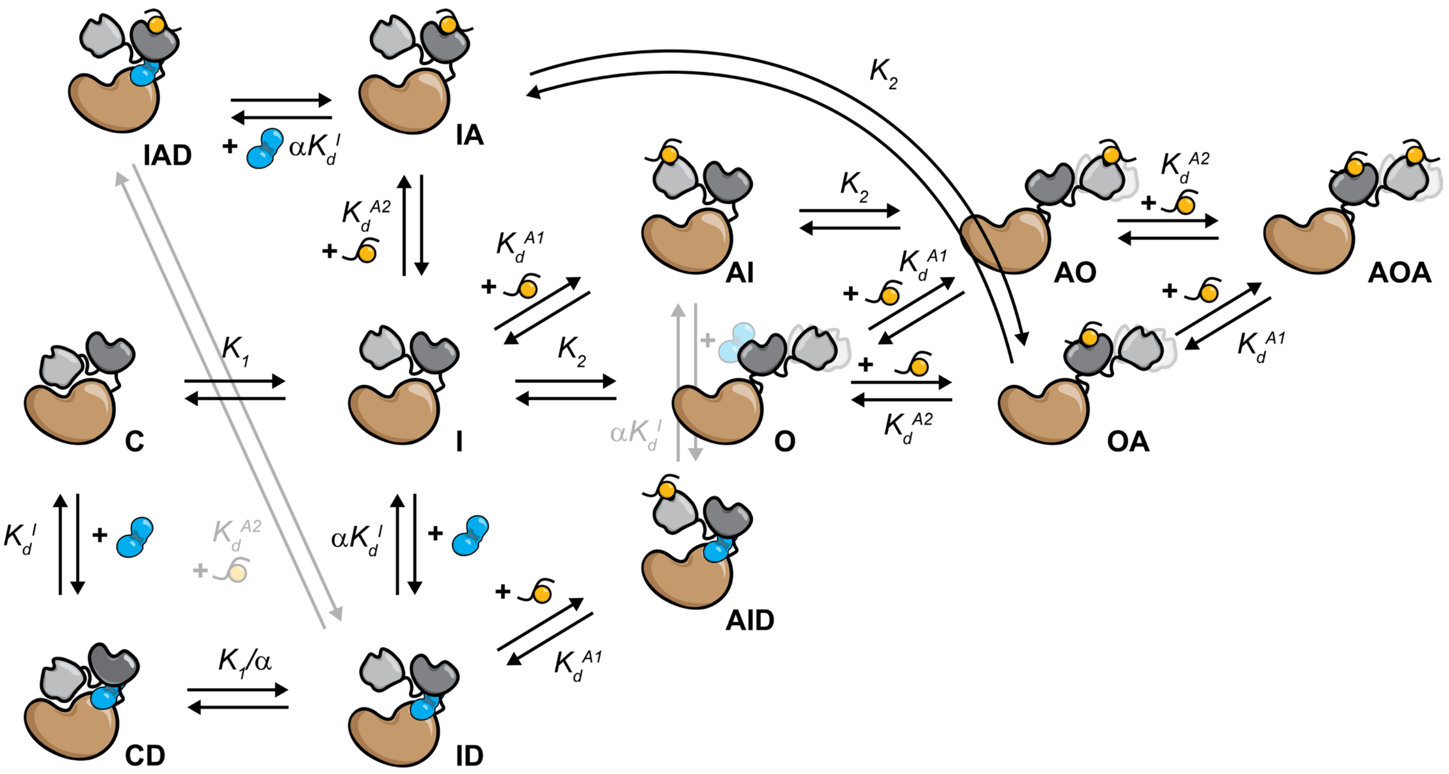


**Figure S61**. **Schematic of a three-conformational-state thermodynamic model of SHP2 activator binding and drug inhibition.** Here, *C*, *I*, and *O* represent the apo closed, intermediate, and open enzyme states, respectively. States bound to the activator (*A*) are denoted by the addition of *A*. Binding to the nSH2 domain is represented by an *A* in front of the state (*e.g.*, *AI*, *AO*), while binding to the secondary site (*e.g.*, cSH2) is represented by an *A* following the state (*e.g.*, *IA*, *OA*). The fully occupied open state is denoted as *AOA*. States bound to the inhibitor (*D*) are denoted by the addition of *D* (*e.g.*, *CD*, *ID*, *AID*, *IAD*).

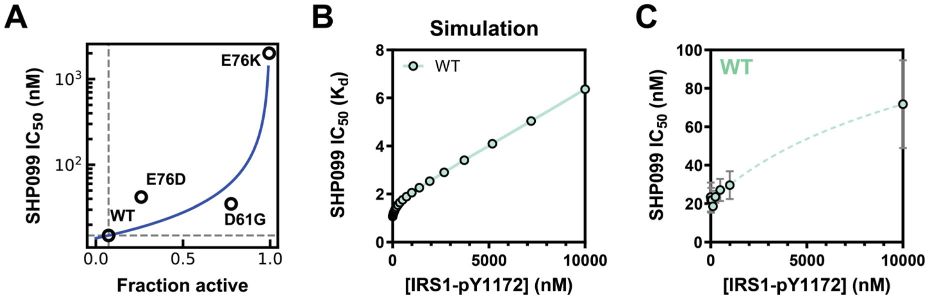


**Figure S62**. **Three-conformational-state model explains SHP099 $IC_{50}$ as a function of activating peptide concentration. (A)** Plate reader assay measured SHP099 $IC_{50}$ versus $f^0_{active}$. **(B)** Simulated SHP099 $IC_{50}$ dependence on IRS1-pY1172 concentration. **(C)** Experimental data of SHP099 $IC_{50}$ dependence on IRS1-pY1172 concentration.

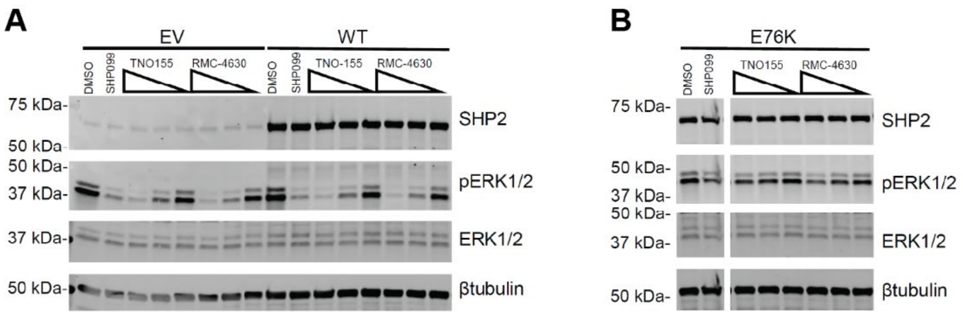


**Figure S63. Assessing cellular SHP2 inhibition assay. (A)** Representative Western blot of U2OS cells transfected with empty vector (EV) or WT SHP2, treated with TNO155 or RMC-4630 (0.1, 1, 10 µM). Controls include DMSO and 50 µM SHP099. **(B)** Representative Western blot of U2OS cells transfected with the drug-resistant mutant E76K, treated as in **(A)**.

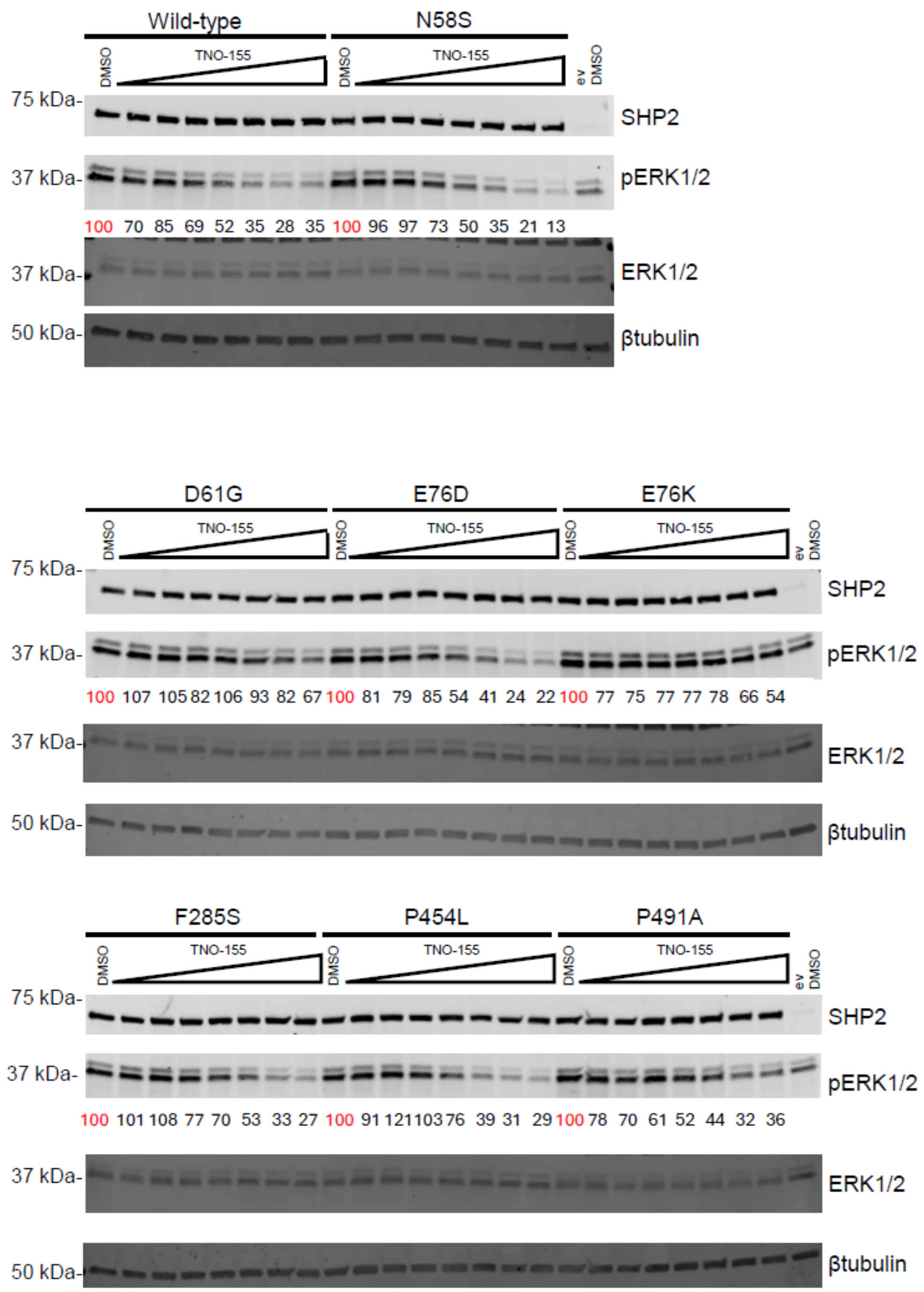


**Figure S64. TNO155 dose-response in SHP2 variant-transfected U2OS cells.** Representative immunoblots of SHP2, pERK1/2, and ERK1/2 in U2OS cells expressing SHP2 variants, treated with DMSO or TNO155 (0.01, 0.03, 0.1, 0.3, 1, 3, 10 μM). Numbers denote the normalized pERK1/2 level. Empty vector (ev) was included as a control.

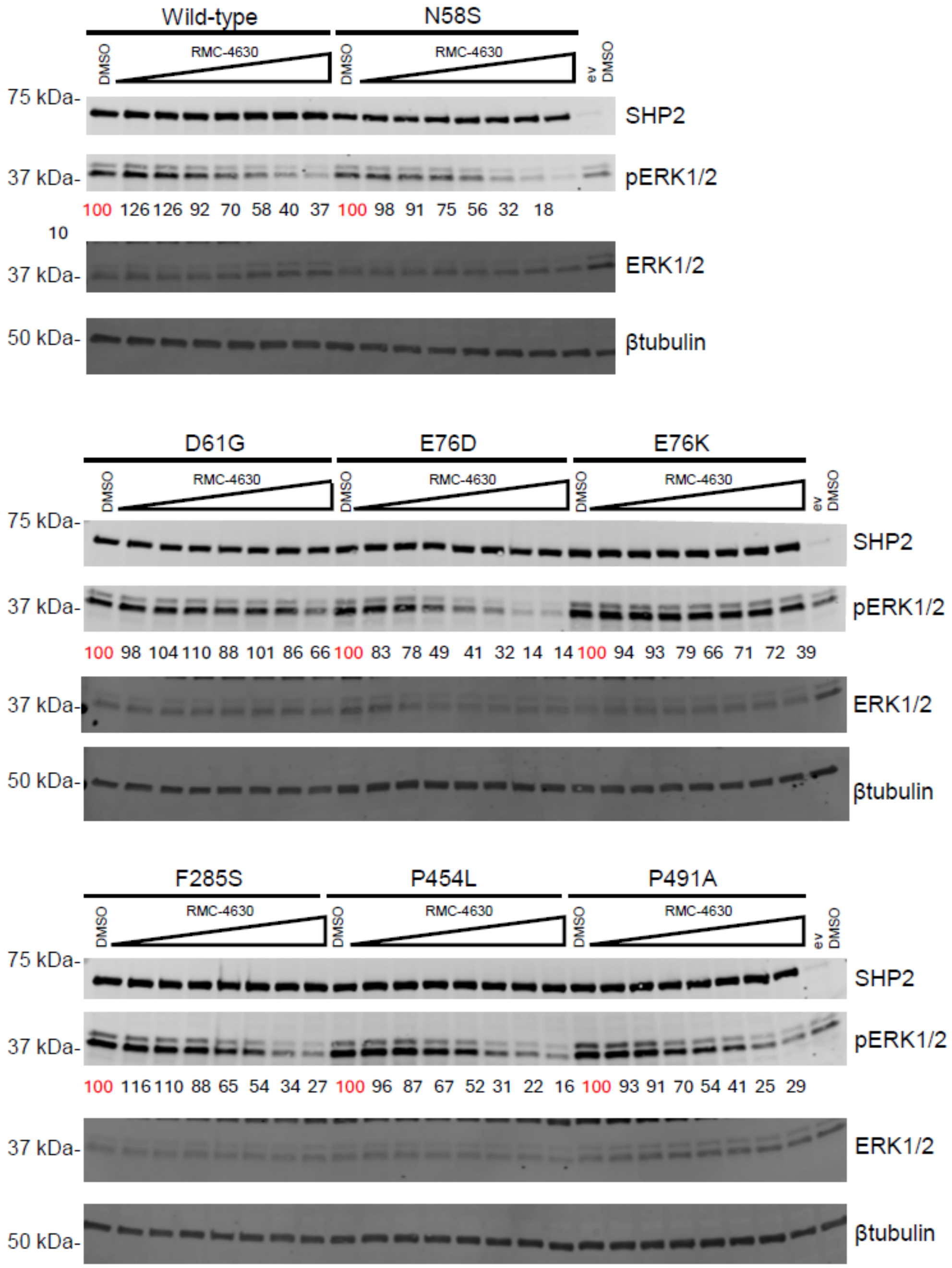


**Figure S65. RMC-4630 dose-response in SHP2 variant-transfected U2OS cells.** Representative immunoblots of SHP2, pERK1/2, and ERK1/2 in U2OS cells expressing SHP2 variants, treated with DMSO or RMC-4630 (0.01, 0.03, 0.1, 0.3, 1, 3, 10 µM). Numbers denote the normalized pERK1/2 level. Empty vector (ev) was included as a control.

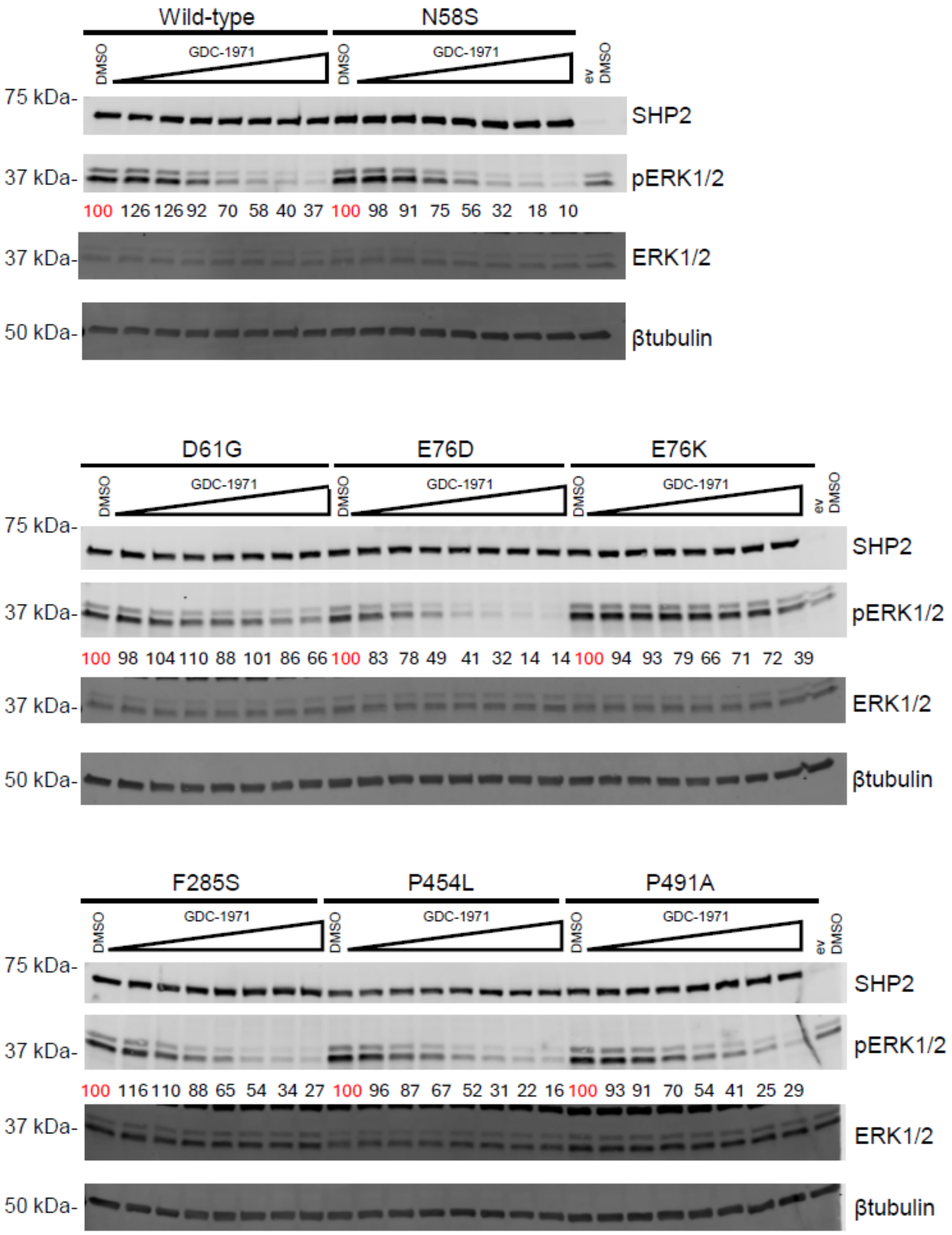


**Figure S66. GDC-1971 dose-response in SHP2 variant-transfected U2OS cells.** Representative immunoblots of SHP2, pERK1/2, and ERK1/2 in U2OS cells expressing SHP2 variants, treated with DMSO or GDC-1971 (0.01, 0.03, 0.1, 0.3, 1, 3, 10 µM). Numbers denote the normalized pERK1/2 level quantified by densitometry. Empty vector (ev) was included as a control.

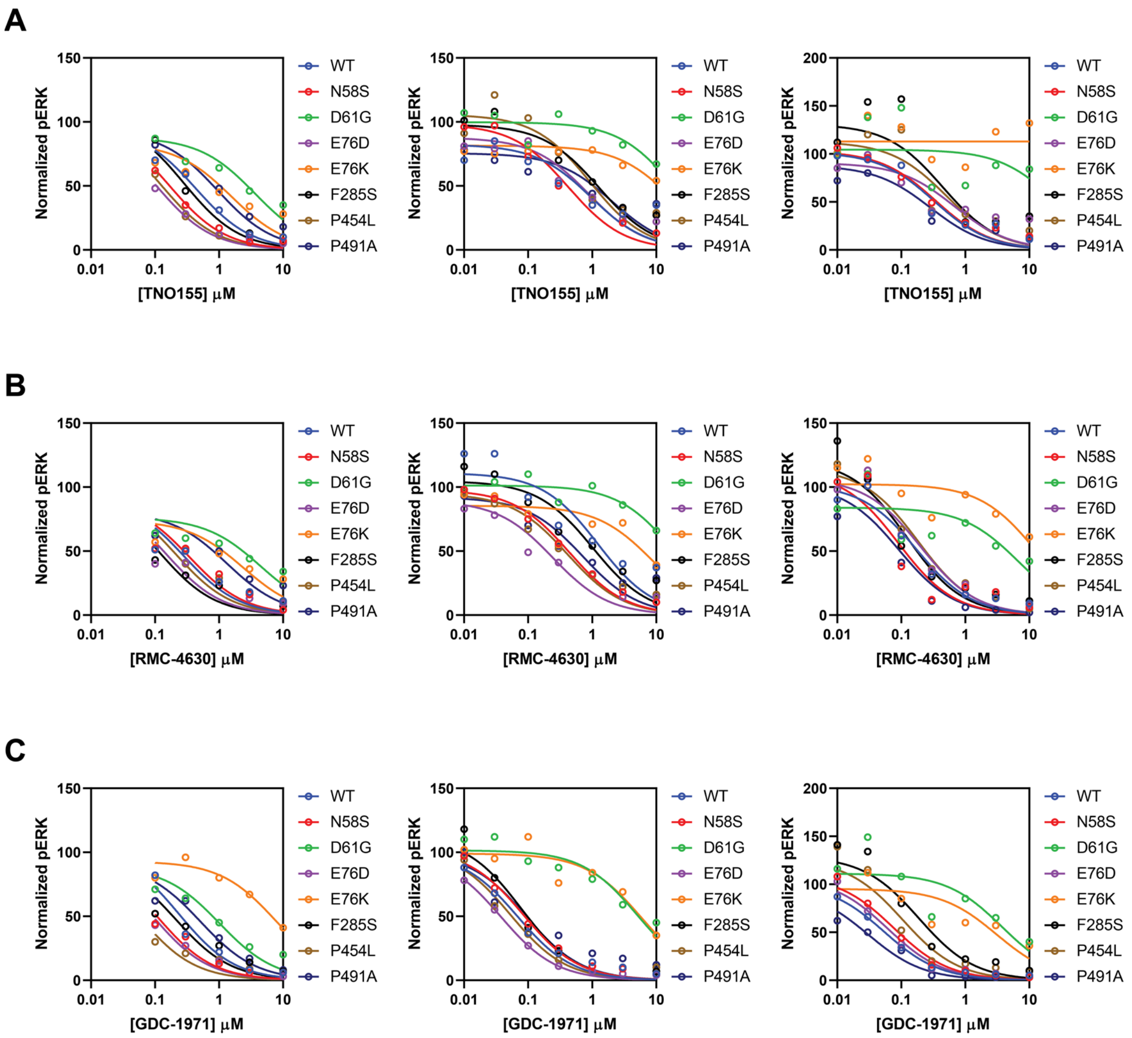


**Figure S67. Dose-dependent reduction of pERK level by SHP2 inhibitors in variant-transfected U2OS cells.** Normalized pERK1/2 levels in U2OS cells transfected with the indicated SHP2 variants as a function of inhibitors. Cells were treated with increasing concentrations of **(A)** TNO155, **(B)** RMC-4630, or **(C)** GDC-1971. The three plots per inhibitor were from three independent biological replicates. Data were fitted using a three-parameter logistic regression model (standard slope) with the lower limit fixed to 0.

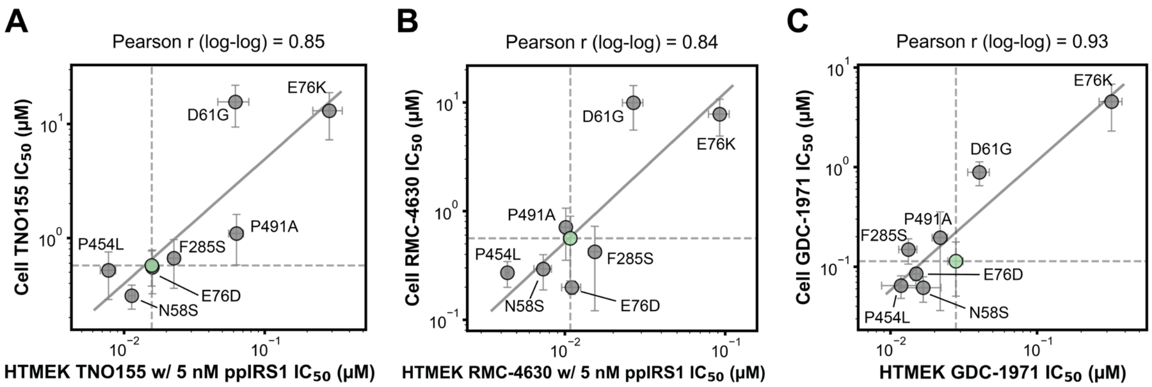


**Figure S68. Correlation between biochemical and cellular sensitivities for allosteric SHP2 inhibitors.** Comparison of $IC_{50}$ values obtained from HTMEK assays versus cellular assays for **(A)** TNO155, **(B)** RMC-4630, and **(C)** GDC-1971 across a panel of SHP2 variants. Data points represent the mean $IC_{50}$ for WT (green) and seven mutants (grey) from at least three independent replicates. Error bars represent SEM. Dashed lines indicate WT values. The solid line represents the linear regression fit on log-transformed data. The Pearson correlation coefficient calculated from the log-log data is indicated above each panel.

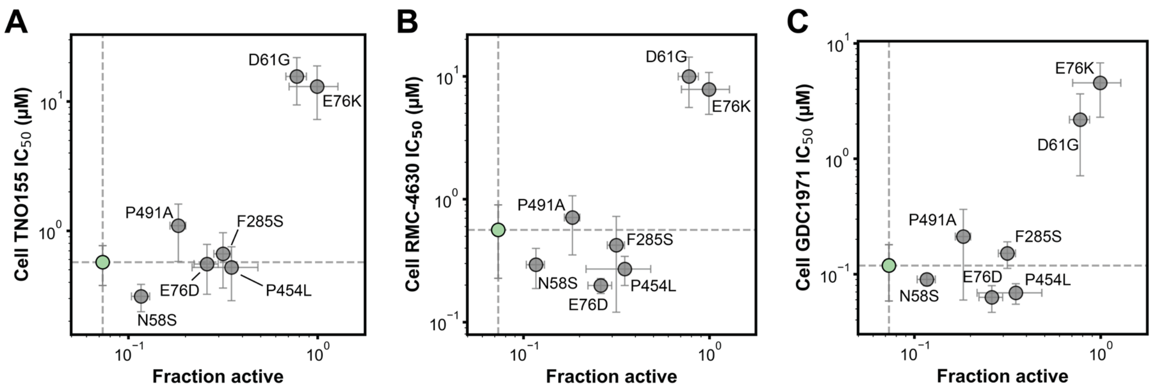


**Figure S69. Dependence of cellular sensitivity to allosteric inhibition on the fraction active of SHP2 variants.** Relationship between the fraction active and cellular $IC_{50}$ values for **(A)** TNO155, **(B)** RMC-4630, and **(C)** GDC-1971 across SHP2 variants. WT SHP2 is highlighted in green. Dashed lines indicate WT values. Data shown as mean ± SEM. Variants with moderate fraction active generally display lower or similar $IC_{50}$ compared to WT. Variants with high fraction active display higher $IC_{50}$ compared to WT. P491A is a mutation at the drug binding site.

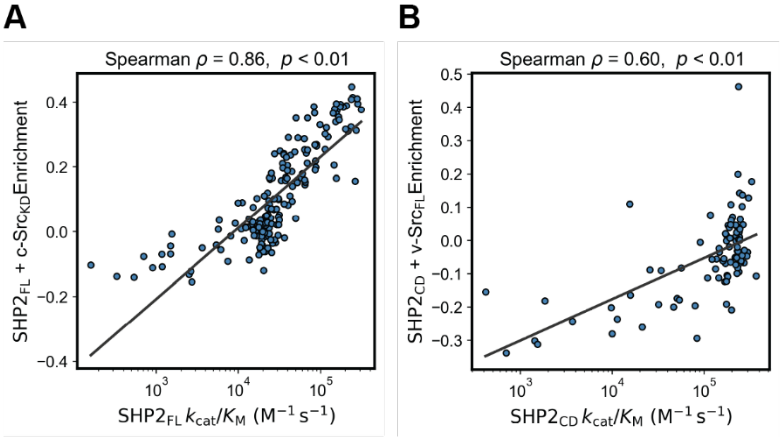


**Figure S70**. **Comparison of deep mutational scanning enrichment scores for SHP2 variants from the literature to $k_{cat}/K_M$ measured in this study.** The enrichment scores are derived from a yeast growth-rescue assay in which exogenously expressed Src kinases inhibits proliferation and more active SHP2 variants are expected to better counter this toxicity (*8*). **(A)** Enrichment score of $SHP2_{FL}$ variants from literature versus $k_{cat}/K_M$ measured in this study. There is a strong positive correlation between the two metrics (Spearman $\rho$ = 0.86, $p < 0.01$). **(B)** Enrichment score of $SHP2_{CD}$ variants from literature versus $k_{cat}/K_M$ measured in this study. There is a moderate positive correlation between the two metrics (Spearman $\rho$ = 0.60, $p < 0.01$).

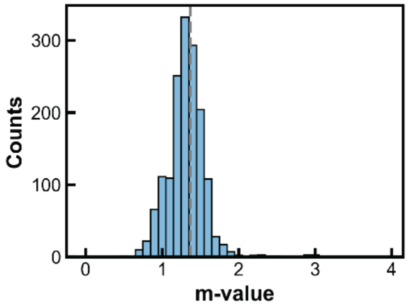


**Figure S71. Histogram of fitted m-values for $SHP2_{CD}$ variants.** The median m-value is 1.38 kcal/mol/M (95% CI: 1.36–1.40 kcal/mol/M)

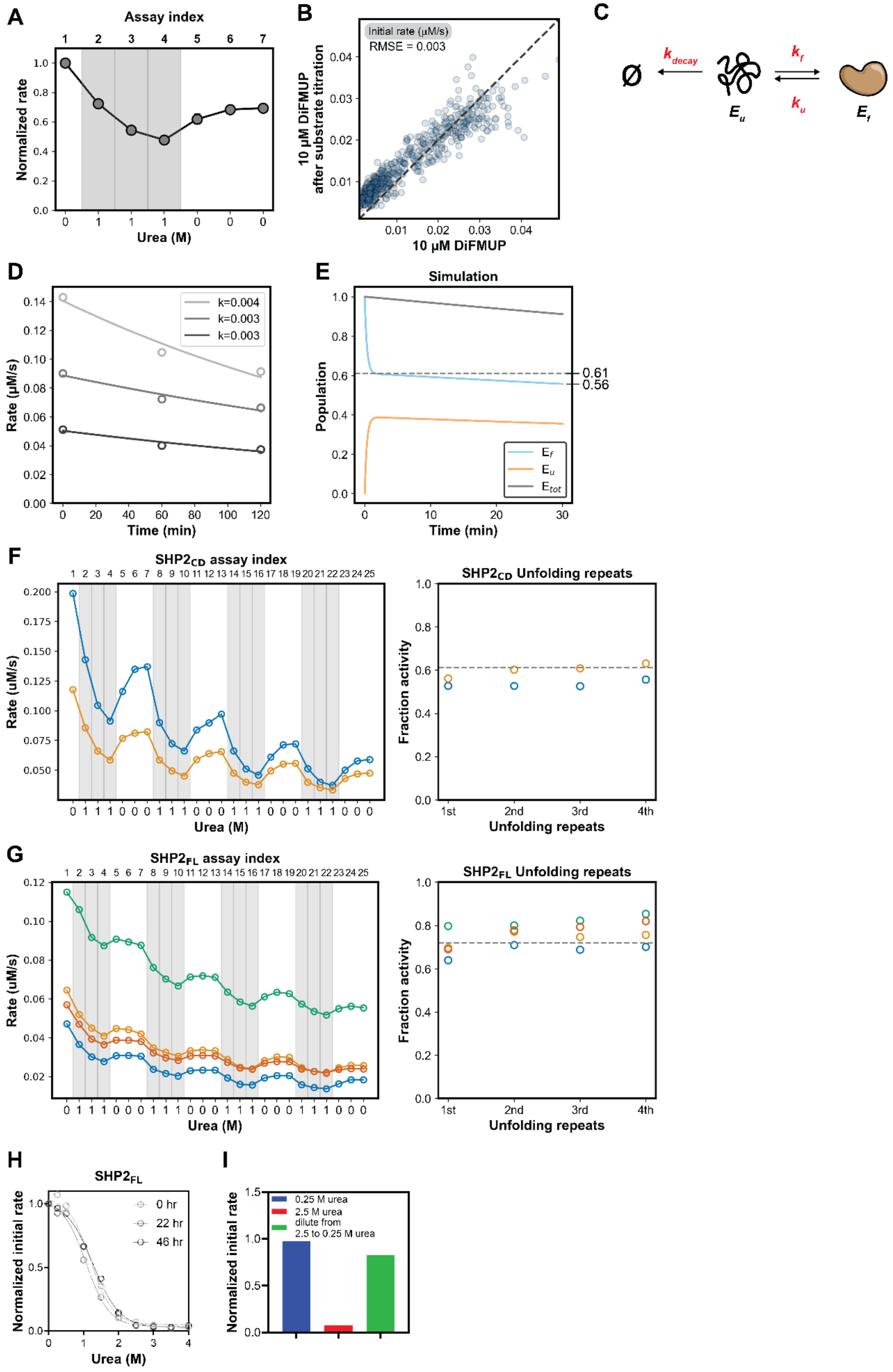


**Figure S72. Optimization of unfolding assay on HT-MEK. (A)** Normalized measured rates of WT $SHP2_{CD}$-mediated DiFMUP hydrolysis over iterative HT-MEK assays in the presence and absence of 1 M urea. “Assay index” denotes the sequential order of the assays performed; fitted initial rates of DiFMUP turnover were normalized to the initial rate of the first assay (assay index 1). **(B)** Scatter plot of measured initial rates of

DiFMUP hydrolysis for a library of $SHP2_{CD}$ variants measured after *vs.* before a series of HT-MEK substrate turnover assays (after a total of about 8 hr). **(C)** Schematic of a two-state folding model with irreversible inactivation from the unfolded state. **(D)** Fitted rates of urea-dependent irreversible inactivation for $SHP2_{CD}$ catalyzed DiFMUP hydrolysis over time at 1 M urea. Each time point represents a kinetic assay in which reaction products were washed away and fresh substrate was introduced while maintaining constant 1 M urea. Assays were conducted at 1-hour intervals. Shades of gray indicate three independent replicates. **(E)** Simulation of time-dependent changes in $E_F$, $E_U$, and $E_{tot}$ $(E_F + E_U)$ following a step change from 0 M to 1 M urea at t=0. **(F)** Sequentially-measured initial rates of WT $SHP2_{CD}$ DiFMUP hydrolysis for kinetic assays performed in the absence and presence of 1 M urea using HT-MEK. Residual activity at 1 M urea was calculated relative to the immediately preceding 0 M urea measurement and is plotted on the right. Gray dashed line indicates the fractional activity at 1 M urea measured using a plate reader. **(G)** Sequentially-measured initial rates of WT $SHP2_{FL}$ DiFMUP hydrolysis for kinetic assays performed in the absence and presence of 1 M urea using HT-MEK. Residual activity at 1 M urea was calculated relative to the immediately preceding 0 M urea measurement and is plotted on the right. Gray dashed line indicates the fractional activity at 1 M urea measured using a plate reader. **(H)** Representative urea dose-response curves for $SHP2_{FL}$ measured after varying preincubation times with urea. **(I)** Retained activity after treating with 0.25 M urea for 46 hr, 2.5 M urea for 46 hr, and 2.5 M urea for 46 hr then dilute to 0.25 M urea.

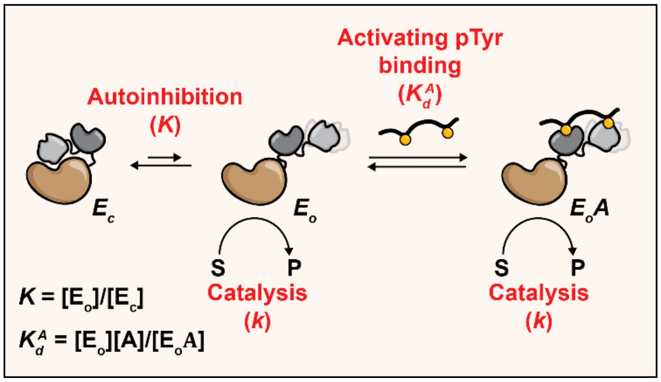


**Figure S73. Schematic of the model for SHP2 activation.**

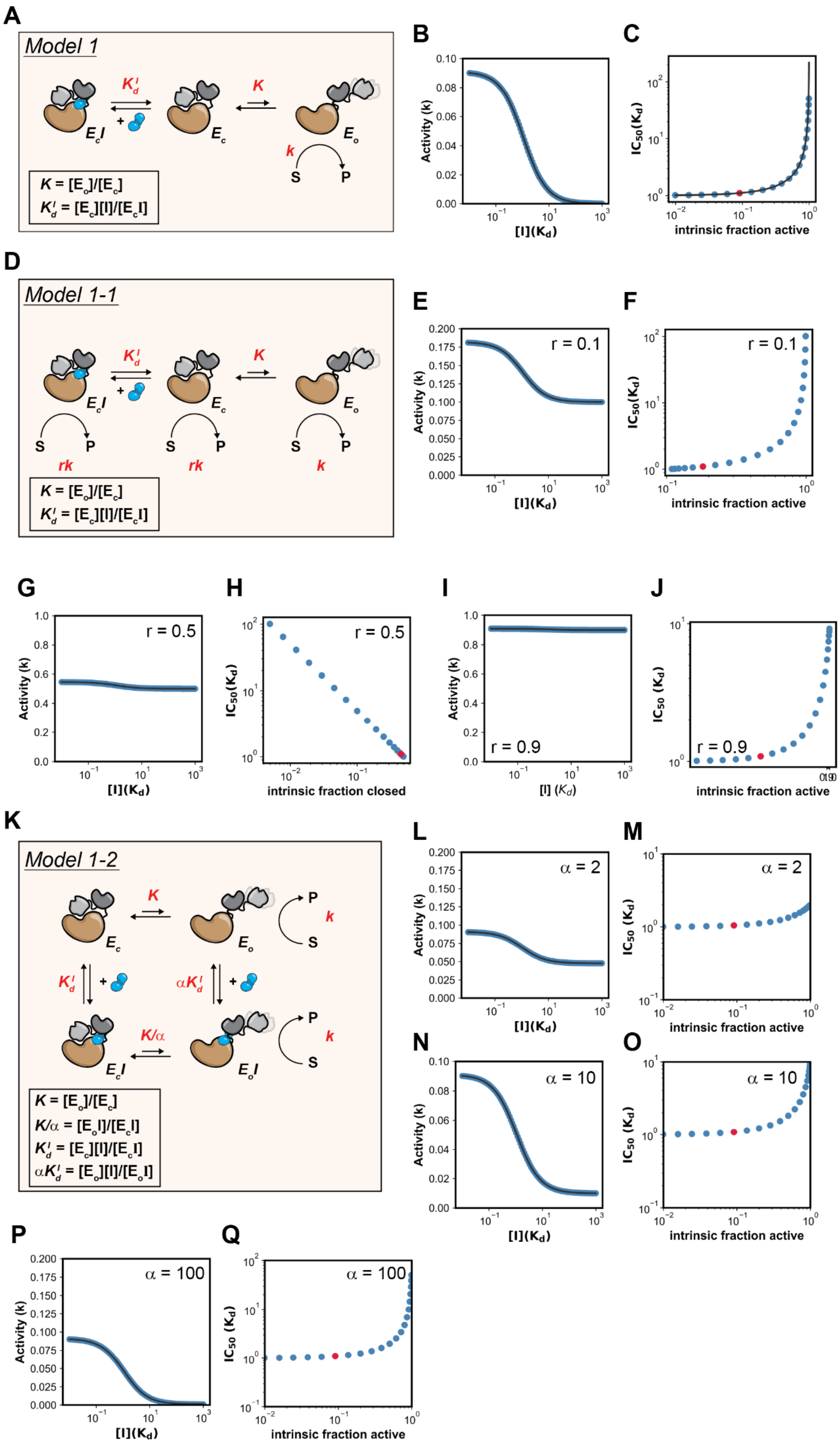
A
Model 1
$K_d^I$
K
$E_cI$
$E_c$
$E_o$
k
S
P
$K = [E_o]/[E_c]$
$K_d^I = [E_c][I]/[E_cI]$
B
Activity (k)
[I](Kd)
C
IC50(Kd)
intrinsic fraction active
D
Model 1-1
rk
k
E
r = 0.1
F
G
r = 0.5
H
intrinsic fraction closed
I
r = 0.9
J
K
Model 1-2
$\alpha K_d^I$
$K/\alpha$
$E_oI$
$K/\alpha = [E_oI]/[E_cI]$
$\alpha K_d^I = [E_o][I]/[E_oI]$
L
α = 2
M
N
α = 10
O
P
α = 100
Q

**Figure S74. Simulation of two conformational state inhibition models. (A)** Schematic of _Model 1_. **(B)** Simulated activity versus inhibitor concentration relationship for WT based on _Model 1_. **(C)** Simulated $IC_{50}$ versus intrinsic fraction active relationship based on _Model 1_. WT is marked in red. **(D)** Schematic of _Model 1-1_. **(E)** Simulated activity versus inhibitor concentration relationship for WT and **(F)** Simulated $IC_{50}$ versus intrinsic fraction active relationship based on _Model 1-1_, with r = 0.1; **(G)** and **(H)** r = 0.5; and **(I)** and **(J)** r =0.9. **(K)** Schematic of _Model 1-2_. **(L)** Simulated activity versus inhibitor concentration relationship for WT and **(M)** Simulated $IC_{50}$ versus intrinsic fraction active relationship based on _Model 1-2_, with α = 2; **(N)** and **(O)** α = 10; and **(P)** and **(Q)** α = 100. WT is marked in red.

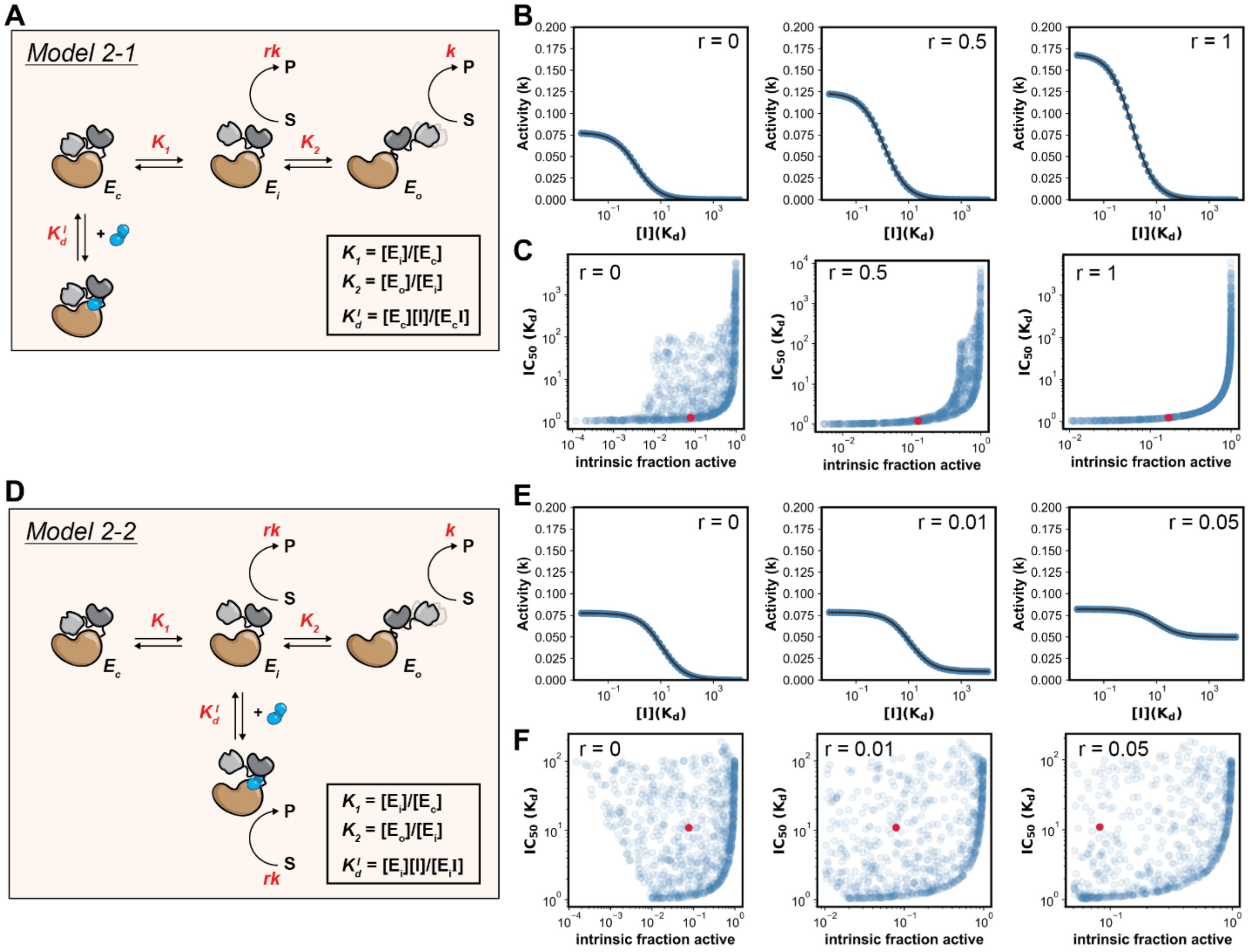


**Figure S75. Simulation of three conformational state inhibition models. (A)** Schematic of _Model 2-1_. **(B)** Simulated activity versus inhibitor concentration relationship for WT based on _Model 2-1_, with r = 0, 0.5, and 1 (left to right). **(C)** Simulated $IC_{50}$ versus intrinsic fraction active relationship based on _Model2-1_, with r = 0, 0.5, and 1 (left to right). WT is marked in red. **(D)** Schematic of _Model 2-2_. **(E)** Simulated activity versus inhibitor concentration relationship for WT based on _Model 2-2_, with r = 0, 0.01, and 0.05 (left to right). **(F)** Simulated $IC_{50}$ versus intrinsic fraction active relationship based on _Model 2-1_, with r = 0, 0.01, and 0.05 (left to right). WT is marked in red.

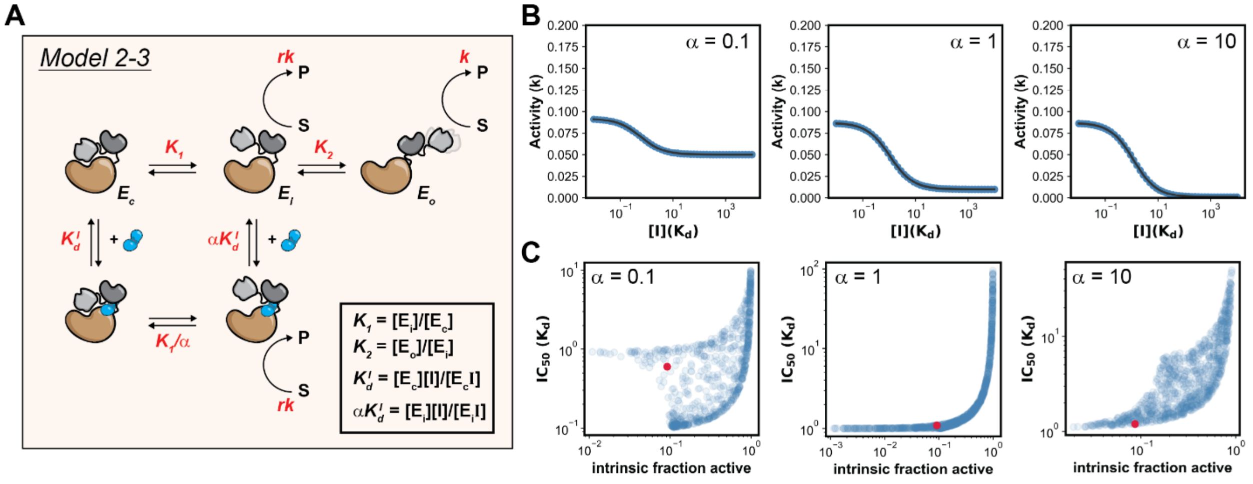


**Figure S76. Simulation of three conformational state inhibition models with inhibitor binding to both closed and the intermediate states. (A)** Schematic of *Model 2-3*. **(B)** Simulated activity versus inhibitor concentration relationship for WT based on *Model 2-3*, with α = 0.1, 1, 10 (left to right). **(C)** Simulated $IC_{50}$ versus intrinsic fraction active relationship based on *Model 2-3*, with α = 0.1, 1, 10 (left to right). WT is marked in red.

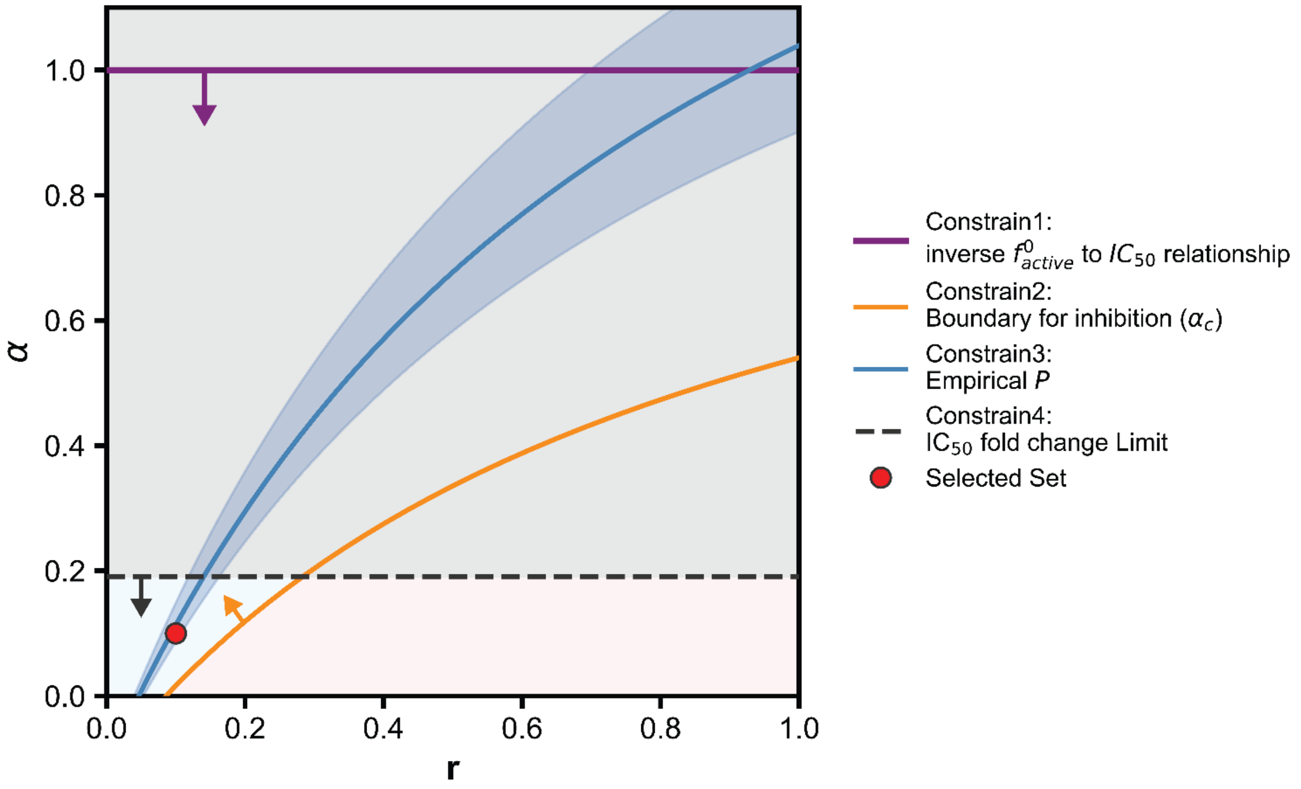


**Figure S77. Space of parameter constraints for simulation.** The relationship between the scaling factor $\alpha$ and parameter $r$ is shown relative to experimental and theoretical limits. The purple line shows the $\alpha < 1$ constrain to show inverse $f^0_{active}$ to $IC_{50}$ relationship. The critical boundary ($\alpha_c$; orange line) separates the activation (light red) and inhibition (light blue) regimes. The steel blue line and shaded band show parameter space that satisfies the plateau ($P$) of experimental inhibition data. The dashed line marks the maximum theoretical threshold for $\alpha$ to produce the experimentally observed $IC_{50}$ fold change from WT and excludes parameter space above it (grey region). The parameter set selected for simulating the three-conformational-state model ($r = 0.1$, $\alpha = 0.1$) is marked in red.

| MutantID | ClinVar_germline_classification | NS/NSML | NS/NSML_counts | Blood_cancer_counts | Solid_cancer_counts | JMML_counts | kNN_prediction |
|---|---|---|---|---|---|---|---|
| *WT* | Benign | — | — | 0 | 0 | 0 | Benign |
| *T2I* | Pathogenic/Likely pathogenic | Noonan syndrome | 9 | 0 | 0 | 0 | Pathogenic |
| *R4G* | Conflicting classifications of pathogenicity | — | — | 0 | 0 | 0 | Pathogenic |
| *N10D* | Uncertain significance | — | — | 0 | 0 | 0 | Benign |
| *N18S* | Benign | — | — | 0 | 0 | 0 | Benign |
| *A31G* | Likely pathogenic | — | — | 0 | 0 | 0 | Benign |
| *K35E* | Uncertain significance | — | — | 0 | 0 | 0 | Benign |
| *K35Q* | Uncertain significance | — | — | 0 | 0 | 0 | Pathogenic |
| *G39R* | Uncertain significance | — | — | 0 | 0 | 0 | Pathogenic |
| *D40G* | Uncertain significance | — | — | 0 | 1 | 0 | Pathogenic |
| *T42A* | Pathogenic | Noonan syndrome | 54 | 3 | 0 | 0 | Pathogenic |
| *L43F* | Uncertain significance | — | — | 0 | 0 | 0 | Pathogenic |
| *V45I* | Uncertain significance | — | — | 0 | 4 | 0 | Benign |
| *A50T* | Uncertain significance | — | — | 1 | 0 | 0 | Benign |
| *T52I* | Likely pathogenic | Noonan syndrome | 3 | 1 | 1 | 0 | Pathogenic |
| *I56T* | Likely pathogenic | — | — | 0 | 0 | 0 | Pathogenic |
| *I56V* | Pathogenic | Noonan syndrome | 2 | 0 | 0 | 0 | Pathogenic |
| *N58S* | Conflicting classifications of pathogenicity | Noonan syndrome | 1 | 1 | 7 | 0 | Pathogenic |
| *N58H* | Pathogenic | Noonan syndrome | 9 | 0 | 0 | 0 | Pathogenic |
| *N58D* | Pathogenic | Noonan syndrome | 34 | 0 | 1 | 0 | Pathogenic |
| *N58K* | Pathogenic | Noonan syndrome | 14 | 2 | 1 | 0 | Pathogenic |
| *N58Y* | Pathogenic/Likely pathogenic | — | — | 13 | 0 | 2 | Pathogenic |
| *T59A* | Conflicting classifications of pathogenicity | Noonan syndrome | 1 | 1 | 0 | 0 | Pathogenic |
| *G60D* | Likely pathogenic | Noonan syndrome | 1 | 0 | 1 | 0 | Pathogenic |
| *G60A* | Pathogenic | Noonan syndrome | 39 | 9 | 0 | 0 | Pathogenic |
| *G60C* | Pathogenic/Likely pathogenic | Noonan syndrome | 2 | 0 | 0 | 0 | Pathogenic |
| *G60V* | Pathogenic/Likely pathogenic | Noonan syndrome | 5 | 59 | 5 | 16 | Pathogenic |
| *G60R* | Pathogenic/Likely pathogenic | — | — | 14 | 1 | 8 | Pathogenic |
| *G60S* | Pathogenic/Likely pathogenic | Noonan syndrome | 3 | 0 | 0 | 0 | Pathogenic |
| *D61Y* | Conflicting classifications of pathogenicity | — | — | 65 | 5 | 33 | Pathogenic |
| *D61H* | Pathogenic | — | — | 11 | 0 | 0 | Pathogenic |
| *D61V* | Pathogenic | — | — | 77 | 3 | 36 | Pathogenic |
| *D61N* | Pathogenic | Noonan syndrome | 82 | 16 | 1 | 1 | Pathogenic |
| *D61G* | Pathogenic | Noonan syndrome | 73 | 10 | 2 | 1 | Pathogenic |
| *D61A* | Pathogenic/Likely pathogenic | Noonan syndrome | 3 | 5 | 1 | 0 | Pathogenic |
| *Y62N* | Likely pathogenic | Noonan syndrome | 1 | 0 | 0 | 0 | Pathogenic |
| *Y62D* | Pathogenic | Noonan syndrome | 57 | 1 | 0 | 0 | Pathogenic |
| *Y63C* | Pathogenic | Noonan syndrome | 160 | 6 | 0 | 0 | Pathogenic |
| *E69K* | Conflicting classifications of pathogenicity | — | — | 53 | 25 | 6 | Pathogenic |
| *E69Q* | Pathogenic | Noonan syndrome | 13 | 0 | 1 | 0 | Pathogenic |
| *E69V* | Pathogenic/Likely pathogenic | Noonan syndrome | 2 | 5 | 0 | 0 | Pathogenic |
| *K70R* | Pathogenic | — | — | 2 | 0 | 0 | Pathogenic |
| *F71V* | Pathogenic | — | — | 0 | 0 | 0 | Pathogenic |
| *F71L* | Pathogenic/Likely pathogenic | Noonan syndrome | 7 | 26 | 4 | 0 | Pathogenic |
| *A72V* | Conflicting classifications of pathogenicity | Noonan syndrome | 3 | 98 | 16 | 31 | Pathogenic |
| *A72S* | Pathogenic | Noonan syndrome | 45 | 2 | 2 | 0 | Pathogenic |
| *A72P* | Pathogenic/Likely pathogenic | Noonan syndrome | 3 | 3 | 0 | 1 | Pathogenic |
| *A72G* | Pathogenic/Likely pathogenic | Noonan syndrome | 37 | 2 | 2 | 0 | Pathogenic |
| *A72T* | Pathogenic/Likely pathogenic | — | — | 74 | 12 | 23 | Pathogenic |
| *A72D* | Likely pathogenic | Noonan syndrome | 1 | 16 | 6 | 1 | Pathogenic |
| *T73I* | Pathogenic | Noonan syndrome | 47 | 30 | 2 | 5 | Pathogenic |
| *T73P* | Pathogenic/Likely pathogenic | — | — | 0 | 0 | 0 | Pathogenic |
| *A75P* | Uncertain significance | — | — | 0 | 0 | 0 | Pathogenic |
| *E76K* | Conflicting classifications of pathogenicity | — | — | 171 | 17 | 89 | Pathogenic |
| *E76G* | Conflicting classifications of pathogenicity | Noonan syndrome | 1 | 69 | 4 | 38 | Pathogenic |
| *E76V* | Pathogenic | Noonan syndrome | 1 | 24 | 1 | 7 | Pathogenic |
| *E76D* | Pathogenic | Noonan syndrome | 18 | 1 | 0 | 0 | Pathogenic |
| *E76A* | Pathogenic | — | — | 29 | 6 | 10 | Pathogenic |
| *E76Q* | Pathogenic/Likely pathogenic | Noonan syndrome | 1 | 28 | 3 | 8 | Pathogenic |
| *Q79K* | Likely pathogenic | — | — | 0 | 0 | 0 | Pathogenic |
| *Q79R* | Pathogenic | Noonan syndrome | 132 | 8 | 0 | 0 | Pathogenic |
| *M82V* | Uncertain significance | — | — | 0 | 1 | 0 | Benign |
| *K91R* | Uncertain significance | — | — | 0 | 0 | 0 | Benign |
| *E97Q* | Uncertain significance | — | — | 0 | 0 | 0 | Benign |
| *D106A* | Pathogenic | Noonan syndrome | 31 | 1 | 0 | 0 | Pathogenic |
| *E110D* | Likely pathogenic | — | — | 0 | 0 | 0 | Pathogenic |
| *E110A* | Pathogenic/Likely pathogenic | — | — | 0 | 0 | 0 | Pathogenic |
| *E110K* | Pathogenic/Likely pathogenic | Noonan syndrome | 4 | 0 | 0 | 0 | Pathogenic |
| *H116Q* | Conflicting classifications of pathogenicity | — | — | 0 | 1 | 0 | Pathogenic |
| *H116N* | Uncertain significance | — | — | 0 | 0 | 0 | Pathogenic |
| *E123D* | Uncertain significance | — | — | 0 | 5 | 0 | Benign |
| *K131R* | Likely benign | — | — | 0 | 0 | 0 | Benign |
| *E139D* | Pathogenic | Noonan syndrome | 79 | 5 | 2 | 3 | Pathogenic |
| *S140C* | Uncertain significance | — | — | 0 | 0 | 0 | Pathogenic |
| *F147C* | Uncertain significance | — | — | 0 | 0 | 0 | Benign |
| *R152H* | Uncertain significance | Noonan syndrome | 1 | 1 | 1 | 0 | Benign |
| *G154A* | Uncertain significance | — | — | 0 | 0 | 0 | Benign |
| *K157R* | Uncertain significance | — | — | 0 | 0 | 0 | Benign |
| *G158A* | Uncertain significance | — | — | 0 | 0 | 0 | Benign |

| | | | | | | | |
|---|---|---|---|---|---|---|---|
| *G163S* | Uncertain significance | — | — | 0 | 1 | 0 | Benign |
| *T168A* | Conflicting classifications of pathogenicity | — | — | 0 | 0 | 0 | Benign |
| *R173P* | Likely pathogenic | — | — | 0 | 0 | 0 | Benign |
| *R173H* | Uncertain significance | — | — | 0 | 1 | 0 | Benign |
| *R186W* | Conflicting classifications of pathogenicity | Noonan syndrome | 1 | 0 | 0 | 0 | Benign |
| *D188H* | Uncertain significance | — | — | 0 | 0 | 0 | Pathogenic |
| *D188N* | Uncertain significance | — | — | 0 | 0 | 0 | Pathogenic |
| *N200Y* | Pathogenic/Likely pathogenic | Noonan syndrome with multiple lentigines | 5 | 0 | 0 | 0 | Benign |
| *T218S* | Uncertain significance | — | — | 0 | 0 | 0 | Pathogenic |
| *T218A* | Uncertain significance | — | — | 0 | 0 | 0 | Pathogenic |
| *I221V* | Conflicting classifications of pathogenicity | Noonan syndrome | 1 | 0 | 0 | 0 | Pathogenic |
| *I221L* | Uncertain significance | — | — | 0 | 0 | 0 | Pathogenic |
| *S228I* | Uncertain significance | — | — | 0 | 0 | 0 | — |
| *Q245E* | Uncertain significance | — | — | 0 | 0 | 0 | Benign |
| *Q256K* | Pathogenic/Likely pathogenic | Noonan syndrome | 1 | 0 | 0 | 0 | Pathogenic |
| *E258K* | Conflicting classifications of pathogenicity | — | — | 0 | 0 | 0 | Pathogenic |
| *E258D* | Pathogenic/Likely pathogenic | Noonan syndrome | 1 | 0 | 1 | 0 | Pathogenic |
| *L262R* | Pathogenic | Noonan syndrome | 6 | 0 | 0 | 0 | Pathogenic |
| *Y263C* | Uncertain significance | — | — | 0 | 0 | 0 | Pathogenic |
| *R265L* | Likely pathogenic | — | — | 0 | 0 | 0 | Pathogenic |
| *R265Q* | Pathogenic | Noonan syndrome | 10 | 4 | 2 | 0 | Pathogenic |
| *G268S* | Pathogenic/Likely pathogenic | Noonan syndrome | 7 | 1 | 0 | 0 | Pathogenic |
| *G268C* | Pathogenic/Likely pathogenic | Noonan syndrome | 2 | 0 | 0 | 0 | Pathogenic |
| *Q269R* | Uncertain significance | — | — | 0 | 0 | 0 | Benign |
| *N275T* | Conflicting classifications of pathogenicity | — | — | 0 | 0 | 0 | Benign |
| *K276R* | Uncertain significance | — | — | 0 | 0 | 0 | Benign |
| *Y279S* | Pathogenic | Noonan syndrome with multiple lentigines | 10 | 0 | 0 | 0 | Pathogenic |
| *Y279C* | Pathogenic/Likely pathogenic | Noonan syndrome with multiple lentigines | 112 | 0 | 3 | 0 | Pathogenic |
| *I282T* | Pathogenic | — | — | 0 | 0 | 0 | Pathogenic |
| *I282V* | Pathogenic | Noonan syndrome | 35 | 0 | 1 | 0 | Pathogenic |
| *I282M* | Pathogenic/Likely pathogenic | Noonan syndrome | 3 | 0 | 3 | 0 | Pathogenic |
| *F285V* | Conflicting classifications of pathogenicity | — | — | 0 | 0 | 0 | Pathogenic |
| *F285S* | Pathogenic | Noonan syndrome | 57 | 10 | 6 | 0 | Pathogenic |
| *F285I* | Pathogenic | Noonan syndrome | 5 | 0 | 0 | 0 | Pathogenic |
| *F285L* | Pathogenic | Noonan syndrome | 24 | 1 | 0 | 0 | Pathogenic |
| *F285Y* | Pathogenic/Likely pathogenic | — | — | 0 | 0 | 0 | Pathogenic |
| *F285C* | Pathogenic/Likely pathogenic | Noonan syndrome | 1 | 0 | 2 | 0 | Pathogenic |
| *D286E* | Uncertain significance | — | — | 0 | 0 | 0 | — |
| *V290I* | Uncertain significance | — | — | 0 | 0 | 0 | — |
| *H293Q* | Uncertain significance | — | — | 0 | 0 | 0 | Benign |
| *N298S* | Conflicting classifications of pathogenicity | — | — | 1 | 0 | 0 | Benign |
| *I305V* | Uncertain significance | — | — | 0 | 0 | 0 | Benign |
| *N308D* | Pathogenic | Noonan syndrome | 402 | 6 | 5 | 0 | Pathogenic |
| *N308S* | Pathogenic | Noonan syndrome | 125 | 2 | 2 | 0 | Pathogenic |
| *N308T* | Pathogenic/Likely pathogenic | Noonan syndrome | 6 | 1 | 2 | 0 | Pathogenic |
| *I309V* | Benign | — | — | 1 | 3 | 0 | Benign |
| *M311T* | Likely benign | — | — | 0 | 0 | 0 | Benign |
| *M311V* | Uncertain significance | — | — | 2 | 0 | 0 | Benign |
| *F314L* | Uncertain significance | — | — | 0 | 0 | 0 | Benign |
| *N320S* | Likely pathogenic | — | — | 0 | 0 | 0 | Benign |
| *K324R* | Uncertain significance | — | — | 0 | 0 | 0 | Benign |
| *C333S* | Uncertain significance | — | — | 0 | 0 | 0 | Benign |
| *Q335P* | Uncertain significance | — | — | 0 | 0 | 0 | Benign |
| *R343Q* | Benign | — | — | 0 | 1 | 0 | Benign |
| *R343W* | Uncertain significance | — | — | 0 | 1 | 0 | Benign |
| *Q347R* | Uncertain significance | — | — | 0 | 0 | 0 | Benign |
| *R351Q* | Benign | Noonan syndrome | 2 | 1 | 5 | 0 | Benign |
| *K364R* | Uncertain significance | — | — | 0 | 0 | 0 | Benign |
| *S365R* | Uncertain significance | — | — | 0 | 0 | 0 | Benign |
| *Y375C* | Uncertain significance | — | — | 0 | 0 | 0 | Benign |
| *V382I* | Uncertain significance | — | — | 0 | 2 | 0 | Benign |
| *H394R* | Uncertain significance | — | — | 0 | 0 | 0 | — |
| *T397M* | Uncertain significance | — | — | 0 | 2 | 0 | Benign |
| *V406I* | Uncertain significance | — | — | 0 | 0 | 0 | Benign |
| *G409A* | Uncertain significance | — | — | 0 | 2 | 0 | Benign |
| *T411M* | Uncertain significance | Noonan syndrome | 4 | 0 | 1 | 0 | Benign |
| *R421W* | Uncertain significance | — | — | 0 | 0 | 0 | Benign |
| *P424L* | Likely pathogenic | — | — | 0 | 1 | 0 | Pathogenic |
| *D425N* | — | — | — | 0 | 2 | 0 | Pathogenic |
| *H426P* | Uncertain significance | — | — | 0 | 0 | 0 | Pathogenic |
| *V428M* | Conflicting classifications of pathogenicity | Noonan syndrome | 1 | 4 | 10 | 0 | Pathogenic |
| *V428L* | Pathogenic/Likely pathogenic | — | — | 0 | 0 | 0 | Pathogenic |
| *H443Y* | Uncertain significance | — | — | 0 | 1 | 0 | Benign |
| *P454L* | Uncertain significance | — | — | 0 | 1 | 0 | Benign |
| *V456M* | Uncertain significance | — | — | 0 | 0 | 0 | Benign |
| *H458Q* | Uncertain significance | — | — | 0 | 0 | 0 | Pathogenic |
| *A461G* | Conflicting classifications of pathogenicity | — | — | 5 | 2 | 0 | Pathogenic |
| *A461S* | Pathogenic | Noonan syndrome with multiple lentigines | 2 | 0 | 1 | 0 | — |

| | | | | | | | |
|---|---|---|---|---|---|---|---|
| ***T468E*** | Conflicting classifications of pathogenicity | — | — | 0 | 0 | 0 | Pathogenic |
| ***T468A*** | Conflicting classifications of pathogenicity | — | — | 0 | 0 | 0 | Pathogenic |
| ***T468S*** | Likely benign | — | — | 0 | 0 | 0 | Pathogenic |
| ***T468M*** | Pathogenic | Noonan syndrome with multiple lentigines | 135 | 6 | 14 | 3 | Pathogenic |
| ***T468P*** | Pathogenic | Noonan syndrome with multiple lentigines | 3 | 0 | 0 | 0 | Pathogenic |
| ***I470T*** | Likely pathogenic | — | — | 0 | 0 | 0 | Benign |
| ***I479T*** | Uncertain significance | — | — | 0 | 0 | 0 | Benign |
| ***D487G*** | Uncertain significance | — | — | 0 | 0 | 0 | Pathogenic |
| ***V490I*** | Uncertain significance | — | — | 0 | 1 | 0 | Benign |
| ***P491S*** | Pathogenic | Noonan syndrome | 23 | 4 | 2 | 0 | Pathogenic |
| ***P491A*** | Pathogenic/Likely pathogenic | Noonan syndrome | 2 | 0 | 0 | 0 | Pathogenic |
| ***P491H*** | Pathogenic/Likely pathogenic | Noonan syndrome | 7 | 2 | 0 | 0 | Pathogenic |
| ***P491T*** | Pathogenic/Likely pathogenic | Noonan syndrome | 1 | 0 | 0 | 0 | Pathogenic |
| ***R498W*** | Pathogenic | Noonan syndrome with multiple lentigines | 12 | 1 | 14 | 0 | Pathogenic |
| ***R498L*** | Pathogenic | Noonan syndrome with multiple lentigines | 5 | 0 | 2 | 0 | Pathogenic |
| ***S499T*** | Uncertain significance | — | — | 0 | 0 | 0 | Pathogenic |
| ***S499F*** | Uncertain significance | Noonan syndrome with multiple lentigines | 1 | 0 | 1 | 0 | — |
| ***Q500H*** | Uncertain significance | — | — | 0 | 0 | 0 | Pathogenic |
| ***R501K*** | Pathogenic/Likely pathogenic | Noonan syndrome | 4 | 1 | 0 | 0 | Pathogenic |
| ***S502T*** | Pathogenic | Noonan syndrome | 16 | 2 | 1 | 1 | Pathogenic |
| ***S502L*** | Pathogenic | Noonan syndrome | 12 | 24 | 9 | 0 | Pathogenic |
| ***S502P*** | — | Noonan syndrome | 1 | 21 | 5 | 1 | Pathogenic |
| ***G503V*** | other | — | — | 29 | 30 | 9 | Pathogenic |
| ***G503R*** | Pathogenic | Noonan syndrome | 21 | 17 | 3 | 1 | — |
| ***G503A*** | Pathogenic | Noonan syndrome | 1 | 46 | 1 | 21 | Pathogenic |
| ***G503E*** | Pathogenic/Likely pathogenic | Noonan syndrome | 7 | 11 | 0 | 0 | Pathogenic |
| ***M504V*** | Pathogenic | Noonan syndrome | 128 | 1 | 0 | 0 | Pathogenic |
| ***Q506P*** | Pathogenic | Noonan syndrome with multiple lentigines | 17 | 6 | 2 | 0 | Pathogenic |
| ***T507K*** | Pathogenic/Likely pathogenic | Noonan syndrome | 2 | 9 | 14 | 0 | Pathogenic |
| ***Q510L*** | Likely pathogenic | Noonan syndrome | 1 | 5 | 5 | 0 | Pathogenic |
| ***Q510E*** | Pathogenic | Noonan syndrome with multiple lentigines | 40 | 2 | 1 | 0 | Pathogenic |
| ***M516T*** | Uncertain significance | — | — | 0 | 0 | 0 | Pathogenic |
| ***E523D*** | Uncertain significance | — | — | 0 | 0 | 0 | Benign |

**Supplementary Table 1. Clinical annotations and biochemical pathogenicity predictions for 190 SHP2 variants in this study.** Clinical classifications and disease associations for each variant, alongside *k*-nearest neighbor (*k*NN) predictions of biochemical pathogenicity (**Fig. 4B**). Columns: **MutantID**, amino acid substitution; **ClinVar_germline_classification**, germline classification from ClinVar; **NS/NSML**, disease subtype annotation from the NSEuroNet database (www.nseuronet.com). Variants with both NS and NSML annotations were classified as NSML; **NS/NSML_counts**, number of cases reported in NSEuroNet; **Blood_cancer_counts**, **Solid_cancer_counts**, and **JMML_counts**, number of cases in the COSMIC database; **kNN_prediction**, predicted pathogenicity (Benign or Pathogenic) from *k*NN classification using biochemical parameters. The table is also available as a separate CSV file.

**Captions for Supplementary Data**

**Supplementary Data 1.** SHP2 kinetic and thermodynamic parameter table, CSV (AuxiliarySupplementaryMaterials_1.csv)

**Supplementary Data 2.** SHP2 TNO155 $IC_{50}$ table, CSV (AuxiliarySupplementaryMaterials_2.csv)

**Supplementary Data 3.** SHP2 RMC-4630 $IC_{50}$ table, CSV (AuxiliarySupplementaryMaterials_3.csv)

**Supplementary Data 4.** SHP2 GDC-1971 $IC_{50}$ table, CSV (AuxiliarySupplementaryMaterials_4.csv)
